# Entry Steps in the Biosynthetic Pathway to Diterpenoid Alkaloids

**DOI:** 10.1101/2025.05.15.654307

**Authors:** Garret P. Miller, Lana Mutabdžija-Nedelcheva, Trine B. Andersen, Imani Pascoe, Kathryn Van Winkle, Maryam Sabbaghan, Alexandre Bouillé, Tomáš Iliaš, Andrej Tekel, Tomáš Pluskal, Björn Hamberger

## Abstract

Both the terpenoid and alkaloid classes of specialized metabolites have received considerable attention for their wide range of practical applications, however, few examples have been identified of these classes intersecting. The diterpenoid alkaloids are one such example of nitrogen-containing terpenoids which are found throughout many independent plant lineages, but primarily within the *Aconitum* (Wolf’s-Bane) and *Delphinium* (Larkspur) genera. While there is considerable interest in these compounds for their wide range of bioactivities, their structural complexity often precludes their production through chemical synthesis, and little progress has been made towards elucidation of their biosynthetic pathways. Here, we employ a comparative transcriptomics approach to identify six enzymatic steps in the biosynthesis of atisinium, conserved across both *Delphinium grandiflorum* and *Aconitum plicatum*. Key to this pathway is a reductase which selectively incorporates ethanolamine over ethylamine into the diterpenoid scaffold. While the majority of diterpenoid alkaloids contain an ethylamine moiety, we demonstrate through isotope labeling in *Aconitum* callus cultures and a computational metabolomics approach that ethanolamine is, unintuitively, the preferred source of nitrogen for these metabolites. Identification of these enzymes and production of a key intermediate in a heterologous host paves the way for biosynthetic production of this group of metabolites with promise for medicinal applications.

## Introduction

To support their defense, interactions with other organisms, and ecological adaptation, land plants have evolved a plethora of structurally diverse specialized metabolites. Such metabolites often feature biologically relevant molecular scaffolds which have proven paramount for the discovery of chemical probes and drugs^1^. Among these, both the alkaloid and terpenoid classes have received considerable attention for their medicinal applications, with prominent examples including taxol^2^ (anti-cancer), artemisinin^3^ (antiplasmodial), morphine^4^ (analgesic), colchicine^5^ (anti-inflammatory), scopolamine^6–8^ (anti-nausea), and vinblastine^9–11^ (anti-cancer). Their importance in the pharmaceutical industry has led to a wealth of research into elucidation of their biosynthetic pathways and production in heterologous hosts, as direct extraction from plants or complete chemical synthesis is often challenging or unfeasible.

Given the unique pathways towards initial scaffold formation, there is little overlap between the terpenoid and alkaloid classes of specialized metabolites. One notable exception is the diterpenoid alkaloids, which are found throughout many independent plant lineages^12–14^, but primarily within the cosmopolitan Ranunculaceae family^15,16^ (Supplementary Figure 1). The biosynthesis of this class of metabolites has not been elucidated, however it is apparent from their structure that it involves the initial formation of a diterpene scaffold followed by nitrogen incorporation, and extensive modifications by various oxygenases, acyltransferases, and methyltransferases.

Plants from the *Aconitum* and *Delphinium* genera have long been used in traditional medicine due to the bioactivity of diterpenoid alkaloids, which primarily accumulate in their root tissue^17–20^. The use of “Fuzi”— the processed lateral root of *A. carmichaelii* (more commonly known as Wolf’s Bane or Aconite)—has been documented for at least two thousand years^16^. The diterpenoid alkaloids have a wide range of potential applications from antifeedants to anti-cancer, antiplasmodial, cholinesterase inhibitors, and analgesics^15,16,21–23^ with lappaconitine, 3-acetylaconitine and crassicauline A used as non-narcotic analgesic drugs^24^. The therapeutic properties of many of these metabolites have prompted an extensive amount of research into their total chemical synthesis^25–29^, however, their structural complexity presents an enormous challenge in chemical synthesis. This is best exemplified by aconitine (Figure 1), a potent neurotoxin with six interconnected rings and fifteen stereocenters. Despite being first isolated in 1833 by P. L. Geiger and numerous attempts to synthesize it chemically, aconitine has not yet been successfully synthesized by chemists^30^.

**Figure 1:**
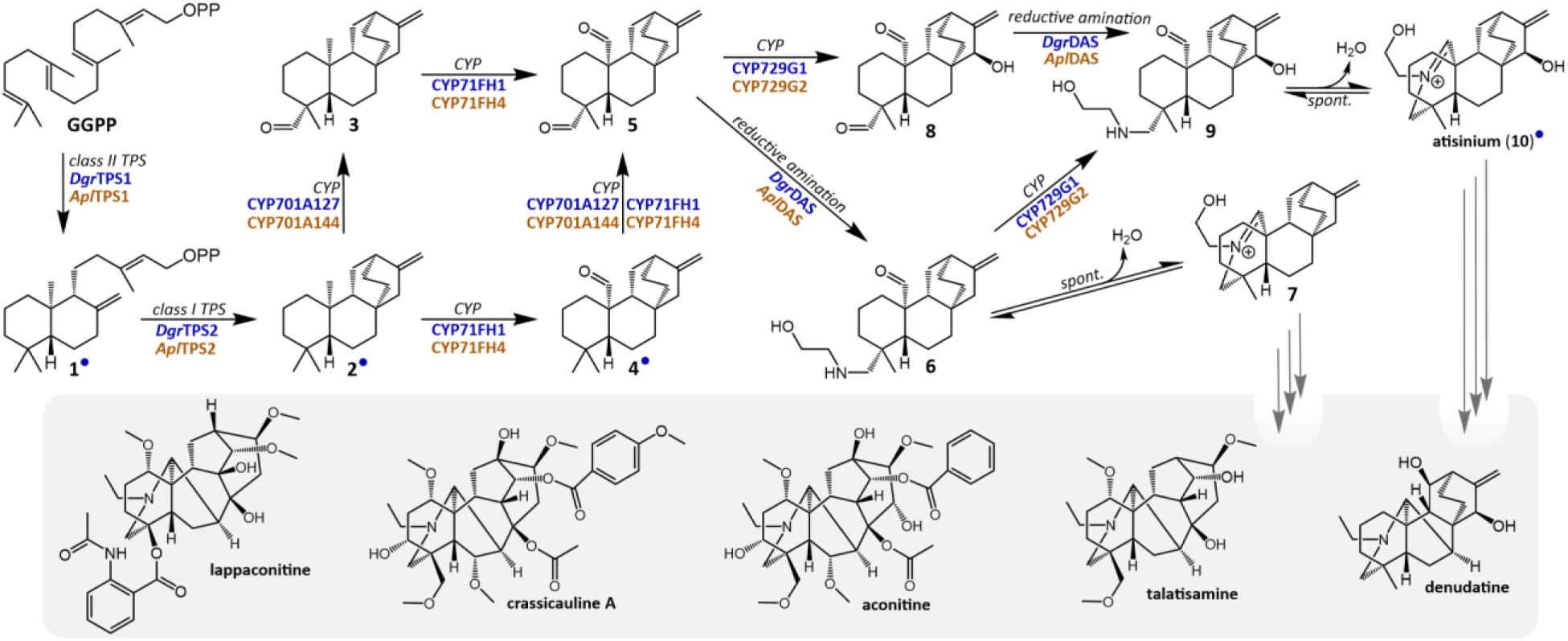
Proposed biosynthetic pathway towards diterpenoid alkaloids. Putative enzyme classes or chemical transformations used to guide our approach are annotated along each arrow, with enzymes identified in this study for each step from *D. grandiflorum* and *A. plicatum* written in blue and orange, respectively. The transformations labeled “reductive amination” likely proceed through spontaneous condensation with ethanolamine followed by reduction by DAS, as described in the text. Compound numbers with blue dots are those supported by NMR or comparison to standards. Rationale for structural assignments and predictions are expanded upon in Supplementary Table 3. Example diterpenoid alkaloid structures are shown below in gray.

Elucidating the biosynthesis of these compounds would ameliorate some of the challenges in their production given the complexity of their scaffolds and the number of required stereospecific oxidations. The lack of current knowledge in their biosynthesis is not for a lack of effort, as there is a large body of work involving transcriptomic analysis of these pathways in various *Aconitum* species^31–36^. Only recently our understanding was further expanded beyond the initial steps involving terpene synthases^36,37^ to include cytochrome P450-mediated oxidations^38^. However, the enzyme(s) responsible for nitrogen incorporation, a defining feature of diterpenoid alkaloids, remain unidentified. To our knowledge, similar work has not yet been done on the neighboring *Delphinium* genus, despite the likely conservation of this pathway between both genera.

To address the gaps in the biosynthetic pathway of diterpenoid alkaloids, we carried out transcriptome sequencing on *Delphinium grandiflorum*, *Aconitum plicatum* and *Aconitum lycoctonum*, and also included public data from four other *Aconitum* species. Transcriptome assembly both for *D. grandiflorum* and for six representatives of the *Aconitum* genus (*A. plicatum*, *A. lycoctonum*, *A. carmichaelii*, *A. japonicum*, *A. kusnezoffii*, and *A. vilmorinianum*)—all of which accumulate diterpenoid alkaloids—allowed for comparative transcriptomics across tissue types and genera, leading to the identification of six enzymes active in this pathway. Furthermore, the public data for *A. vilmorinianum*—a root tissue timecourse study^31^—allowed for coexpression analysis, resulting in the identification of a novel reductase active in the pathway which has little homology to previously characterized enzymes. This reductase catalyzes a key step in the pathway, supporting the formation of atisinium (**10**, Figure 1)—a bioactive diterpenoid alkaloid and potential intermediate in the biosynthesis of more complex diterpenoid alkaloids^23^. Despite the abundance of diterpenoid alkaloids with ethylamine groups attached to their central terpene scaffolds, we further demonstrate that ethanolamine is the preferred substrate for this reductase and the primary nitrogen source for the majority of detected diterpenoid alkaloids in *Aconitum* plants. Identification of these entry steps allowed for reconstruction of a minimal pathway sufficient for *de novo* biosynthesis of atisinium (**10**), and will serve as the basis for further pathway discovery towards more complex diterpenoid alkaloid natural products and their biosynthetic production in heterologous hosts.

## Results

### A Pair of TPSs Cyclize GGPP to *ent*-atiserene

The majority of diterpenoid alkaloids in the Ranunculaceae family can be divided into three groups based on the number of carbons in their backbone structure (C_18_, C_19_, and C_20_), each of which is further divided into multiple subgroups represented by 46 distinct scaffolds^15,24^. Despite considerable structural differences between them which reflect variations in their biosynthetic pathways, they appear to share the same initial biosynthesis with the majority of scaffolds believed to arise primarily from *ent*-atiserene (**2**)^24,39^. Our initially-proposed biosynthetic pathway is detailed in Figure 1. Initially, the cyclization pattern of geranylgeranyl diphosphate (GGPP) follows a class II terpene synthase (TPS) mechanism, with three distinct stereocenters which suggest the involvement of an *ent*-copalyl diphosphate (**1**, *ent*-CPP) synthase. The ring pattern of the majority of diterpenoid alkaloids is consistent with that of *ent*-atiserene (**2**), suggesting the involvement of a class I TPS to convert *ent*-CPP (**1**) to this scaffold. Subsequent nitrogen incorporation would first require oxidative functionalization of key methyl groups on the *ent*-atiserene (**2**) scaffold, likely to aldehydes (**3**-**5**), suggesting the involvement of one or more cytochromes P450 (CYPs). The final incorporation of this nitrogen group likely involves a reductive amination and marks the transition from diterpenoids to diterpenoid alkaloids^24^.

In order to identify candidate enzymes for this pathway, we opted to cross-reference transcriptomic data across multiple species and tissue types. We isolated and sequenced RNA from several tissues of *Aconitum plicatum* (root, flower, leaf, and stem), *Aconitum lycoctonum* (root, flower, leaf, stem, and fruit), and *Delphinium grandiflorum* (root, leaf, and flower), and carried out *de novo* transcriptome assembly for each. Furthermore, datasets from *A. carmichaelii* (root, leaf, flower, bud; PRJNA415989)^33^, *A. japonicum* (root, root tuber, leaf, flower, stem; PRJDB4889), *A. kusnezoffii* (leaf, principal root, and lateral root; PRJNA670255)^38^ and *A. vilmorinianum* (root time course; PRJNA667080)^31^ from the NCBI Sequence Read Archive (SRA) were included as well.

A BLAST search of the *D. grandiflorum* transcriptome against a reference set of plant TPSs (Supplementary Dataset 1) revealed fourteen putative TPS genes between the two subfamilies typically implicated in diterpene biosynthesis: class II TPSs from the TPS-c subfamily, and class I TPSs from the TPS-e subfamily. Only three of these were exclusively expressed in root tissue, matching the tissue-specific accumulation of diterpenoid alkaloids. *Dgr*TPS1 (TPS-c) and *Dgr*TPS2 (TPS-e) appeared to be the most likely candidates, as they belong to the pair of subfamilies typically implicated in labdane-related diterpene biosynthesis.

Full-length genes for *Dgr*TPS1 and *Dgr*TPS2 were cloned from *D. grandiflorum* root cDNA for *Agrobacterium-*mediated transient expression in *Nicotiana benthamiana*. Two isoforms of *Dgr*TPS2, named *Dgr*TPS2a and *Dgr*TPS2b, were cloned from cDNA and both were tested. All screening through transient expression in *N. benthamiana* included coexpression with *Cf*DXS and *Cf*GGPPS (to increase precursor supply of GGPP^40^). GC-MS analysis of hexane extracts revealed that *Dgr*TPS1 acts as a copalyl diphosphate (CPP) synthase (Supplementary Figure 3A). Coexpression of an enantioselective *ent*-kaurene synthase (*Nm*TPS2)^41^ led to production of *ent*-kaurene, suggesting an absolute stereochemistry for the product of *Drg*TPS1 consistent with *ent*-CPP (**1**) (Supplementary Figure 3A). Furthermore, *Dgr*TPS2a and 2b showed conversion of **1** (Figure 2B) to a new product with a fragmentation pattern (Figure 2C) similar to that of *ent*-atiserene (**2**)^42^ for both isoforms. To confirm the identity of this new product as **2**, transient expression in *N. benthamiana* was scaled up with *Dgr*TPS1 and *Dgr*TPS2a, and the product was purified through silica chromatography and confirmed through NMR (Supplementary Table 1 and Supplementary Figure 4). Since both isoforms of *Dgr*TPS2 exhibit the same function, *Dgr*TPS2a was used for further testing and is referred to as *Dgr*TPS2 hereafter. Cloning and characterization of the respective orthologs from *Aconitum plicatum* revealed that both *Apl*TPS1 and *Apl*TPS2 have this same activity and enantioselectivity, and demonstrates the conservation of the entry steps of this pathway across both genera (Supplementary Figure 3C).

**Figure 2:**
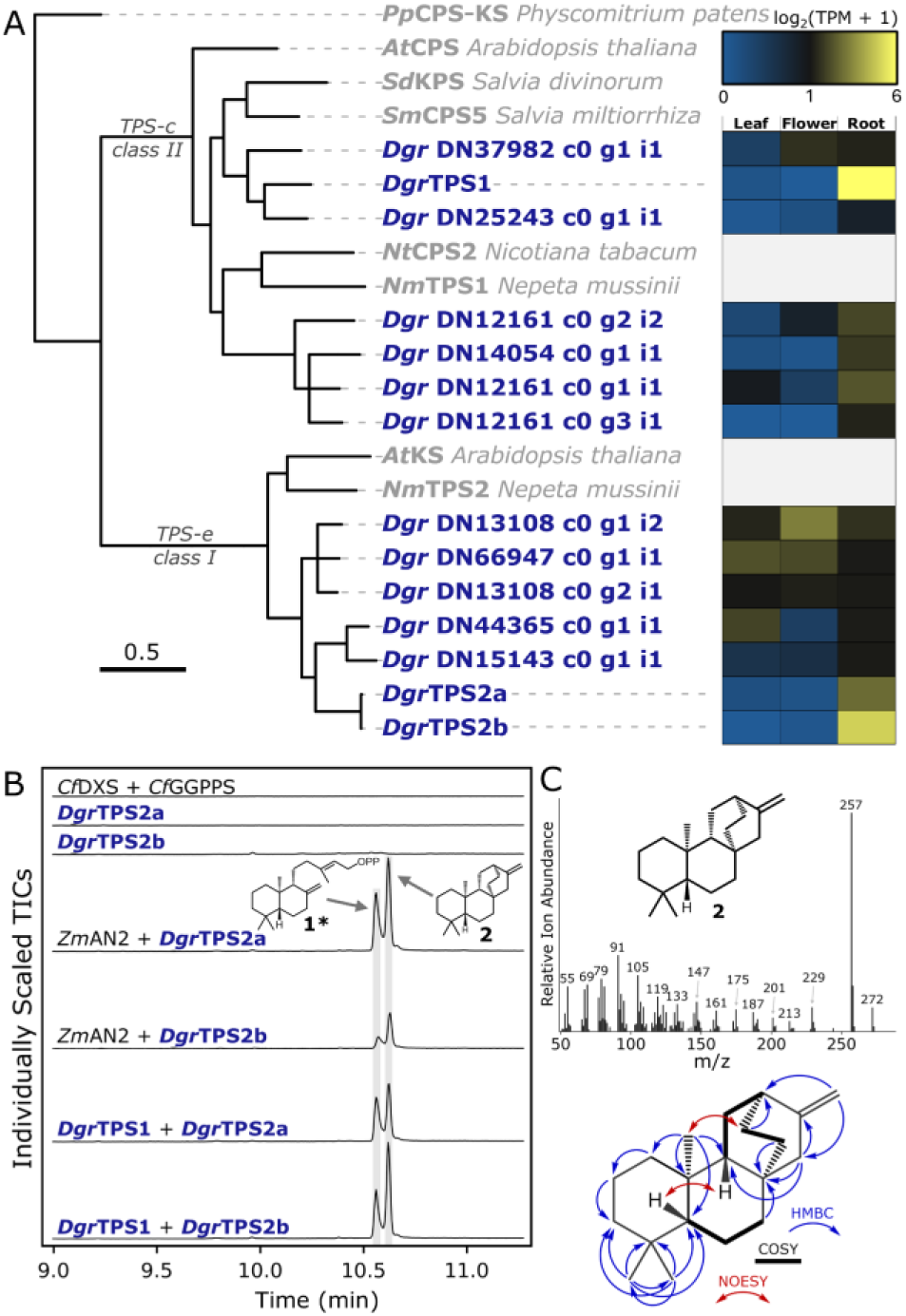
Identification of a pair of terpene synthases which produce ent-atiserene. **A)** Maximum likelihood phylogenetic tree of predicted *D. grandiflorum* TPS-c and TPC-e sequences and their respective expression across tissue types. Reference TPS sequences are in gray with *Pp*CPS-KS (*Physcomitrium patens*) used as an outgroup, scale bar represents substitutions per site, and branches with less than 50% bootstrap support have been collapsed. Expanded view of this tree with *Aconitum* sequences is given in Supplementary Figure 2. **B**) GC-MS analysis of *Dgr*TPS1 and *Dgr*TPS2 coexpression in *N. benthamiana*. *Zm*AN2 (*Zea mays*) serves as a reference *ent*-CPP (**1**) synthase. Marking of **1**\* indicates the dephosphorylated derivative of **1**, as **1** itself is not detectable by GC-MS. Each assay has *Cf*DXS and *Cf*GGPPS (*Coleus forskohlii*) coexpressed in addition to those listed. **C**) Mass spectrum (70 eV EI) of **2** from panel B, and select HMBC, NOESY, and COSY correlations for *ent*-atiserene (**2**) purified from *N. benthamiana* expressing *Dgr*TPS1 and *Dgr*TPS2a.

### Three Cytochromes P450 Oxidize the *ent*-atiserene Scaffold

Following the confirmation that a pair of terpene synthases produce **2**, we continued with our proposed biosynthetic pathway to search for cytochromes P450 (CYPs) which can carry out oxidations of methyl groups 19 and 20 to aldehydes. The proposed intermediate *ent*-atiserene-19-al (**3**) resembles the central metabolite *ent*-kaurenoic acid—a key compound in the primary metabolic pathway towards gibberellins^43^—which is synthesized from GGPP through the activity of a class II/class I TPS pair and a CYP from the CYP701A subfamily^43^. Given the characterization above for a class II/class I TPS pair, it is plausible that the gene responsible for converting **2** to **3** is a recent duplicate of a CYP701A, especially given the occurrence of polyploidization within the Delphinieae tribe (containing *Aconitum* and *Delphinium*) of the Ranunculaceae family^44–46^.

In contrast to the TPS family, the identification of CYPs presents a challenge due to the number of genes that may be present in any given plant^47^. In our transcriptome assemblies for *D. grandiflorum* and the six *Aconitum* species, a BLAST search against a reference set of CYP sequences (Supplementary Dataset 1) yielded 2,123 predicted CYP transcripts after clustering by 99% sequence identity, with 284 for *D. grandiflorum* alone and roughly two to four hundred for each *Aconitum* assembly.

To narrow this down to a manageable number to test, we took advantage of the assumed conservation of this pathway between neighboring genera and tissue-specific accumulation of metabolites. The total CYP transcripts from each assembly were first assigned to individual clans through a sequence-similarity network, and individual phylogenies were made for distinct clans (Figure 3A and Supplementary Figures 5-7). We then filtered these transcripts to include only those in *D. grandiflorum* with high root expression and with a root-expressed ortholog in each *Aconitum* assembly. Expression values for each pathway gene are given in Supplementary Figure 8, and orthologs were inferred from grouping in phylogenetic trees as in the example in Figure 3A and shown in Supplementary Figures 5-7. This narrowed a list of 284 possible CYPs from *D. grandiflorum* down to just six for testing.

**Figure 3:**
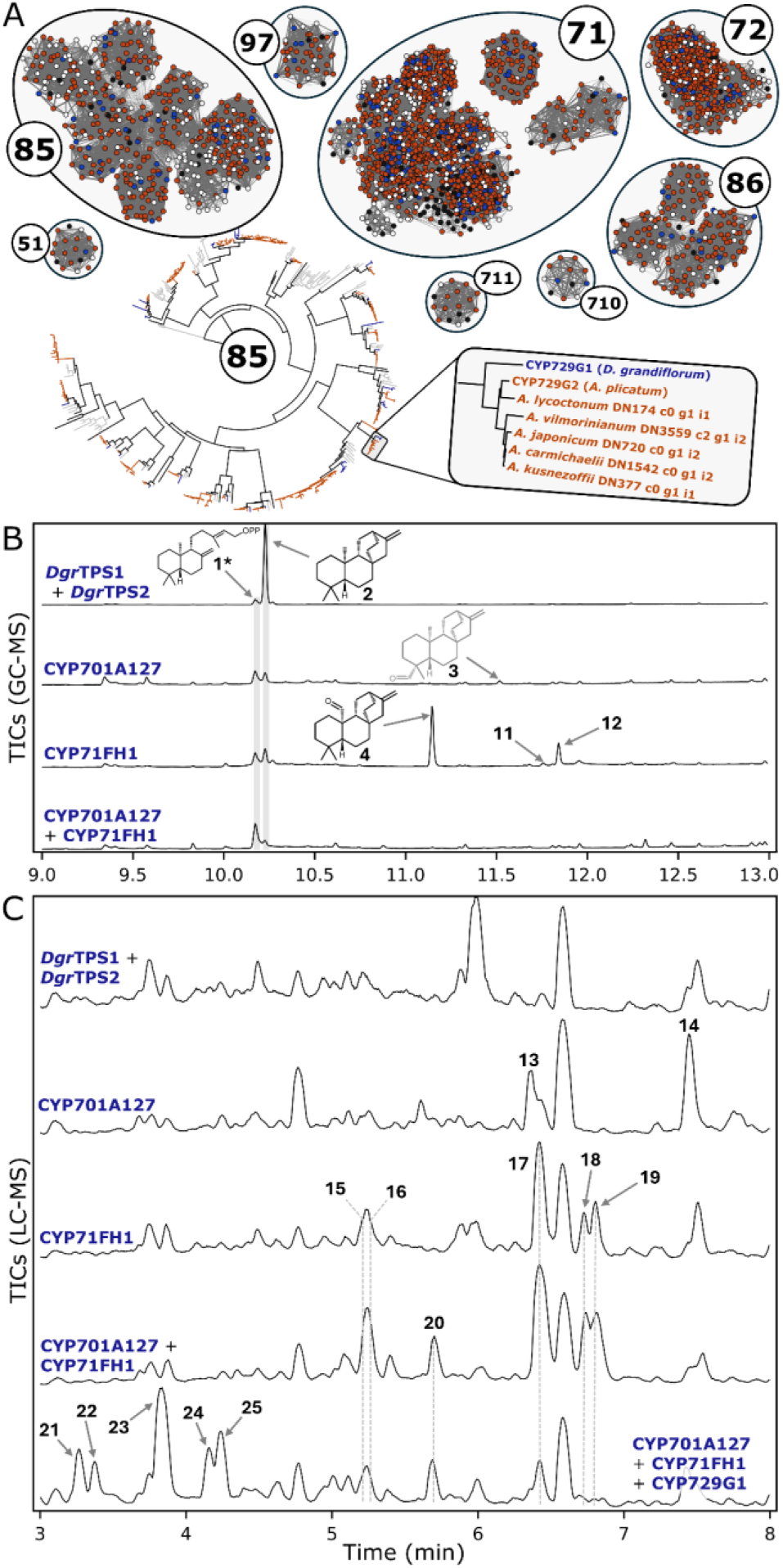
Search strategy and characterization of three CYPs active in the pathway. **A**) Visual representation of the search strategy for candidate CYPs. CYPs mined from each transcriptome were grouped into distinct clans through a sequence similarity network, and each clan was built into an individual phylogenetic tree. Groups of CYPs which match the relative speciation pattern shown in the example–which also had high root expression–were selected for testing. *D. grandiflorum* sequences in blue, *Aconitum spp.* in orange, other Ranunculaceae in white, and reference sequences in black **B**) GC-MS analysis and **C**) LC-MS analysis of CYP candidates expressed in *N. benthamiana*. All assays have *Cf*DXS, *Cf*GGPPS, *Dgr*TPS1, and *Dgr*TPS2 coexpressed in addition to those listed. Mass spectra for all compounds are given in Supplementary Figures 9 and 12.

These six CYPs were cloned from *D. grandiflorum* root cDNA and tested through transient expression in *N. benthamiana*. Each candidate was coexpressed with *Dgr*TPS1 and *Dgr*TPS2, and products were analyzed via GC-MS following ethyl acetate extraction. CYP701A127 and CYP71FH1 both showed activity in oxidizing the *ent*-atiserene (**2**) backbone (Figure 3B). Coexpression with either of these CYPs showed a depletion in **2** and the production of respective metabolites with a molecular ion at *m/z* 286 and retention of *m/z* 257 as the highest abundance fragment ion (Compounds **3** and **4**; Supplementary Figure 9), consistent with two sequential oxidations of **2** to a carbonyl. A major product with a molecular ion at *m/z* 300 (compound **12**) is also seen with CYP71FH1, which would suggest a net addition of two oxygen atoms and four oxidations from **2**.

For the products of CYP71FH1, we scaled up biosynthesis in *N. benthamiana* to purify compounds and attempt to solve structures by NMR. While sufficient quantities were simple to produce through expression and extraction from roughly 30 g of infiltrated tissue, purification of the two major products from each other proved challenging. One fraction purified through a silica column was sufficiently enriched for the *m/z* 286 product that we could confirm the identity of **4** as *ent*-atiserene-20-al through NMR (Supplementary Figures 10-11). The structure of the *m/z* 300 product (**12**) was not determined. The products of CYP701A127 gave weak signals by GC and may have been shuttled away to other products through conversion by endogenous *N. benthamiana* enzymes. We tentatively assigned CYP701A127’s product as *ent*-atiserene-19-al (**3**) based on its similar fragmentation pattern to **4** (Supplementary Figure 9) and phylogenetic placement in the CYP701A subfamily (Supplementary Figure 5).

In our proposed biosynthetic pathway, we assumed that a pair of CYPs could oxidize both methyl groups at carbons 19 and 20 to aldehydes, and so we tested whether coexpression of both of these enzymes would further the pathway. Coexpression of each CYP and TPS revealed a depletion of both **2** and of both CYPs’ respective products when analyzing ethyl acetate extracts by GC-MS (Figure 3B). These assays were also analyzed by LC-MS on 80% methanol extracts, which revealed two products detectable through coexpression of CYP701A127 (**13** and **14**), five from CYP71FH1 (**15**-**19**), and one additional product with coexpression of both enzymes (**20**) (Figure 3C and Supplementary Figure 12). Four of the products present with both CYPs coexpressed are an accumulation of those observed with CYP71FH alone (compounds **15**-**19**, including its major product **17**), suggesting that these are products different than those detected by GC-MS for CYP71FH1 alone. This suggests that CYP71FH1 may share a partial functional redundancy with CYP701A127 in oxidizing the C19 methyl group (Supplementary Figure 13). We cannot rule out, however, the possibility that an endogenous *N. benthamiana* enzyme (in particular a member of the CYP701A subfamily) could carry out the function of CYP701A127 in its absence when each other pathway enzyme is present.

We further characterized this pair of CYPs against the remaining four candidates. Coexpression of both TPSs, both CYPs, and each remaining CYP candidate revealed that CYP729G1 can oxidize products formed by CYP701A127 and CYP71FH1 (Figure 3C). The molecular ions for products **21-25** suggest that they are each a single hydroxylation difference (additional 16 m/z) from major products for CYP701A127 and CYP71FH1 (**15**-**19**) alone (Supplementary Figure 12, and Supplementary Table 4). Additional CYP products observed in trace quantities are given in Supplementary Figure 14 and Supplementary Table 4. Given that CYP701A127 and CYP71FH1 work in tandem and are both likely to carry out oxidations of relevant methyl groups to aldehydes, we opted to test candidates for further steps in the pathway to account for the possibility that the large number of products may accumulate primarily due to the absence of a downstream enzyme.

### Continuation of the Previously Proposed Biosynthetic Pathway

In many alkaloid biosynthetic pathways, the formation of an alkaloid scaffold typically involves the accumulation of both an amine and aldehyde precursor^48^. The nitrogen present in the majority of diterpenoid alkaloids in *Aconitum* and *Delphinium* appears to be derived from ethylamine due to the attached -CH_2_CH_3_ group (e.g. aconitine; Figure 1), while some metabolites presumably incorporate methylamine (-CH_3_) or ethanolamine (-CH_2_CH_2_OH)^15,16^—the origin of which could come from decarboxylation of alanine, glycine, or serine, respectively. Serine decarboxylases are present in primary metabolism, and ethanolamine is a major component of plant phospholipids and is present as a free compound in plant cells^49^. A duplication of one of these enzymes in *Camellia sinensis* has been shown to decarboxylate alanine into ethylamine (AlaDC) in theanine biosynthesis^50^, although this is a specialized metabolism enzyme which is not present across all plants. Additionally, *Spiraea japonica*—an evolutionarily distinct plant which makes similar compounds—has been shown to produce isotopically labeled diterpenoid alkaloids through the addition of labeled serine^51^.

In contrast to the steps elucidated thus far, the reaction of an amine and aldehyde to form an alkaloid scaffold could occur either spontaneously or through enzyme catalysis given the inherent reactivity between aldehydes and primary amines. Many of the major compounds detected by LC-MS in CYP screening have exact masses consistent with the product of a condensation between a diterpenoid scaffold and ethanolamine (Supplementary Table 4). While the predicted dialdehyde products **5** and **8** were not directly observed, the exact masses for the major products **17** (predicted neutral formula C_22_H_33_NO_2_) and **23** (predicted neutral formula C_22_H_33_NO_3_) seen in our LC-MS data (Supplementary Figure 12) are consistent with condensation of ethanolamine with **5** and **8**, respectively. An iminium ion formed through condensation of an amine and aldehyde is inherently unstable, and quenching of this cation through either a substitution or reduction^48^ can avoid hydrolysis separating them back into their constituent parts. In the case of diterpenoid alkaloids, it likely follows both mechanisms based on the number of carbon-carbon bonds present on both attachment points for the nitrogen-containing group (Figure 4A). Carbon 20 almost always contains an extra carbon-carbon bond relative to **2** and the intermediate **4**, while carbon 19 does not, similar to both **2** and the putative intermediate **3**. This suggests that incorporation at carbon 19 involves a reductase, and at carbon 20 a spontaneous intramolecular condensation.

**Figure 4:**
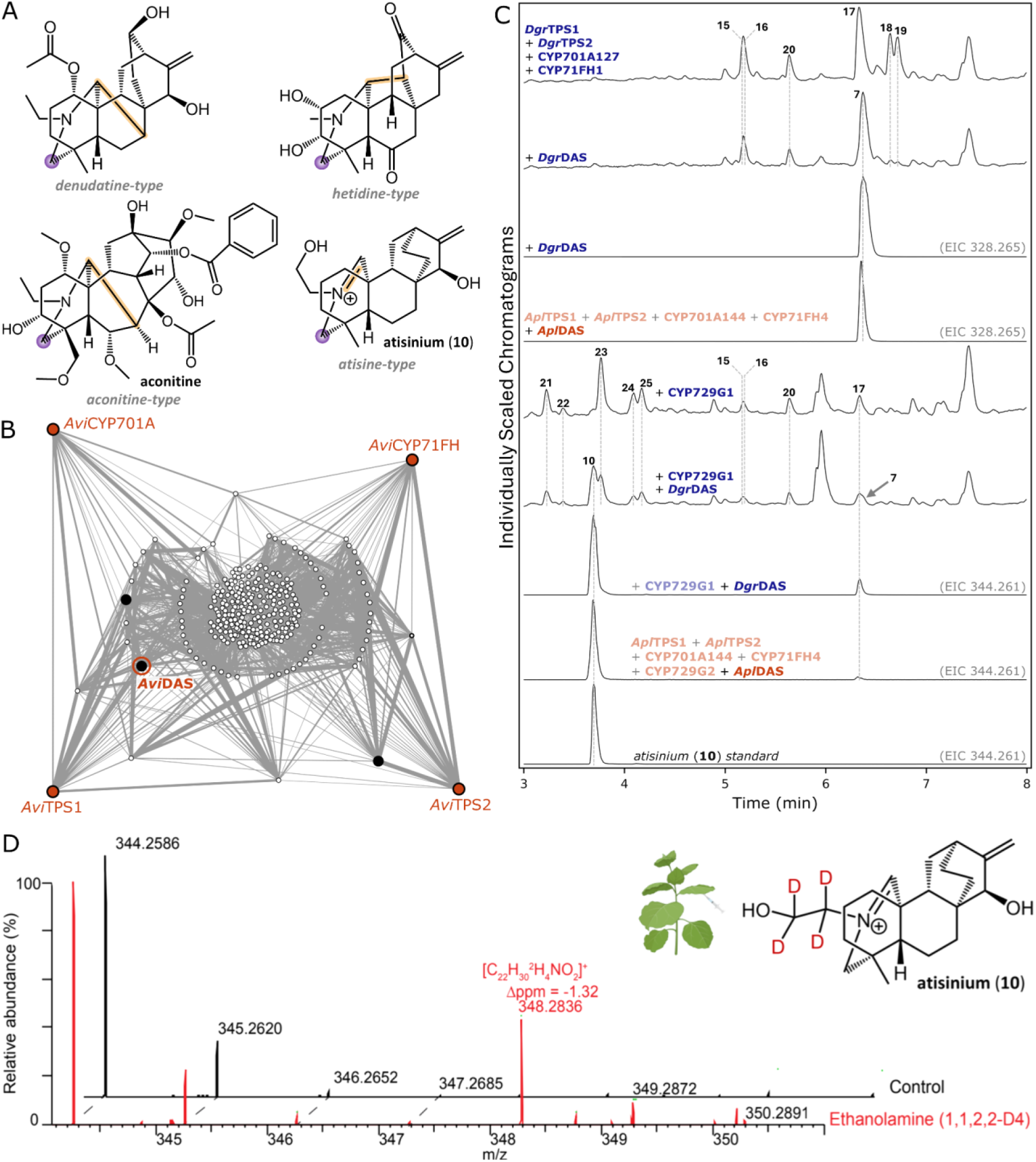
Nitrogen incorporation into diterpenoid alkaloids. **A**) Four example skeleton types of diterpenoid alkaloids. Highlighted in purple is a common attachment point (C19) for nitrogen-containing groups that does not typically form an additional carbon-carbon bond, and in yellow are examples of additional carbon-carbon bonds formed on the nitrogen-containing group’s second attachment point (C20), or a second carbon-nitrogen bond in an unresolved iminium ion in the case of **10**. **B**) Coexpression analysis on *A. vilmorinianum* roots. Four predicted orthologs to the first four pathway genes are drawn as orange nodes, and all genes coexpressed with these four are drawn in a network with nodes representing transcripts and edge width representing magnitude of coexpression between them. Transcripts connected to a greater number of orange nodes are shown further to the outside, with transcripts in the center only connected by two degrees of separation. Drawn in black are three putative reductases which were selected for testing, with DAS highlighted. **C**) LC-MS chromatograms for putative reductase testing in *N. benthamiana*. All genes have *Cf*DXS and *Cf*GGPPS coexpressed in addition to those listed, as well as *Dgr*TPS1, *Dgr*TPS2, CYP701A127, and CYP71FH1, or their respective *A. plicatum* orthologs. *D. grandiflorum* genes are written in blue and respective orthologs of each gene from *A. plicatum* are written in orange. Chromatograms are TICs except where otherwise indicated. **D**) Isotope pattern of atisinium, after coinfiltration of isotopically labeled ethanolamine (1,1,2,2-D_4_) with Agrobacteria carrying atisinium-forming genes from *A. plicatum* in *N. benthamiana*.

The search for a putative reductase is not straightforward when considering how many different enzyme families from which this function could evolve. To search for the next step(s), we carried out a coexpression analysis to identify genes which were coexpressed with the first four enzymes already characterized in the pathway (*Dgr*TPS1, *Dgr*TPS2, CYP701A127, and CYP71FH1). Given that our RNA sequencing only contained single replicates of each tissue type, we carried out this analysis on data collected for *A. vilmorinianum* instead (Figure 4B), which involved sequencing three replicates of root tissue at three different stages of development^31^. Among the genes found to be highly coexpressed with the *A. vilmorinianum* orthologs of our first four pathway genes, three were identified as putative reductases, and so the respective orthologs from *D. grandiflorum* and *A. plicatum* were selected for characterization.

### Coexpression Analysis Reveals that a Predicted Reductase is Active in the Pathway

Each of these three putative reductases were cloned from *D. grandiflorum* root cDNA and tested for activity through transient expression in *N. benthamiana* and LC-MS analysis on 80% methanol extracts. Testing of each candidate was carried out along with either the first four pathway enzymes (*Dgr*TPS1, *Dgr*TPS2, CYP701A127, and CYP71FH1) or these four plus CYP729G1. In addition, we cloned every respective *A. plicatum* ortholog of each CYP listed above and included these in testing for the same combinations.

Coexpression of *Dgr*DAS (diterpenoid alkaloid synthase) with only the first four pathway enzymes led to a suppression in previously observed products (**15**-**20**) and the formation of a new peak **7** with a proposed neutral formula of C_22_H_33_NO (exact mass 328.2656 in ESI+; calculated m/z 328.2640: Figure 4C and Supplementary Figure 15). Coexpression of *Dgr*DAS with the first four pathway enzymes and CYP729G1 did not suppress all of CYP729G1’s products (**21**-**25**), however did lead to the formation of a new peak **10** which was identified as atisinium (Figure 4C and Supplementary Figures 15 and 16), a diterpenoid alkaloid with antiplasmodial activity^23^. Comparison of MS/MS fragmentation spectra of **10** and **7** showed highly similar fragmentation patterns, thus supporting their proposed structural similarity and the role of CYP729G1 in the biosynthetic pathway of atisinium (Supplementary Figure 17). To confirm the conservation of this biosynthetic step between both genera, the *A. plicatum* DAS ortholog (*Apl*DAS) was cloned and characterized, which also resulted in atisinium (**10**) formation upon coexpression with *A. plicatum*’s respective orthologs for preceding pathway genes (Figure 4C). Notably, phylogenetic analysis reveals that *Dgr*DAS and *Apl*DAS (Pfam domain PF13460) belong to a distinct clade with homologs found in green algae, bryophytes, lycophytes, and angiosperms, however no other enzyme within this clade has been characterized to our knowledge (Supplementary Figure 18 and Supplementary Table 2). Comparison of MS/MS fragmentation spectra of **10** and **7** showed highly similar fragmentation patterns, thus supporting their proposed structural similarity and the role of CYP729G1 in the biosynthetic pathway of atisinium (Supplementary Figure 17).

Although the nitrogen in **7** and **10** appears to be derived from ethanolamine, consistent with predicted chemical formulas for **17** and **23**, many diterpenoid alkaloids contain an ethylamine group. We carried out an initial test to determine the source of nitrogen through coexpression of an alanine decarboxylase from *C. sinensis* (AlaDC)^48^, which would presumably increase the supply of ethylamine *in planta*. We observed very little change in product profile upon coexpression of this enzyme with preceding pathway enzymes except for trace amounts of one additional peak per combination (**26**, or **27** with CYP729G1 included). The compounds **26** and **27** have exact masses consistent with a condensation between ethylamine and the putative intermediates **5** and **8** (predicted neutral formulas C_22_H_33_NO and C_22_H_33_NO_2_, respectively), although **17** and **23** (or **7** and **10** with addition of *Dgr*DAS) were still seen as the major products (Supplementary Figure 15). We therefore sought to test the substrate specificity of *Apl*DAS to assess its potential involvement in formation of various scaffolds that serve as branching points in diterpenoid alkaloid biosynthetic pathways. In addition to the heterologous expression of characterized genes in *N. benthamiana*, we co-infiltrated isotopically labeled ethanolamine (1,1,2,2-D₄) or ethylamine (D₅). Atisinium (**10**) isotopic peak distribution demonstrated the incorporation of labeled ethanolamine (Figure 4D), whereas no novel peaks corresponding to an ethylamine-derived analogue of atisinium were detected. We observed a similar incorporation of labeled ethanolamine in compounds **17** and **23** in the absence of *Apl*DAS (Supplementary Figure 19). This raised the question of whether ethanolamine is the preferred source of nitrogen across the majority of diterpenoid alkaloids, or if DAS is simply a branching point towards a more narrow range of ethanolamine-containing diterpenoid alkaloids.

### Ethanolamine as a preferred source of nitrogen in Aconitum diterpenoid alkaloids

In order to better understand the biosynthesis of diterpenoid alkaloids and particularly the source of nitrogen, we attempted to establish *A. plicatum* hairy root cultures and to perform labeled substrate feeding in sterile plant cuttings. However, hairy root cultures could not be established and sterile cuttings did not remain viable under the set incubation conditions. We therefore developed callus cultures of *A. plicatum* as an alternative system. Comparative metabolomic analysis of callus cultures and *A. plicatum* tissues (leaf, root and flower) showed that callus cultures produce a significant amount of terpenoid alkaloids, indicating that they represent a suitable model for investigation of DA biosynthesis (Figure 5A). Established callus cultures were fed with either isotopically labeled ethanolamine (1,1,2,2-D_4_), ethylamine (D_5_), or 1 x PBS (control) over the course of one month. Due to the scarcity of available standards and lack of diterpenoid alkaloids in spectral libraries, we opted for computational solutions, in order to obtain a list of all detected and predicted diterpenoid alkaloids from callus LC-MS analysis (Supplementary Figure 20; See Methods for details). Computational solutions for metabolite annotation problems in untargeted metabolomics have seen an expansion in recent years, with SIRIUS^52^ being one of the most prominent. It relies on isotope patterns and fragmentation trees for molecular formula prediction, with further steps involving formula ranking (ZODIAC^53^), prediction of a molecular fingerprint of the query compound through CSI:FingerID^54^ and compound class prediction (CANOPUS^55^). Relying on this approach, features were detected and aligned across all samples using MZmine 3^56^, ensuring consistent *m/z* and retention time assignment for each feature across feeding conditions. Out of 1,331 aligned features with MS/MS spectra, 144 were classified by SIRIUS as belonging to terpenoid alkaloids. Filtering only for features that were consistently present across the three feeding conditions, defined as being detected in at least one replicate of each condition, as well as the ones whose classification had a probability score higher than 0.6 further reduced the list to 60 putative terpenoid alkaloids. An additional feature, atisinium (**10**), was added manually, as it was not ranked as the top annotation by the computational pipeline and was consequently filtered out in later steps, thus resulting in a final table with 61 detected features. Interestingly, out of those, 41 features exhibited presence of deuterium-labeled isotopologue peaks exclusively in ethanolamine-fed callus cultures. More specifically, two features showed exclusively the M+4D isotopologue, six features displayed both M+3D and M+4D isotopologues, while the remaining features showed only the M+3D isotopologue (Figure 5B). No features showing the presence of isotopologue corresponding to M+5D were detected with our pipeline. Both atisinium and aconitine showed incorporation of deuterium atoms in the ethanolamine-feeding condition, as evident from their isotope patterns (Figure 5C, D). The incorporation was further supported with MS/MS spectra, which confirmed the mass shift while retaining the same fragmentation pattern for both compounds (Supplementary Figure 21). To further investigate the origin of nitrogen in these compounds, callus cultures were fed with isotopically labeled ʟ-serine (2,3,3-D₃, ^15^N), resulting in a low-intensity isotopologue peaks for both compounds, consistent with incorporation of three deuterium atoms and one ^15^N (Supplementary Figure 22), in agreement with serine decarboxylation to ethanolamine.

**Figure 5:**
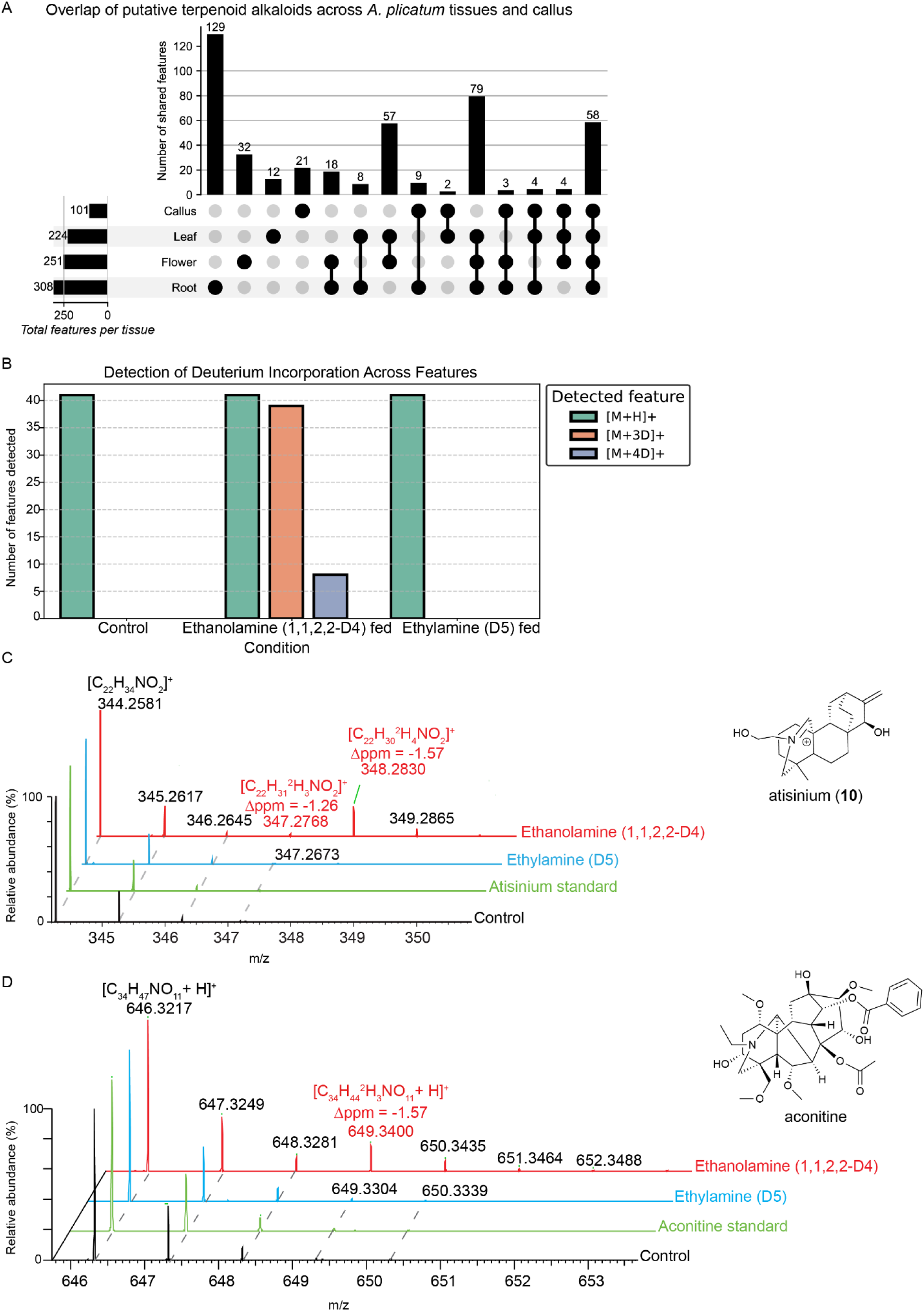
Deuterium incorporation in callus culture. **A)** An UpSet plot showing shared and unique putative terpenoid alkaloids across callus cultures and leaf, flower and root tissue of *A. plicatum.* Filled circles indicate presence of a feature in the given tissue, while vertical bars represent the number of features shared among the indicated tissue combinations. **B)** LC-MS features classified as “Terpenoid alkaloids” by CANOPUS across feeding conditions for which M+3D and M+4D isotopologues were detected, indicating the role of ethanolamine in their biosynthesis. As no features corresponding to M+5D were detected, this is omitted from the figure. **C)** Isotope pattern of atisinium, showing isotopologue peaks corresponding to the incorporation of three and four deuterium atoms in ethanolamine-fed callus cultures. D) Isotope pattern of aconitine, showing the isotopologue peak corresponding to incorporation of three deuterium atoms in ethanolamine-fed callus cultures.

Despite ethylamine being intuitively inferred as the source of nitrogen, given its presence as the side group in the majority of diterpenoid alkaloids (aconitine included, Figure 5D), no incorporation was observed. The difference in the number of detected putative terpenoid alkaloids and those with incorporated labeled substrate could reflect the presence of stored metabolites versus those actively synthesized during the feeding. Furthermore, the detection of secreted labeled diterpenoid alkaloid compounds in the agar could suggest their role in plant-plant communication, which has been poorly investigated so far^57^.

## Discussion

Through a combination of comparative transcriptomics and coexpression analysis, we have identified six enzymes sufficient to reconstitute the biosynthetic pathway towards diterpenoid alkaloids conserved between the *Delphinium* and *Aconitum* genera within the Ranunculaceae family. There are hundreds of diterpenoid alkaloids in this family, and the identification of these enzymes will serve as the basis for further pathway discovery towards specific metabolites. This work highlights the utility of cross-referencing transcriptomic data between genera as an orthogonal filter for selection of candidate enzymes beyond the analysis of a single species, as it likely would not have been possible to identify all of these enzymes otherwise given the inherent complexity of these pathways.

The size and ploidy of genomes throughout *Delphinium* and *Aconitum* vary widely, exemplified by *Delphinium montanum*, which is an autotetraploid with a predicted genome size of roughly 40 Gb^46^ (2n = 32^58^). The seven species studied here have a range of predicted ploidy levels (*D. grandiflorum*: 2n = 16; *A. carmichaelii*: 2n = 32/64 – depending on cultivar; *A. japonicum*: 2n = 32; *A*. *vilmorinianum*: 2n = 16; *A. kusnezoffii:* 2n=32; *A. plicatum:* 2n=32; *A. lycoctonum: 2n=16*)^58–60^, and it has been suggested that, at least in the *Aconitum* genus, there may have been multiple recent events of polyploidization and diploidization^45^. This fits with the model of our initial biosynthetic pathway—and the phylogenetic relationships of these genes—in which we predicted that the first three steps may be recent duplications of primary metabolism enzymes given the similarity of these predicted intermediates to those in gibberellin biosynthesis^43^. Following our characterization of the TPS pair which produce *ent*-atiserene (**2**), Mao *et al.*^37^ published a characterization of the entire TPS family from *A. carmichaelli* and identified enzymes orthologous to ours (*Dgr*TPS1/*Apl*TPS1 and *Dgr*TPS2/*Apl*TPS2), as well as recently-duplicated paralogs of each which produce *ent-*CPP (**1**) and *ent*-kaurene. Very recently, Luo, Zhou, *et al*.^38^ identified CYPs which oxidize the *ent*-atiserene backbone from *A. carmichaelli* and *A. coreanum*, including a member of the CYP701A subfamily (typically involved in gibberellin biosynthesis^61^) which is orthologous to our CYP701A127 and CYP701A144 in addition to an ortholog of CYP71FH1 and CYP71FH4. It remains to be seen whether duplications of downstream gibberellin pathway genes could also be involved in further biosynthetic steps in diterpenoid alkaloid biosynthesis.

It should be noted that *Dgr*TPS1 and *Apl*TPS1—being *ent*-CPP (**1**) synthases—are technically not enzymes that produce specialized metabolites. Given their high and exclusive expression in roots relative to their putative central metabolism paralogs, however, they are likely dedicated to specialized metabolism. A similar phenomenon is seen in both *Oryza sativa*^62^ and *Zea mays*^63^, where two copies of an *ent*-CPP (**1**) synthase are present; one which is involved in gibberellin biosynthesis and another which is inducible by pathogens for the production of defensive *ent*-CPP-derived specialized metabolites. Given the presence of duplicate *ent*-CPP (**1**) synthases in each of these independent lineages of plants, there is likely a strong evolutionary pressure for the ability to tightly regulate these competing pathways.

As we transitioned between different classes of enzymes in the pathway, we varied the approach to identify each based on what information was necessary. For the TPSs, for example, few enough transcripts were present in our assembly that we relied solely on data from *D. grandiflorum*. For the CYPs, cross-referencing both *Delphinium* and *Aconitum* datasets was essential given the presence of two to three hundred unique transcripts in each assembly. Orthologous genes present across each species have persisted over roughly 27 million years since the speciation between the common ancestors of each genera^64^, and choosing to work across both allowed us to filter these hundreds of candidates down to just six. Finally, even with tissue and species-specific transcriptomic data, the following step for nitrogen incorporation was not obvious; therefore, coexpression analysis allowed us to search for new candidates without prior knowledge of which enzyme families to search.

Throughout the process of characterizing various steps in the pathway, not every intermediate product was identified prior to moving forward with following steps. Often it can be difficult to differentiate “actual” intermediates in terms of whether the observed products are relevant to the pathway of interest, or the result of an incomplete reconstruction of the pathway or a heterologous host’s interference. In the process of discovering the biosynthetic pathway for the diterpenoid forskolin from *Coleus forskohlii*, for example, coexpression of an incomplete set of genes in *N. benthamiana* led to an accumulation of many side products that were not present once the entire pathway was reconstructed (five CYPs acting on a single diterpene scaffold and at least sixteen total products)^65^. A similar–but deconstructive–example can be seen with accumulation of precursors and side products for the scopolamine pathway in *Atropa belladonna* following virus-induced gene silencing of various pathway steps^6^. This may be reflective of the capacity for specialized metabolism to evolve towards a range of different products with only a small change (e.g. a single gene deletion) in their respective biosynthetic pathways. We identified the activity of the two TPSs and confirmed our predicted activity of the first two CYPs, but thereafter we decided to test enzymes in different combinations to identify new steps in case the side products seen were due to a similar phenomenon. Given the appearance of multiple products with the first four pathway enzymes and convergence to primarily a single product **7** upon coexpression with *Dgr*DAS or *Apl*DAS, the initial range of products beyond the predicted intermediate **17** could be the result of incomplete pathway reconstruction.

We initially proposed that ethylamine was the source of nitrogen in this pathway due to the abundance of diterpenoid alkaloids that have been identified with an ethyl group bonded to the nitrogen. The presence of a minor product forming upon coexpression with AlaDC–which would presumably increase the supply of ethylamine–was expected based on the presence of reactive aldehydes in our intermediates, however, very little product was observed in this case. We demonstrate here through isotope feeding in *N. benthamiana* that the preferred substrate for *Apl*DAS is ethanolamine, and in *A. plicatum* callus cultures that diterpenoid alkaloids generally incorporate ethanolamine preferentially over ethylamine, even in cases like aconitine where an ethylamine group is present in the final product. It remains to be seen how this is implicated in further biosynthetic steps towards more complex diterpenoid alkaloids. Similarly, probing substrate specifics and kinetics of *Apl*DAS through recombinant assays *in vitro* could be valuable in complementing our *in planta* characterization. We attempted engineering CYP-based substrate production through yeast microsomes^66^. However, our results indicated the need for substantial optimization for recombinant CYPs.

Beyond this immediate question, more remain in the discovery of diterpenoid alkaloid pathways. Perhaps the most important is the differentiation between the C20 and C19/C18 metabolites, and at which point this occurs. With the central scaffold-forming steps investigated here, which are presumably shared between all diterpenoid alkaloids, further pathway discovery work towards more highly decorated metabolites within this class can be investigated. Some may be a significant challenge, exemplified by some of the compounds shown in Figure 1, as there are likely many more enzymatic steps involved in their biosynthesis. Further discovery of such downstream steps will likely require different methodology than employed here given the species-and tissue-specificity of some of these products. The hydroxylation introduced by CYP729G1 and CYP729G2 appears to be present in many C20 compounds^24^, however it remains to be seen if this may be implicated in the rearrangement towards C19 compounds and if **7** and **10** represent metabolic branch points, as not all CYP729G orthologs across species studied here have identical expression patterns across tissue types relative to other pathway enzymes (Supplementary Figure 8). Considering this variable expression, further work may benefit from correlating metabolomic and transcriptomic data to differentiate chemical conversions present in distinct lineages and tissue types in conjunction with the cross-species transcriptomics employed here.

## Materials and Methods

### Plant material, RNA isolation, and cDNA synthesis

*D. grandiflorum* plants were grown in a greenhouse under ambient photoperiod and 24 °C day/17 °C night temperatures. RNA isolation, quality assessment, RNA sequencing, and cDNA synthesis for *D. grandiflorum* was carried out as described in Miller *et al.* 2020^67^. Total RNA was isolated from flowers, leaves, and roots with the Spectrum Plant Total RNA Kit (Sigma-Aldrich), and DNA was removed with the DNA-free DNA Removal Kit (Thermo Fisher Scientific). Quality of the resulting RNA was assessed with a Qubit (Thermo Fisher Scientific) and RNA-nano assays (Agilent Bioanalyzer 2100). RNA sequencing was carried out on an Illumina HiSeq 4000 at Novogene (Sacramento, CA, USA) *Aconitum plicatum* and *Aconitum lycoctonum* plants were collected from the Charles University Botanical Garden. For these species, RNA was isolated from roots, flowers, leaves, stems, and (for *A. lycoctonum*) fruits using the RNeasy kit (Qiagen) according to the manufacturer’s instructions. The transcriptome library preparation and sequencing were performed at the Beijing Genomics Institute using the DNBSEQ Eukaryotic Strand-specific mRNA library.

### D. grandiflorum and Aconitum spp. de novo transcriptome assembly and analysis

RNA-seq data were obtained through RNA sequencing on an Illumina HiSeq 4000 for *D. grandiflorum*, and from DNBseq platform for *A. plicatum* and *A. lycoctonum*. Raw data are accessible on the NCBI Sequence Read Archive (SRA; ncbi.nlm.nih.gov/sra) under the accession PRJNA1261909. Additional data for *A. carmichaelii* (PRJNA415989)^33^, *A. japonicum* (PRJDB4889), *A. kusnezoffii* (PRJNA670255), and *A. vilmorinianum* (PRJNA667080)^31^ were obtained from the SRA. Reads were filtered for quality by a Phred score of 10 or higher and adapters trimmed with TrimGalore (v0.6.10; github.com/FelixKrueger/TrimGalore). Unless otherwise specified, a maximum of 150 million read pairs (or single-end reads) were used for each transcriptome assembly, and were taken evenly across samples.

Transcriptome assembly and analysis followed a similar pipeline as described in Miller et al. 2020^67^. Transcriptomes were assembled *de novo* with Trinity (v2.8.5)^68^ and open reading frames and translations were found with TransDecoder (v5.7.1)^69^. Transcriptomes were clustered at 99% sequence identity by local alignment to at least 15% of the larger sequence with CD-HIT (v4.8.1)^70,71^, and any sequences filtered out by this step were also filtered out from the translated TransDecoder output. Expression levels were calculated with Salmon (v0.14.1)^72^ with the transcriptome filtered by CD-HIT used as the index. The results of this analysis are accessible at doi.org/10.5281/zenodo.15384649, and a Nextflow pipeline which automates this process can be found at github.com/GarretPMiller/DiterpenoidAlkaloids.

Initial assembly of the *D. grandiflorum* transcriptome resulted in incomplete transcripts for *Dgr*TPS1 and *Dgr*TPS2 (only ∼75% coverage of reference sequences), and although this was prior to our characterization of these enzymes, we noted that these transcripts were most likely misassembled given their high expression and likelihood of being involved in the pathway. Reassembly of the *D. grandiflorum* transcriptome was therefore done with only data acquired from root tissue, with reads from each tissue type mapped to this assembly. Transcripts for both of these genes in the new assembly aligned to the entire length of reference sequences, and so this assembly was used for further analysis.

### Sequence Similarity Networks and Phylogenetic Trees

Sequences from all transcriptome assemblies, predicted genes from two other Ranunculaceae species (*Thalictrum thalictroides*: PRJNA439007; *Coptis chinensis*: PRJNA662860), and reference sequences (given in Supplemental Dataset 1) were used to build the sequence similarity network and all phylogenetic trees for TPS and CYP discovery. The CYP sequence similarity network was made with BLAST (v2.7.1+)^73^ at a sequence identity threshold of 42% and visualized with Cytoscape^74^. Sequences which only had one connection (primarily fungal contaminants and low percent identity reference sequences) were removed from the network. Multiple sequence alignments used for phylogenetic tree construction were made with Clustal Omega (v1.2.4)^75^ (--iterations 3, otherwise default settings), and maximum likelihood phylogenetic trees were made with RAxML (v8.2.12)^76^ (model: PROTGAMMAAUTO; algorithm: a). All trees were the result of 1,000 bootstrap replicates, except where otherwise indicated. Outgroups are indicated in their respective figure legends. Vector images of all trees used for TPS and CYP discovery are accessible at doi.org/10.5281/zenodo.15384649.

Sequences homologous to *Dgr*DAS and *Apl*DAS were mined from each *Delphinium* and *Aconitum* transcriptome assembly in addition to predicted genes from a wide range of model species as shown in Supplemental Table 2. All sequences meeting a BLASTp threshold of 1e-10 with *Dgr*DAS as the query sequence from these assemblies, model species, and UniProt were grouped into a sequence similarity network with a sequence identity threshold of 41%. All sequences which were not separated entirely from *Dgr*DAS were further filtered by clustering at 100% identity (by amino acid sequence, as previous clustering was done at the transcript level) and by those with an alignment length of at least 110 amino acids (∼45% of query sequence) in the initial BLASTp search. The phylogenetic tree shown in Supplemental Figure 18 was made as described above, with the exception that no outgroup was specified.

### Coexpression analysis

Our assembly for *A. vilmorinianum* was used for coexpression analysis. To minimize the computational burden, we reduced the analysis through clustering by 99% identity with CD-HIT (v4.8.1)^70,71^, calculated expression levels through mapping reads to this clustered transcriptome, and eliminated any transcript with no samples that had at least 20% the expression level (in TPM) as any sample for either TPS. Coexpression analysis was carried out as described by Wisecaver *et al.* 2017^77^ (pipeline at: https://github.itap.purdue.edu/jwisecav/mr2mods). The resulting coexpression network shown in Figure 4B shows only genes with one or two degrees of separation from any of the first four genes in the pathway (respective orthologs from *A. vilmorinianum* of *Dgr*TPS1/*Apl*TPS1, *Dgr*TPS2/*Apl*TPS2, CYP701A127/CYP701A144, and CYP71FH1/CYP71FH4) based on a mutual rank (MR) cutoff of e^(-(MR-1)/5) > 0.01. Putative reductases from these coexpressed genes were identified by InterProScan (v5.59)^78^ annotations.

### Cloning

Candidate genes were PCR-amplified from root cDNA and cloned into pEAQ-HT^79^ through In-Fusion cloning. Constructs for *Zm*AN2, *Nm*TPS1, and *Nm*TPS2 in pEAQ (used as positive controls for *ent*-CPP, (*+*)-CPP, and *ent*-kaurene biosynthesis, respectively) were made by Johnson et al. 2019^41^. Nucleotide (verified by Sanger sequencing) and amino acid sequences for all *Delphinium* and *Aconitum* genes tested here are given in Supplemental Dataset 1.

### Transient expression in N. benthamiana, product scale-up, and NMR analysis

Transient expression in *N. benthamiana* for screening assays was carried out as described in Miller *et al.* 2020^67^, with the exception of solvents used to extract each set of assays as described in the main text. For the co-infiltration of ethanolamine (1,1,2,2-D_4_) and ethylamine (D_5_), the compounds were dissolved in distilled water, filter sterilized, and co-infiltrated in *N. benthamiana* leaves 1 day after the candidate genes were infiltrated. For *ent*-atiserene and *ent*-atiserene-20-al scaleup, three whole plants were infiltrated with a syringe, and approximately 15/30 g of fresh weight were extracted with hexane/ethyl acetate (respectively). Products were purified through silica chromatography with 10% ethyl acetate:90% hexane as the mobile phase. Initial purification was carried out with approximately 100 mL of oven-dried silica, and fractions were collected in approximately 3 mL increments and assessed for purity by GC-FID. Fractions containing desired products were further purified with approximately 1.5 mL oven-dried silica in a Pasteur pipette, with fractions collected in 1 mL increments and purity assessed by GC-FID. NMR analysis was carried out on a Bruker 800 MHz spectrometer equipped with a TCl cryoprobe using CDCl_3_ as the solvent. CDCl_3_ peaks were referenced to 7.26 and 77.00 ppm for ^1^H and ^13^C spectra, respectively.

### GC-MS analysis

All GC-MS analyses were performed on hexane or ethyl acetate extracts (described for each case in the text) with an Agilent 7890A GC with an Agilent VF-5ms column (30 m x 250 µm x 0.25 µm, with 10m EZ-Guard) and an Agilent 5975C mass spectrometer. The inlet was set to 250°C splitless injection of 1 µL, He carrier gas (1 ml/min), and the detector was activated following a 3 min solvent delay. Mass spectra were generated using 70 eV electron ionization with a scan range of *m/z* 50 to 350. The following method was used for analysis of each sample presented in the text: temperature ramp start 40 °C, hold 1 min, 40 °C/min to 200 °C, hold 2 min, 20 °C/min to 280 °C, 40 °C/min to 320 °C; hold 5 min. For initial GC-MS analysis of *Apl*TPS1 and *Apl*TPS2 products shown in Supplementary Figure 3, the GC method varied by the following parameters: 275 °C splitless injection, scan range of *m/z* 50 to 400, 4.5 minute hold after reaching 200 °C, 20 °C/min to 240 °C, 10 °C/min to 280 °C. Figures for chromatograms and mass spectra were generated with Pyplot.

### LC-MS analysis

All LC-MS analyses (except as specified below) were performed on 80% methanol : 20% H_2_O *N. benthamiana* extracts with a Waters Xevo G2-XS quadrupole ToF mass spectrometer with a Waters ACQUITY column manager and Waters ACQUITY BEH C18 column (2.1 x 100 mm; 1.7 µm). Injection volume for each sample was 10 µL, and flow rate was set to 0.3 mL/min with a column temperature of 40 °C. The mobile phase consisted of 10 mM ammonium formate (pH 2.8) (Solvent A) and acetonitrile (Solvent B) with the following method: initial 99% A : 1 % B, continuous gradient to 2% A : 98% B over 12 min, hold for 1.5 min, continuous gradient to 99% A : 1% B over 0.1 min, hold 1.5 min. Mass spectra were generated through electrospray ionization in positive-ion mode with leucine enkephalin as a lockmass, and continuum peak acquisition were collected with a mass range of *m/z* 50-1500 and a scan duration of 0.2 s. Capillary and cone voltage were 3.0 kV and 40 V, respectively, cone and desolvation gas flow rates were 40 and 600 L/h, respectively, and source and desolvation temperatures were 100 °C and 350 °C, respectively. High-energy spectra were generated with argon as the collision gas and a voltage ramp from 20 to 80 V. Figures for chromatograms and mass spectra were generated with Pyplot.

LC-MS analyses of isotopically labeled atisinium were performed on 80% MeOH extracts of *N. benthamiana* with a Vanquish Flex UHPLC system interfaced to an Orbitrap ID-X Tribrid mass spectrometer using heated electrospray ionization (H-ESI). For LC analysis, Waters ACQUITY BEH C18 column (2.1 x 150 mm; 1.7 µm) was used. Injection per sample was 1 µL, with flow rate of 0.35 mL/min with the column temperature of 40 °C. The mobile phase consisted of water with 0.1% formic acid (Solvent A) and acetonitrile with 0.1% formic acid (Solvent B) with the following method: initial 95% A : 5 % B, continuous gradient to 100% B over 15.5 min, hold for 1.8 min, and a continuous gradient to 95% A over 2 minutes. Mass spectra were generated through H-ESI in positive mode, and continuum peak acquisition was collected with a mass range of *m/z* 100-1000. Capillary voltage was at 3.0 kV, ion transfer tube temperature was 325 °C, auxiliary gas flow rate 10 L/min, vaporizer temperature 350 °C, sheath gas flow rate 50 L/min, MS resolution 60,000 (at *m/z* 200), RF Lens 45% and maximum injection time 118 ms. The same method was used for the analysis of callus cultures, except the resolution was increased to 120,000 (at *m/z* 200) to improve the distinction of isotopic peaks arising as a result of deuterium incorporation. For ʟ-serine (2,3,3-D_3_, ^15^N) fed callus, the resolution was increased to 500,000 (at *m/z* 200) to allow separation of the ^15^N isotopic peak. As these measurements were acquired in a separate run, control samples were re-acquired under the same conditions to enable direct comparison.

### Callus culture establishment, substrate feeding and metabolite extraction

*A. plicatum* petioles were cut into 0.5 cm pieces and sterilized by immersion in 70% ethanol over 2 minutes, followed by 20 minutes in 20% bleach (sodium hypochlorite) and rinsed with sterile distilled water. The sterilized petioles were placed on ½ Murashige and Skoog (MS) medium (Duchefa Biochemie, M0222) supplemented with 0.8% agar (Sigma-Aldrich, A7921), 3% sucrose, 0.1 mg/L kinetin and 1 mg/L 1-Naphthaleneacetic acid (NAA). Cultures were incubated in darkness at room temperature until callus formation was observed. Established calli were then transferred to fresh medium and maintained under the same conditions.

For the substrate feeding studies, calli were divided into the following groups: (1) ethanolamine (1,1,2,2-D_4_)-fed (Cambridge Isotopes Laboratories, DLM-552-0.1), (2) ethylamine (D_5_)-fed (Cambridge Isotopes Laboratories, DLM-3471-0.5), (3) ʟ-serine-fed (2,3,3-D_3_,^15^N) (Cambridge Isotopes Laboratories DNLM-6863-0.25) and (4) control group to which 1x PBS was added. A sterile 1 mM solution of each substrate was applied directly on the callus surface, biweekly over the course of one month. During this period, calli were kept in darkness at room temperature. For the metabolite extraction, calli were collected in 2 mL Safe-Lock Eppendorf tubes, weighed, and extracted with ethyl acetate at a 1:10 (w/v) ratio. Samples were homogenized using a steel bead in a Qiagen TissueLyser. After centrifugation at 18,000 x g, the supernatant was collected and transferred to LC-MS vials. The same extraction protocol was used for *A. plicatum* tissues—leaf, root and flower.

### Computational metabolomics analysis

Raw LC-MS data files were converted to open format (.mzML) using MSConvert^80^ tool from ProteoWizard^81^ package. Untargeted feature detection was performed in MZmine 3^56^ (version 4.4.3). Briefly, the mass detection was based on the factor of lowest signal, which was set at 6.001 and 2.5 for MS1 and MS2 respectively. All signals below this threshold were discarded. Extracted ion chromatograms (EIC) were built for each *m/z* value that was detected over a minimum of 5 consecutive scans with minimum intensity of 1.0E5, with *m/z* tolerance between scans being 0.002 (or 10 ppm), and minimum absolute height of 5.0E5. Chromatograms were smoothed using a Savitzky–Golay filter (window size = 5 scans), after which EIC were resolved using the Local Minimum Feature Resolver module with the following parameters: chromatographic threshold= 93%; minimum search range: 0.05; minimum absolute height 5.0E4; minimum ratio of peak top/edge: 1.90; peak duration range: 0.03-0.60; minimum scans: 6. MS2 scans were paired to precursor features using an *m/z* tolerance of 0.01 (10 ppm) and retention-time constraints based on feature edges. Across all samples, features were aligned (Join aligner module) with an *m/z* tolerance of 0.002 or 5 ppm (weight=3) and a retention-time tolerance of 0.03 min (weight 1). Feature List Row Filter was used to filter features with retention time 2-17 minutes, and *m/z* 272-900. Peak Finder was used to look for missing peaks with the following parameters: intensity tolerance: 20%; *m/z* tolerance: 0.0010 (5 ppm); retention time tolerance: 0.03 minutes; minimum scans: 5. Duplicate Peak Filter module was used with following parameters: filter module: new average; *m/z* tolerance: 0.0008 (5 ppm); retention time tolerance: 0.03 minutes. This generated a feature list where each detected isotopic peak was retained as an individual feature. The second feature list was generated in MZmine 3 for SIRIUS software (v6.0.7) analysis using the same steps as the previous batch file, with addition of two extra modules which were necessary for SIRIUS analysis. Isotope grouper module was used with the following parameters: *m/z* tolerance of 0.0015 or 3 ppm; retention time tolerance: 0.03 min; maximum charge: 2; representative isotope: most intense, and Isotope Finder module was used with *m/z* constraints 0.0005 or 3 ppm and elemental constraints (H,C,N,O,S). The steps used for comparison of callus and *A. plicatum* tissues were the same as the aforementioned list, with minimum absolute height set to 5.0E5 in Local Minimum Feature Resolver module. For SIRIUS analysis of both datasets, default parameters were used except for the instrument type, which was set to Orbitrap. Molecular formula prediction was done based on isotope pattern and fragmentation tree reconstruction, with formula ranking refined using ZODIAC. Feature properties and compound class were computed using CSI:FingerID module and CANOPUS, respectively. Jupyter Notebook was used to filter SIRIUS results, creating a database of all compounds classified as terpenoid alkaloids. For each feature in the database, a theoretical *m/z* was calculated corresponding to incorporation of three, four or five deuteriums. Features in the MZmine output matching these *m/z* values within a 5 ppm mass tolerance and retention time tolerance of 0.05 min, were reported as showing incorporation of the labeled substrate The used script, as well as the batch files and the final output table can be freely accessed at https://github.com/lalalana5/Callus-culture-isotope-labelling/tree/main. Each predicted compound with incorporated deuterium was manually verified by inspecting the raw data. Features resulting from carryover between injections were excluded from further analysis.

## Data Availability

RNA sequencing data for *D. grandiflorum*, *A. plicatum*, and *A. lycoctonum* can be found at the NCBI Sequence Read Archive under the BioProject accession number PRJNA1261909. Nucleotide sequences verified by Sanger sequencing for each characterized gene can be found in GenBank with the following accessions: (*Dgr*TPS1: PV652189), (*Apl*TPS1: PV652199), (*Dgr*TPS2a: PV652190), (*Dgr*TPS2b: PV652191), (*Apl*TPS2: PV652200), (CYP701A127: PV652192), (CYP701A144: PV652201), (CYP71FH1: PV652193), (CYP71FH4: PV652202), (CYP729G1: PV652194), (CYP729G2: PV652203), (*Dgr*DAS: PV652195), (*Apl*DAS: PV652204). Additional CYP sequences from *D. grandiflorum*: (CYP71FH2: PV652196), (CYP72A932: PV652197), (CYP71FJ1: PV652198). These sequences, along with amino acid translations and all amino acid sequences of reference TPSs and CYPs used to build the phylogenetic trees can be found in Supplemental Dataset 1. Transcriptome assemblies, open reading frames, and expression values across RNA-seq samples are provided in the following Zenodo repository: doi.org/10.5281/zenodo.15384649. All phylogenetic trees are also available in this repository (doi.org/10.5281/zenodo.15384649). A Nextflow pipeline to carry out the transcriptome assembly and analysis process done here can be found at: github.com/GarretPMiller/DiterpenoidAlkaloids. Raw mzML files from the LC-MS analysis of *A. plicatum* callus culture feeding can be found in the Zenodo repository: https://doi.org/10.5281/zenodo.19150401. The script, batch files, and final output table for the computational metabolomics of callus cultures can be found at: github.com/lalalana5/Callus-culture-isotope-labelling/tree/main.

## Acknowledgments

We would like to acknowledge three core facilities at Michigan State University including the Institute for Cyber-Enabled Research, the Mass Spectrometry and Metabolomics Core, and the Max T. Rogers NMR Facility, as well as Advanced Research Computing (ARC) at the University of Michigan. We would also like to thank Sean Johnson and Gina Roeppischer for technical assistance in *D. grandiflorum* RNA isolation, Prof. Tomáš Urfus from Charles University for providing us with *A. plicatum* and *A. lycoctonum* plants, and Prof. Tetiana Satarova from the Institute of Experimental Botany, Czech Academy of Sciences, for training in callus culture initiation and maintenance.

## Author contributions

Conceptualization: GPM, LMN, TP and BH; Methodology: GPM, LMN; Software: GPM, LMN; Formal Analysis: GPM, LMN, IP, KVW, TBA; Validation: GPM, LMN, TBA, MS, AB, TI, AT; Writing - original draft & editing: GPM, LMN; Supervision: TP, BH.

## SUPPLEMENTARY INFORMATION

**S. Figure 1.**
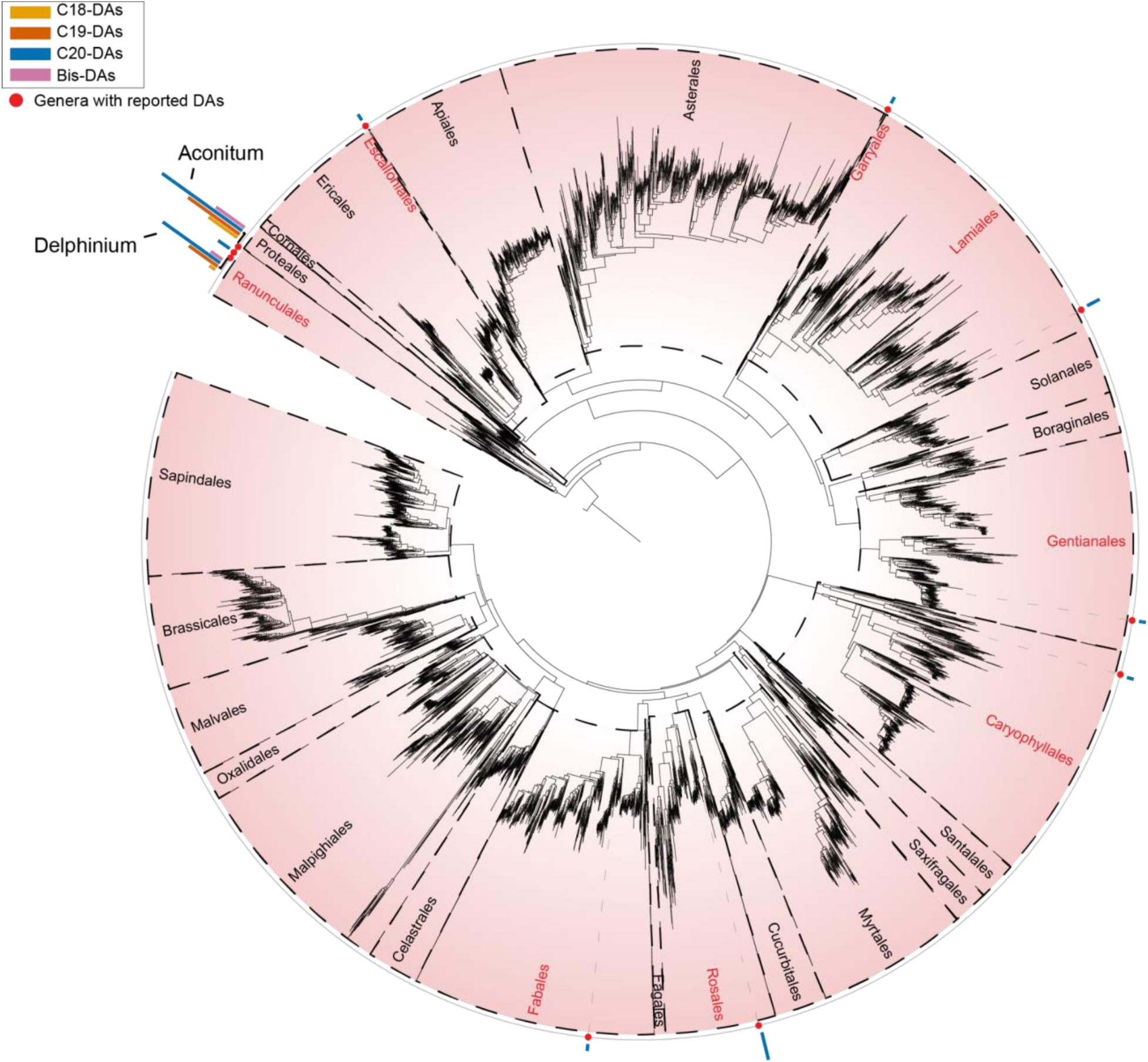
Angiosperm tree of life from Zuntini et al.^82^ with mapped occurrence of diterpenoid alkaloids. The mapping is based on the literature review, highlighting the largest diversity of DA scaffolds observed within the Ranunculaceae family. Plant orders in the tree are separated by black dashed lines, whereas each leaf in the tree corresponds to the representative species of the genus, as described in the original publication. The original tree can be accessed at https://itol.embl.de/tree/14723112167277531728383616.

**S. Figure 2.**
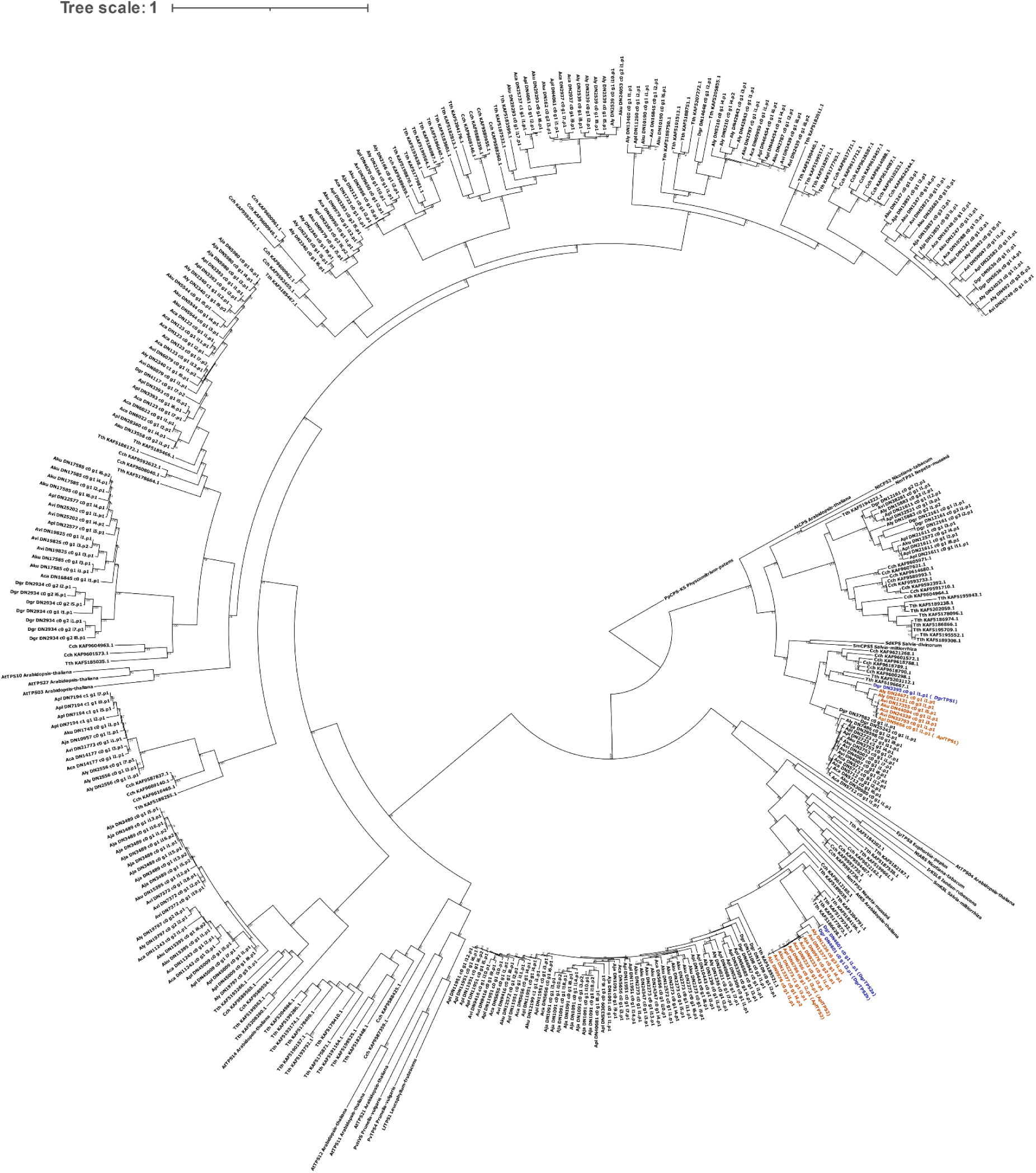
Maximum likelihood phylogenetic tree of candidate TPSs. This tree is an expanded view of that shown in Figure 2A. Branch lengths indicate substitutions per site and numbers at nodes represent percent support from 1,000 bootstrap replicates. A bifunctional *ent*-CPP/*ent*-kaurene synthase from *Physcomitrium patens* is used as an outgroup. Leaves in blue (*D. grandiflorum*) and orange (*Aconitum spp.*) represent sequences either characterized in this study or respective orthologs from other species. Sequences are from open reading frames of assembled transcripts and so duplicates or assembly artifacts may be present. Abbreviations: Tth: *Thalictrum thalictroides*; Cch: *Coptis chinensis*; Dgr: *Delphinium grandiflorum*; Apl: *Aconitum plicatum*; Aly: *Aconitum lycoctonum*; Aca: *Aconitum carmichaelii*; Aja: *Aconitum japonicum*; Avi: *Aconitum vilmorinianum*; Aku: *Aconitum kusnezoffii*.

**S. Figure 3.**
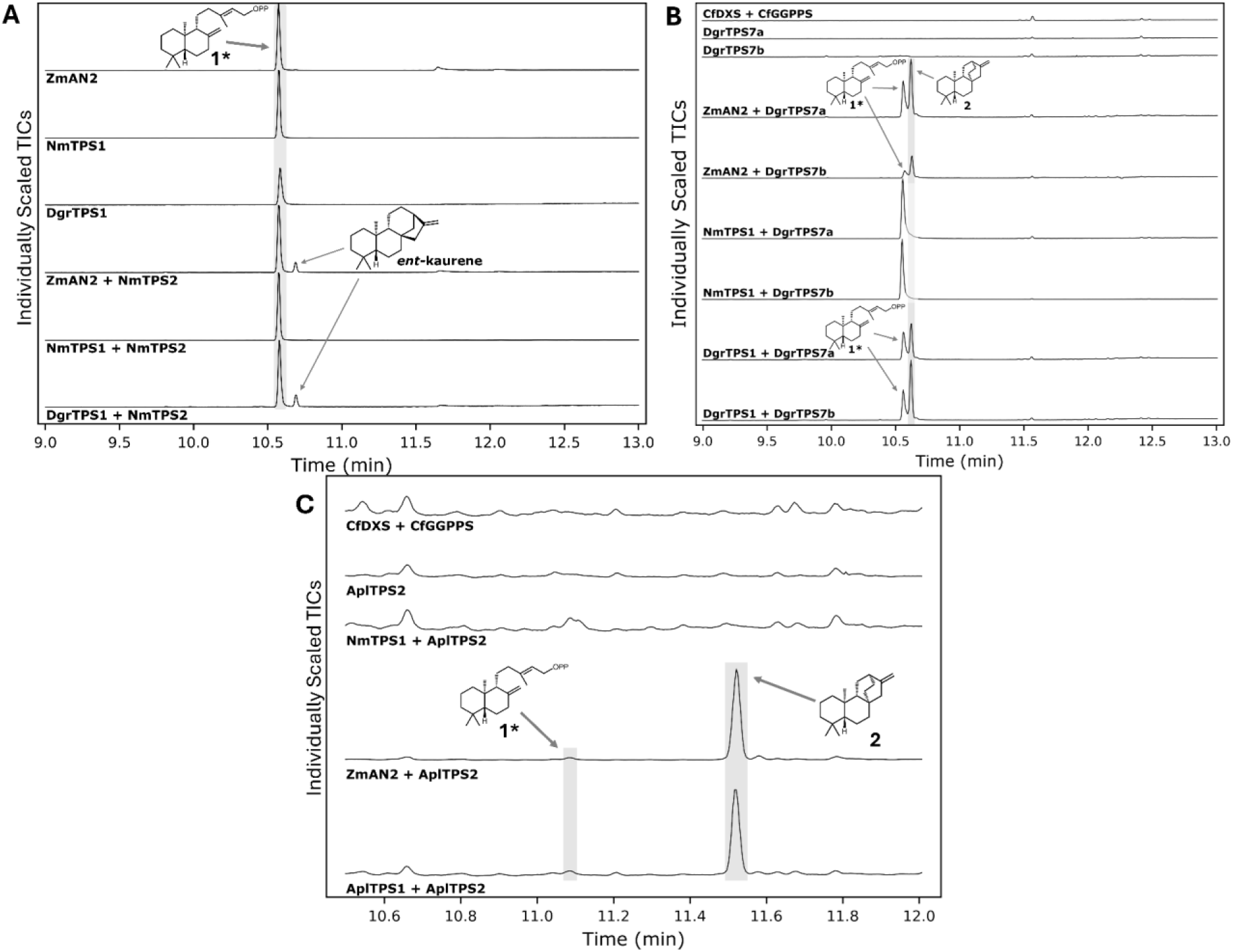
Function and conservation of TPSs. Each panel shows GC-MS chromatograms of hexane extracts of *N. benthamiana* infiltrations. Each assay includes *Cf*DXS and *Cf*GGPPS in addition to those listed. **A)** *Dgr*TPS1 functions as an *ent*-CPP synthase. *Zm*AN2 is a reference *ent*-CPP synthase, and *Nm*TPS1 forms (+)-CPP. **B)** Expanded view of Figure 2B showing the enantioselectivity of *Dgr*TPS2a/2b. **C**) Both *A. plicatum* TPS orthologs have the same function and enantioselectivity as *Dgr*TPS1 and *Dgr*TPS2. Retention times are not equal to other GC traces due to the use of a different GC method, as described in Methods. **1\***, the dephosphorylated derivative of ent-CPP (**1**).

**S. Figure 4:**
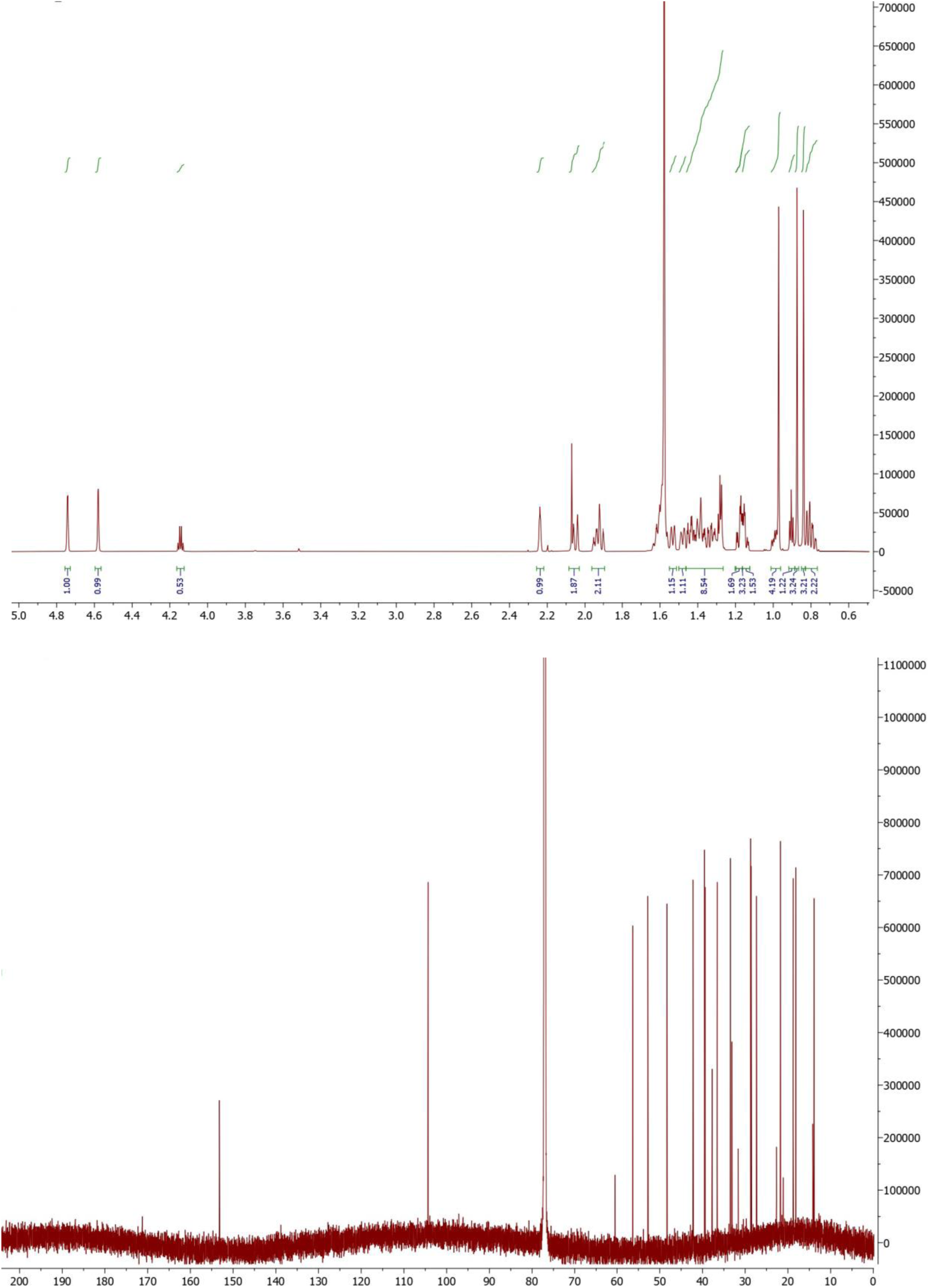
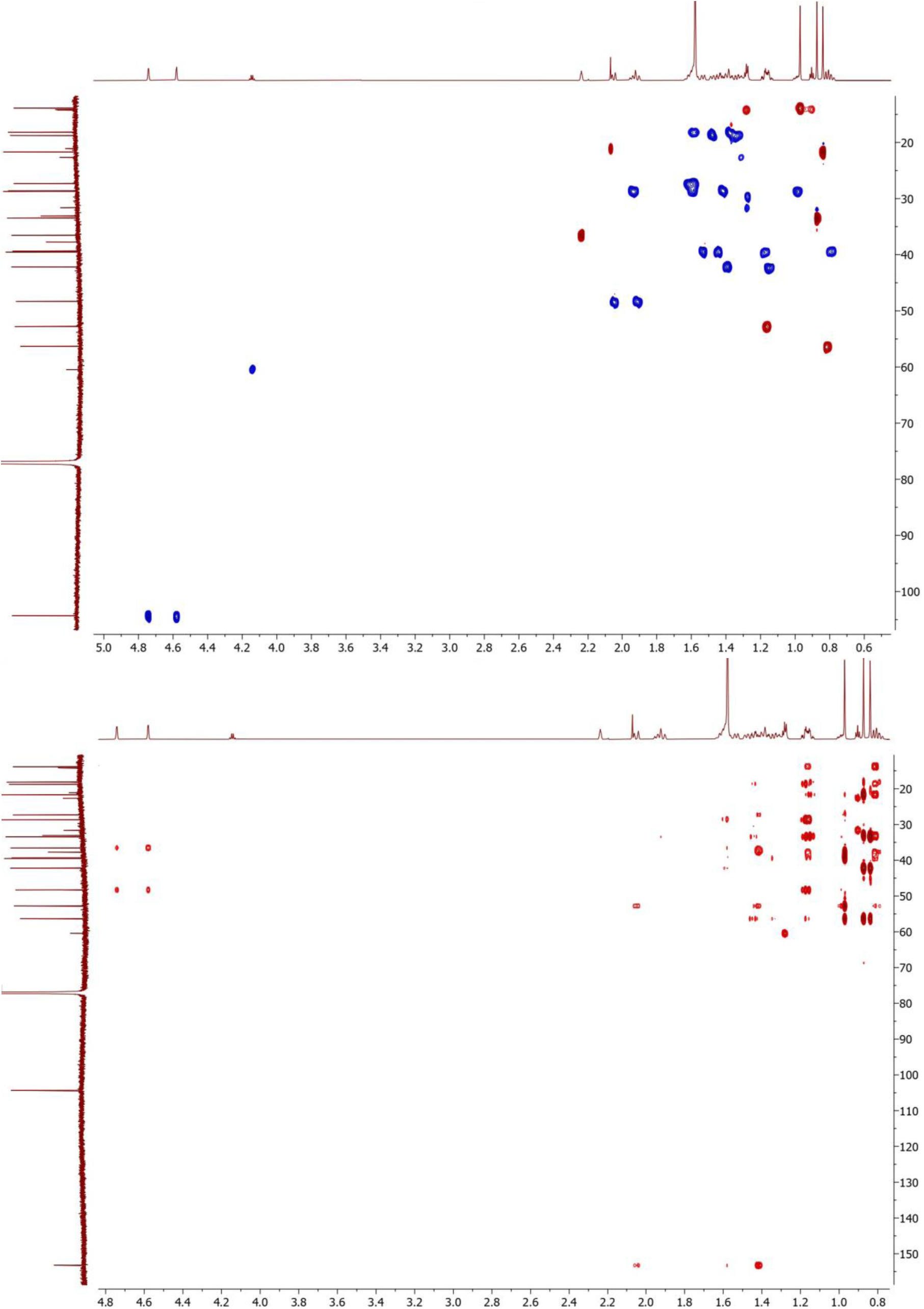
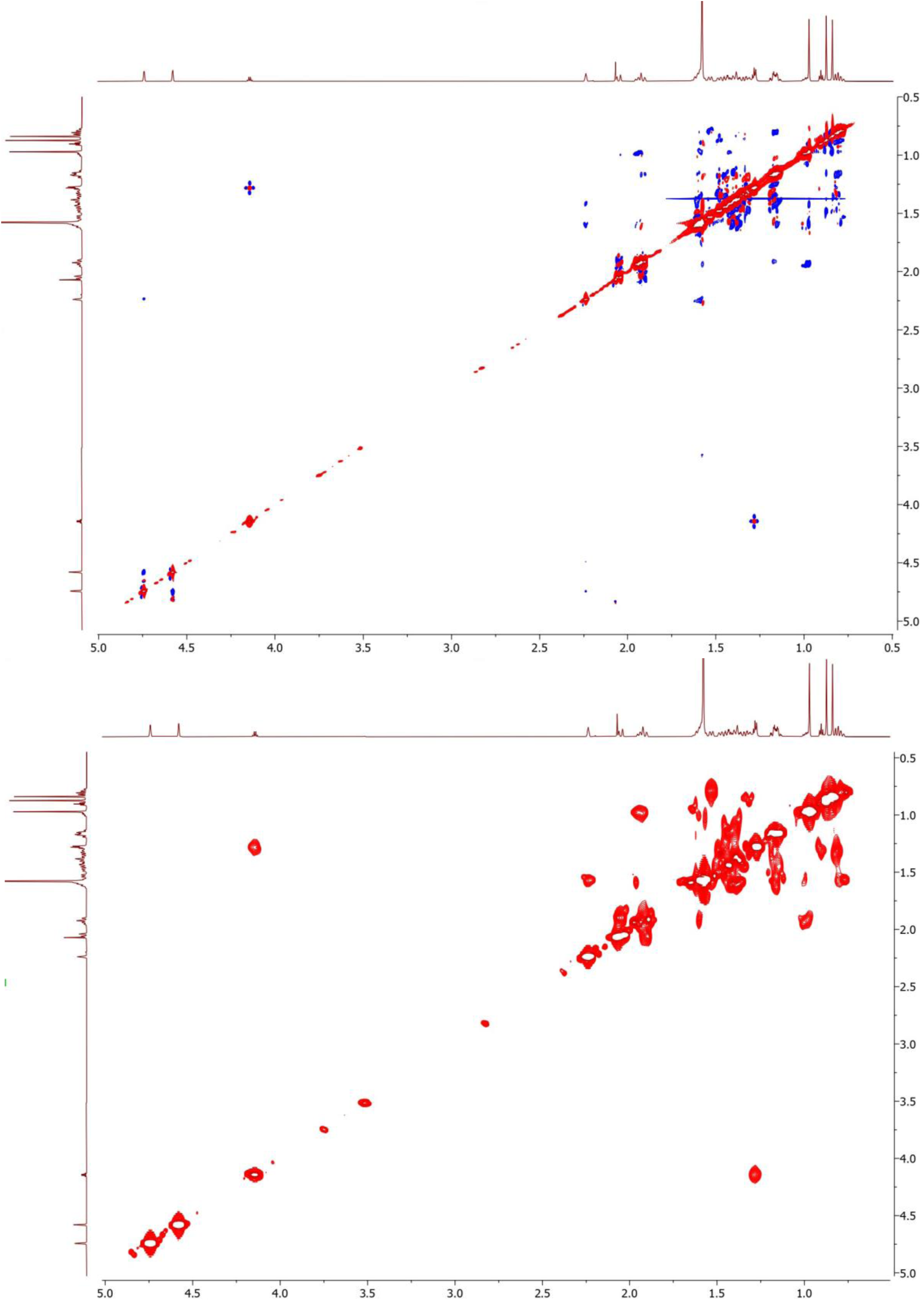
^1^H, ^13^C, HSQC, HMBC, COSY, and NOESY NMR spectra for ent-atiserene (**2**).

**S. Figure 5:**
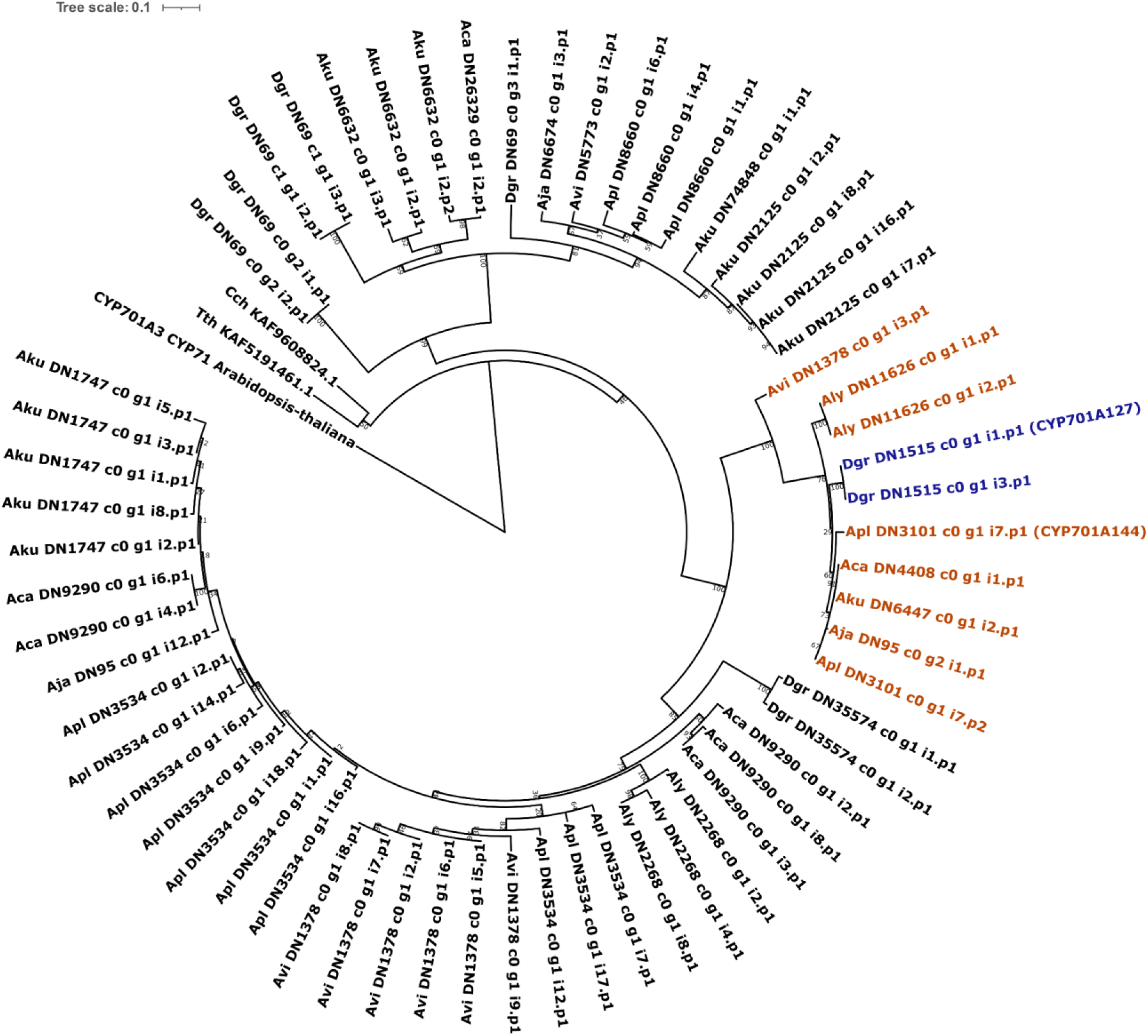
Maximum likelihood phylogenetic tree of candidate CYPs from the CYP701 family. This tree represents the second largest cluster within the CYP71 clan shown in the sequence similarity network in Figure 3A. Branch lengths indicate substitutions per site and numbers at nodes represent percent support from 1,000 bootstrap replicates. CYP701A3 from *Arabidopsis thaliana* is used as an outgroup. Leaves in blue (*D. grandiflorum*) and orange (*Aconitum spp.*) represent sequences either characterized in this study or respective orthologs from other species, containing CYP701A127 (*D. grandiflorum*) and CYP701A144 (*A. plicatum*). Sequences are from open reading frames of assembled transcripts and so duplicates or assembly artifacts may be present. Abbreviations: Tth: *Thalictrum thalictroides*; Cch: *Coptis chinensis*; Dgr: *Delphinium grandiflorum*; Apl: *Aconitum plicatum*; Aly: *Aconitum lycoctonum*; Aca: *Aconitum carmichaelii*; Aja: *Aconitum japonicum*; Avi: *Aconitum vilmorinianum*; Aku: *Aconitum kusnezoffii*.

**S. Figure 6:**
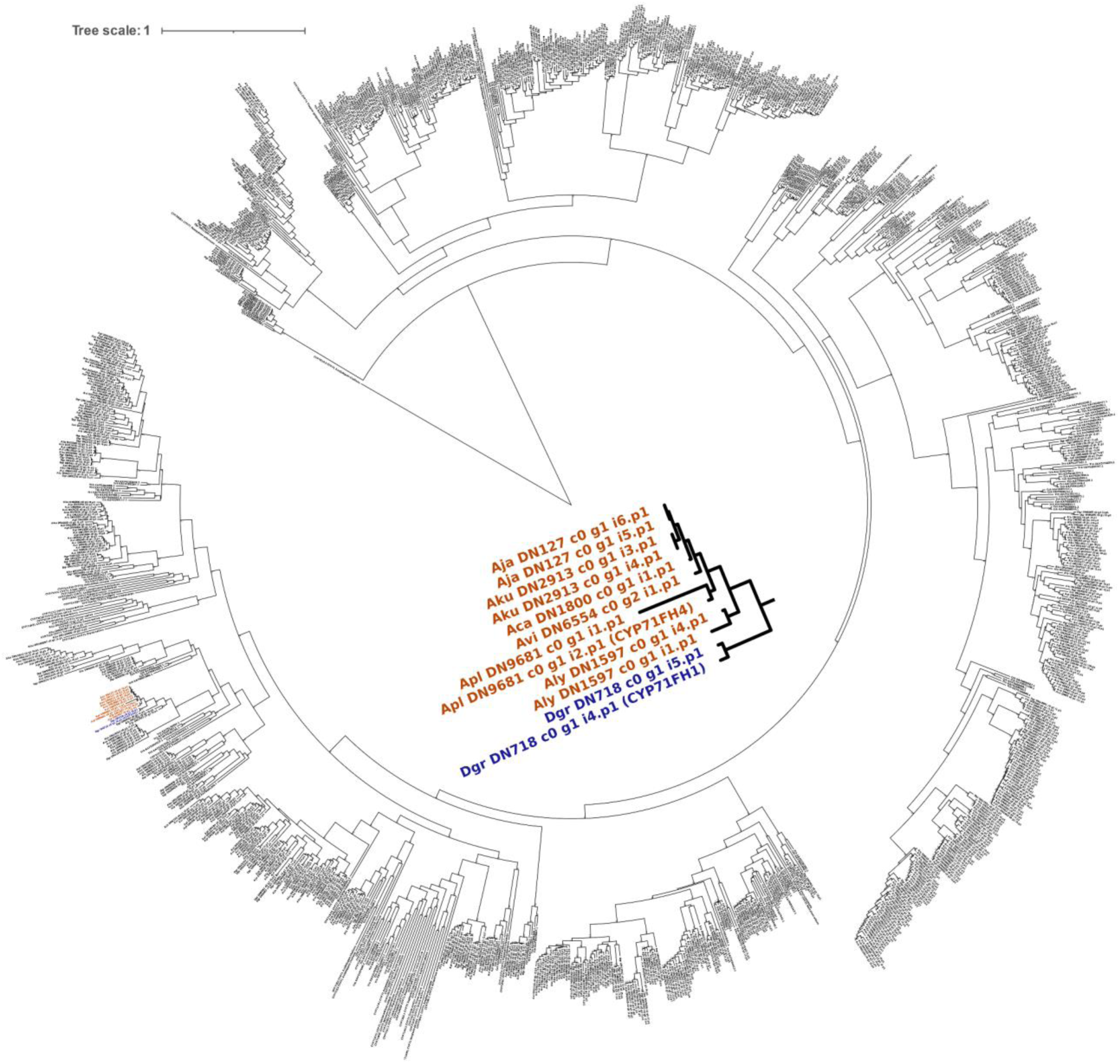
Maximum likelihood phylogenetic tree of candidate CYPs from the CYP71 clan. This tree represents the largest cluster within the CYP71 clan shown in the sequence similarity network in Figure 3A. Branch lengths indicate substitutions per site and numbers at nodes represent percent support from 100 bootstrap replicates. CYP701A3 from *Arabidopsis thaliana* is used as an outgroup. Leaves in blue (*D. grandiflorum*) and orange (*Aconitum spp.*) are enlarged in the center and represent sequences either characterized in this study or respective orthologs from other species and contains CYP71FH1 (*D. grandiflorum*) and CYP71FH4 (*A. plicatum*). Sequences are from open reading frames of assembled transcripts, and so duplicates or assembly artifacts may be present. Abbreviations: Tth: *Thalictrum thalictroides*; Cch: *Coptis chinensis*; Dgr: *Delphinium grandiflorum*; Apl: *Aconitum plicatum*; Aly: *Aconitum lycoctonum*; Aca: *Aconitum carmichaelii*; Aja: *Aconitum japonicum*; Avi: *Aconitum vilmorinianum*; Aku: *Aconitum kusnezoffii*.

**S. Figure 7:**
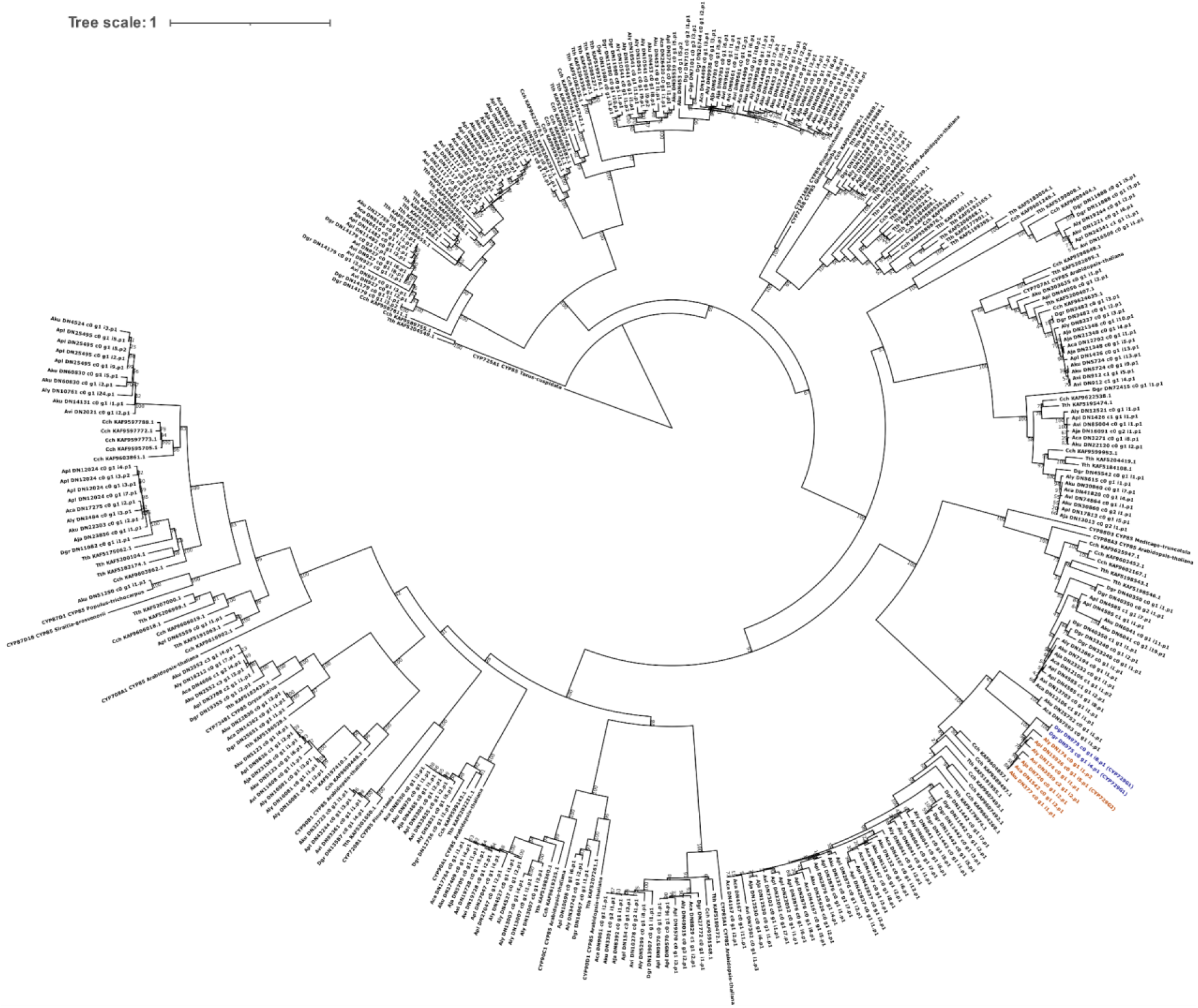
Maximum likelihood phylogenetic tree of candidate CYPs from the CYP85 clan. This is an expanded view of the tree shown in Figure 3A. Branch lengths indicate substitutions per site and numbers at nodes represent percent support from 1,000 bootstrap replicates. CYP701A3 from *Arabidopsis thaliana* is used as an outgroup. Leaves in blue (*D. grandiflorum*) and orange (*Aconitum spp.*) represent sequences either characterized in this study or respective orthologs from other species, containing CYP729G1 (*D. grandiflorum*) and CYP729G2 (*A. plicatum*). This highlighted region is enlarged in Figure 3A. Sequences are from open reading frames of assembled transcripts, and so duplicates or assembly artifacts may be present, and a duplicate transcript from *D. grandiflorum* was omitted for clarity within Figure 3A. Abbreviations: Tth: *Thalictrum thalictroides*; Cch: *Coptis chinensis*; Dgr: *Delphinium grandiflorum*; Apl: *Aconitum plicatum*; Aly: *Aconitum lycoctonum*; Aca: *Aconitum carmichaelii*; Aja: *Aconitum japonicum*; Avi: *Aconitum vilmorinianum*; Aku: *Aconitum kusnezoffii*.

**S. Figure 8:**
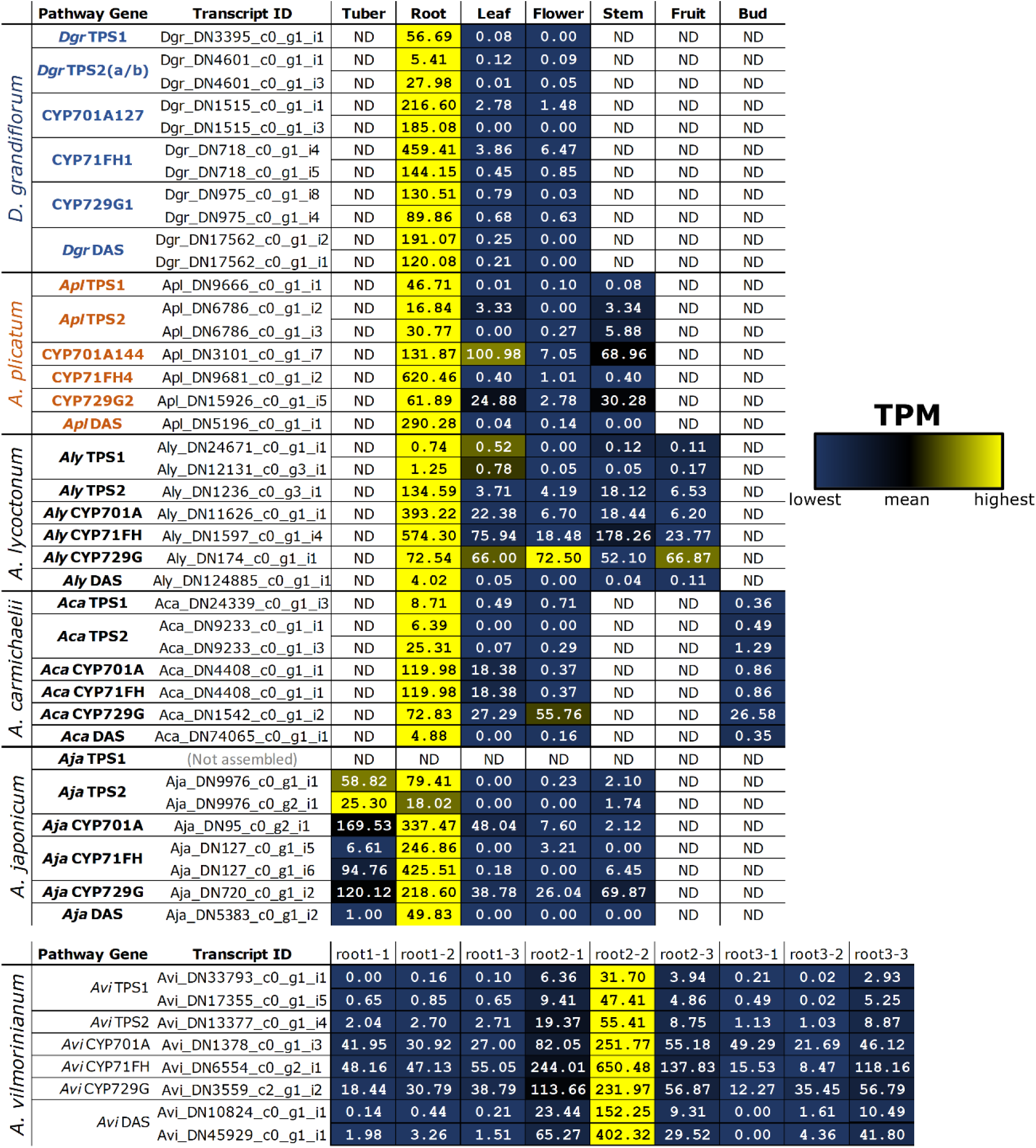
Calculated expression values of assembled transcripts associated with characterized enzymes, and their respective orthologs from other *Aconitum* species. Numbers represent transcripts per million (TPM) values within each respective assembly. Color scale is individualized to each row to highlight relative expression of each transcript between tissue types. Some characterized enzymes had multiple associated transcripts in transcriptome assemblies, and are shown here individually. All data shown above, and additional data for *A. kusnezoffii*, are given in Supplementary Dataset 1.

**S. Figure 9:**
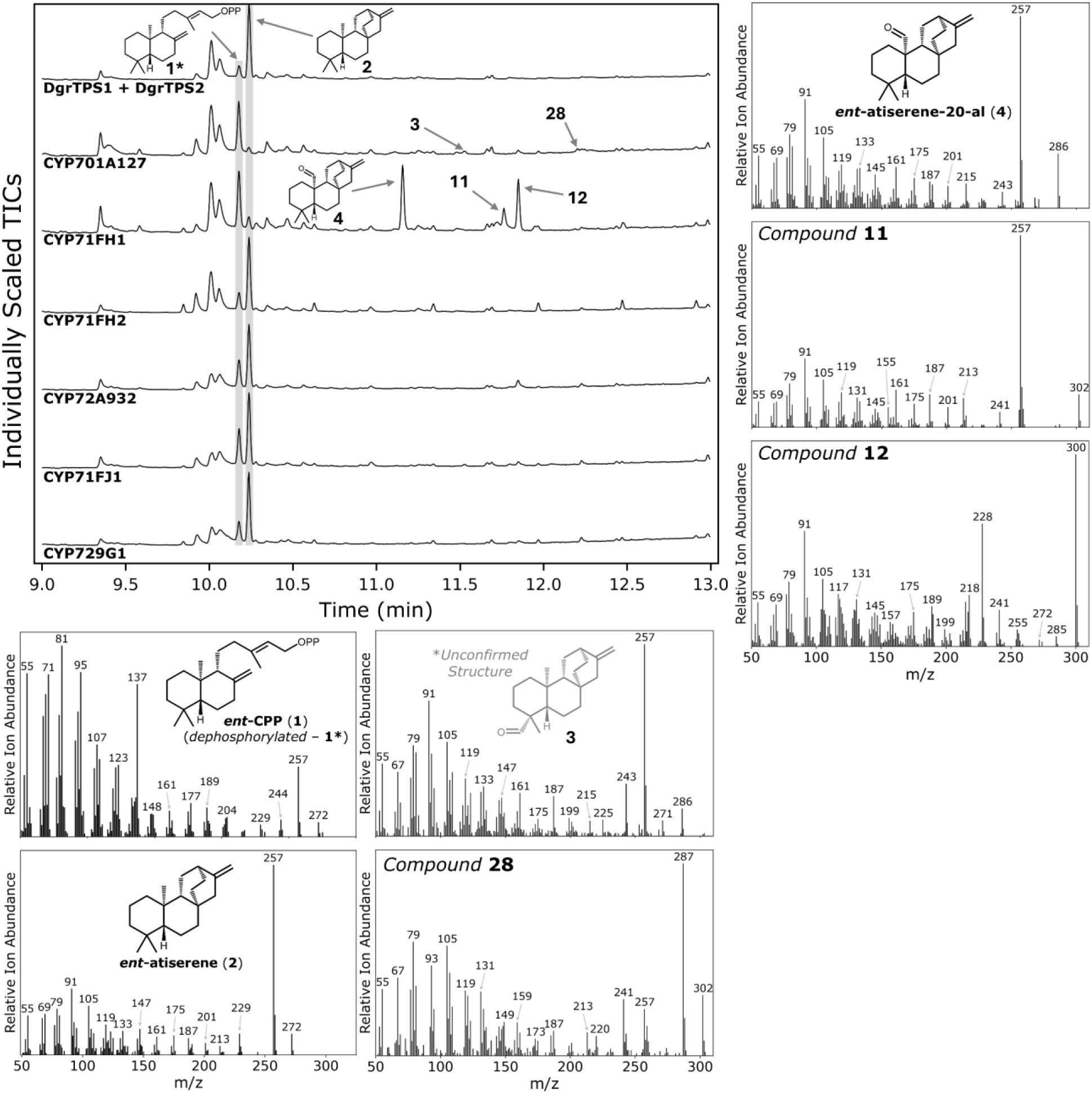
Initial screening of six CYPs with both TPSs. (Top Left) GC-MS chromatograms of EtOAc extracts of *N. benthamiana* infiltrations. Each assay includes *Cf*DXS, *Cf*GGPPS, *Dgr*TPS1, and *Dgr*TPS2 in addition to those listed. Compound **27** was not observed consistently across assays, likely due to low abundance. (Bottom Left) Mass spectra for compounds made through coexpression of *Dgr*TPS1 and *Dgr*TPS2. (Bottom Middle) Mass spectra for compounds made through coexpression of CYP701A127 with previous pathway steps. (Right) Mass spectra for compounds made through coexpression of CYP71FH1 with previous pathway steps.

**S. Figure 10:**
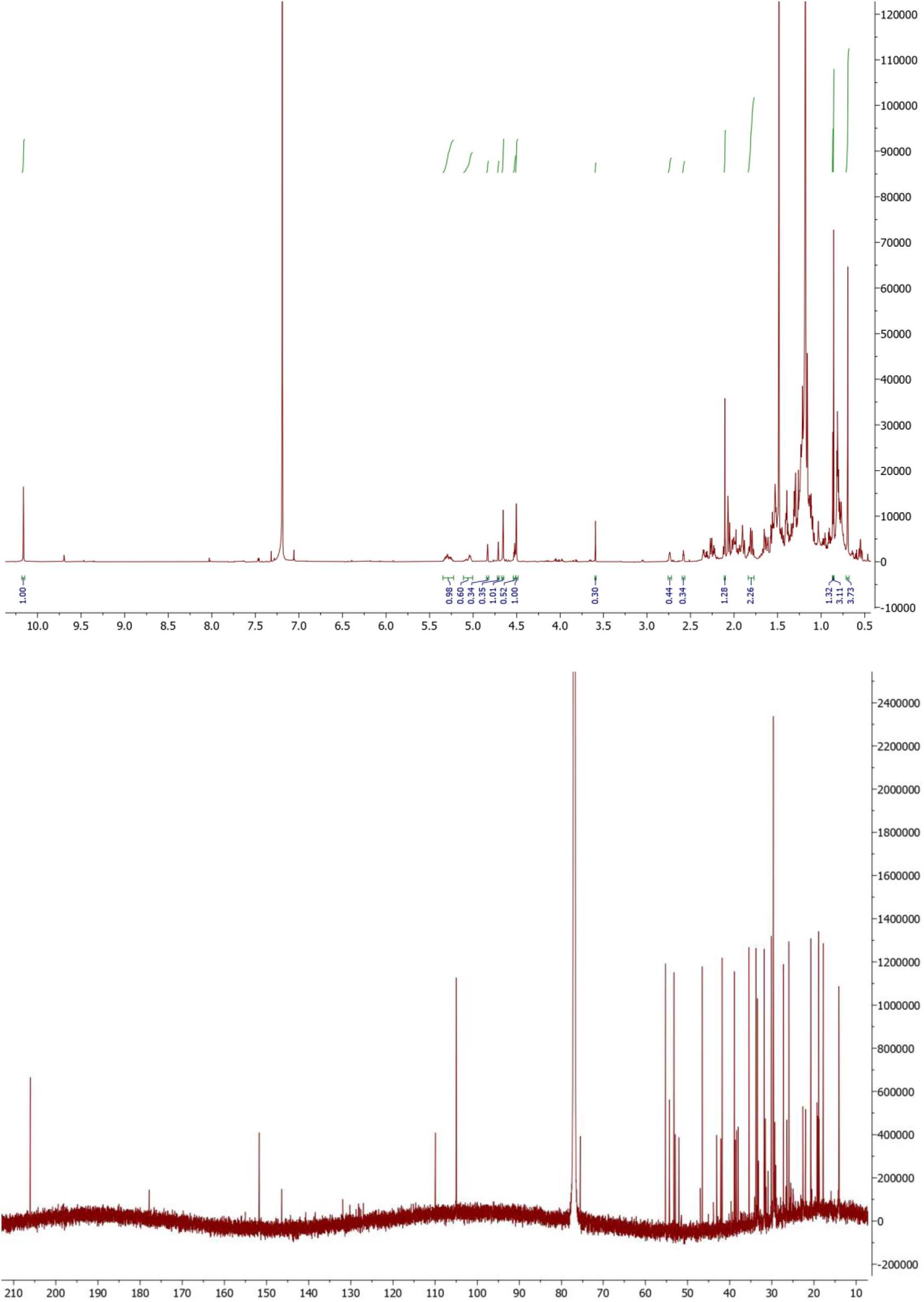
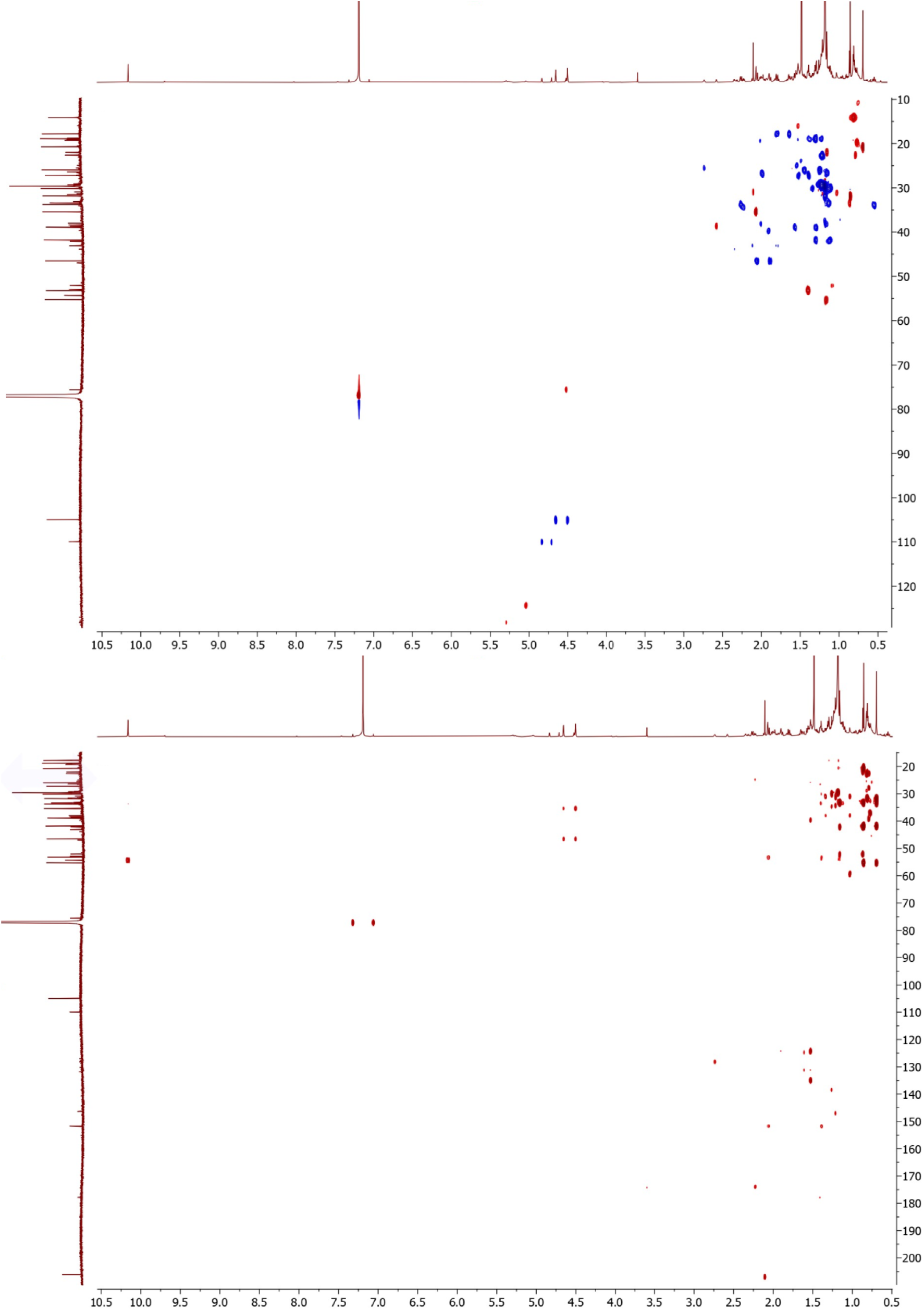
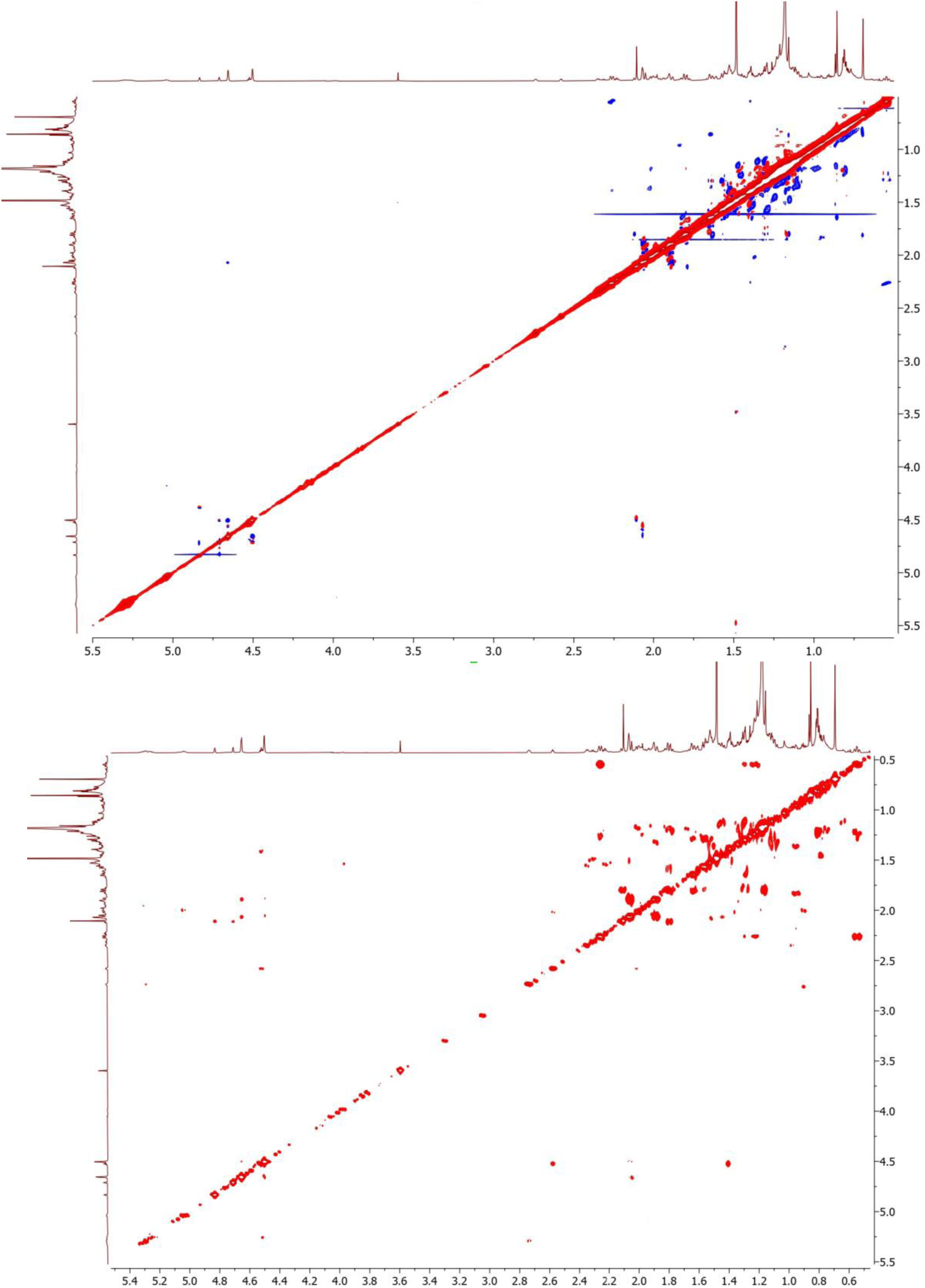
^1^H, ^13^C, HSQC, HMBC, NOESY, and COSY NMR spectra of ent-atiserene-20-al (**4**). Aldehyde peak is present in ^1^H spectrum at 10.16 ppm, which has the same integration value as terminal alkene protons (4.50 and 4.66 ppm). This product was not completely purified from **12** (peaks at 4.71 and 4.83 ppm are likely terminal alkene protons for **12**).

**S. Figure 11:**
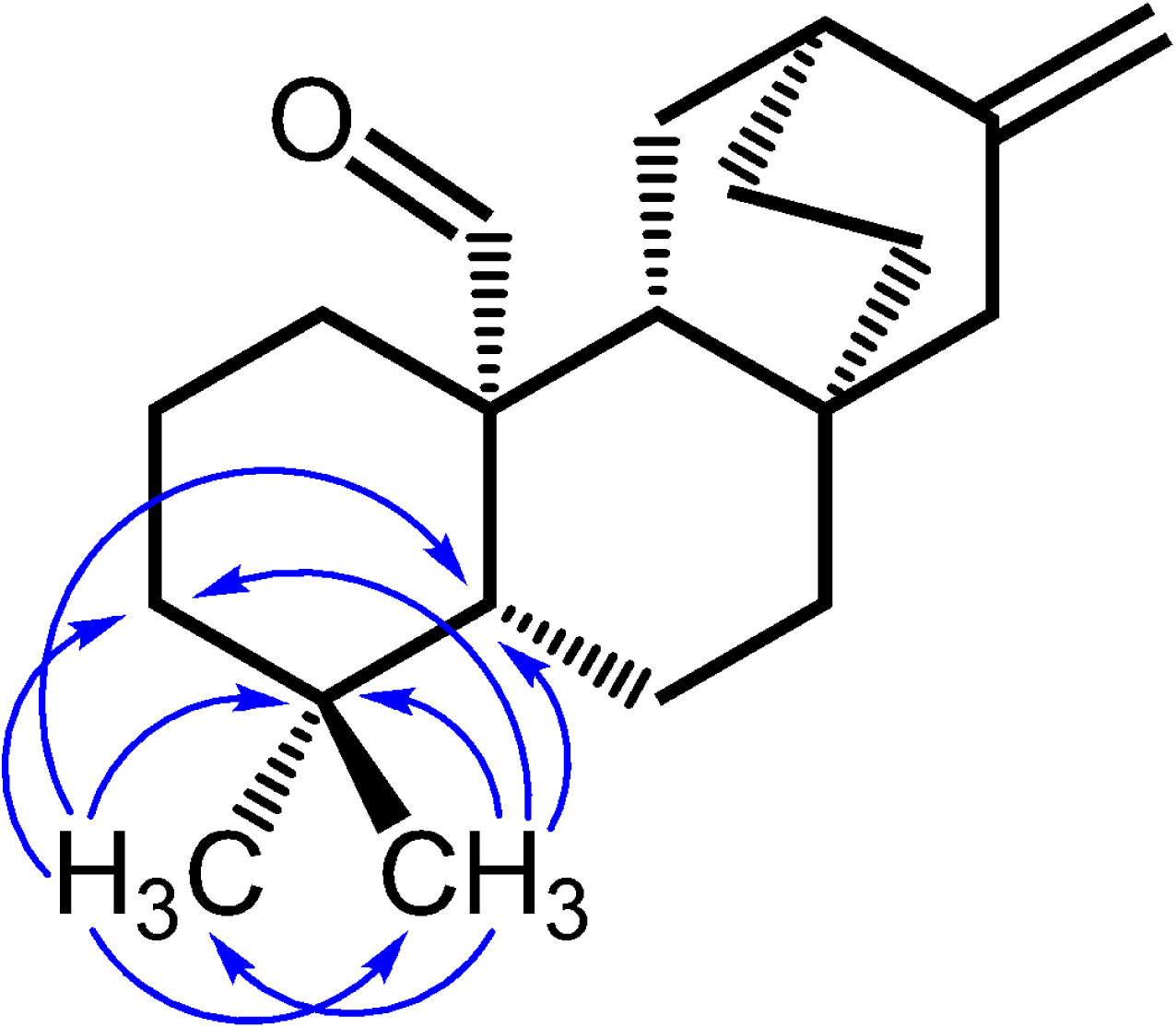
Select HMBC correlations for ent-atiserene-20-al (**4**). Correlations drawn show methyl groups for carbons 18 and 19 are retained following conversion of *ent*-atiserene (**2**) by CYP71FH1.

**S. Figure 12:**
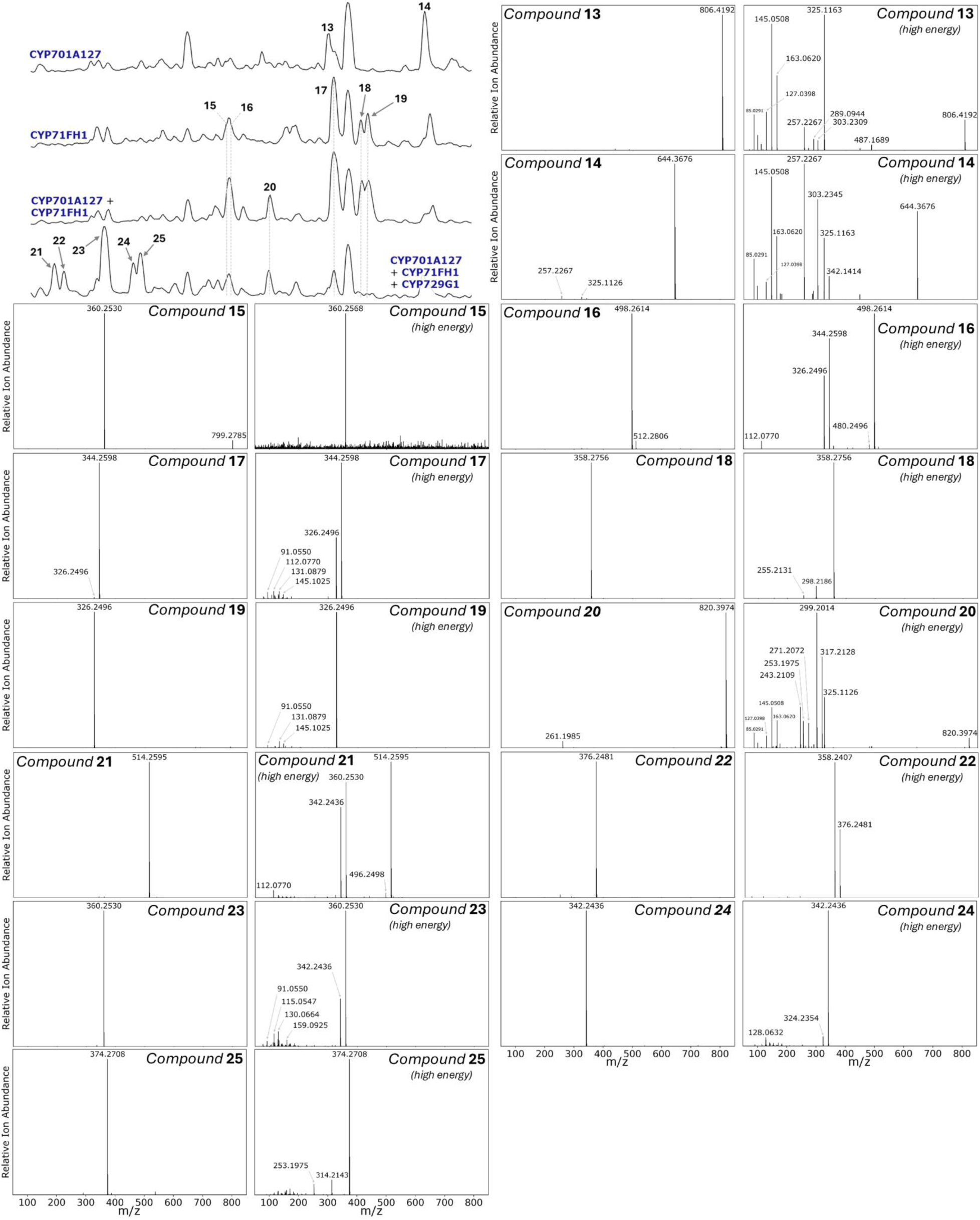
Mass spectra for all compounds shown in Figure 3C in the main text. Mass spectra were recorded in positive ion mode, and high energy spectra were generated with a voltage ramp from 20 to 80 V. Relevant portion of Figure 3C is shown in the top left.

**S. Figure 13:**
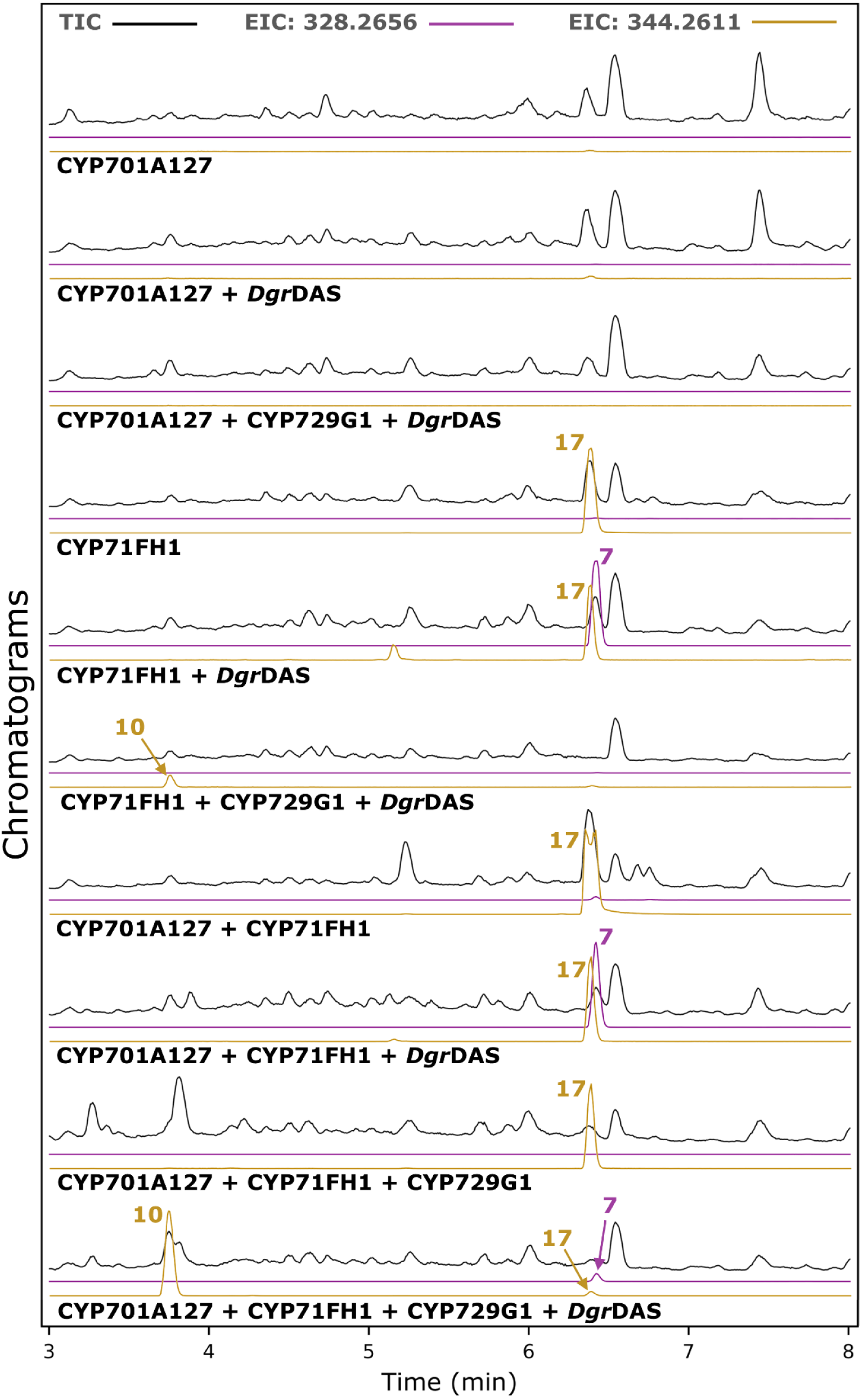
*CYP71FH1 complements a lack of CYP701A127 in atisinium (**10**) formation in* N. benthamiana *assays*. LC-MS chromatograms of 80% MeOH extracts of *N. benthamiana* infiltrations. Each assay includes *Cf*DXS, *Cf*GGPPS, *Dgr*TPS1, and *Dgr*TPS2 in addition to those listed.

**S. Figure 14:**
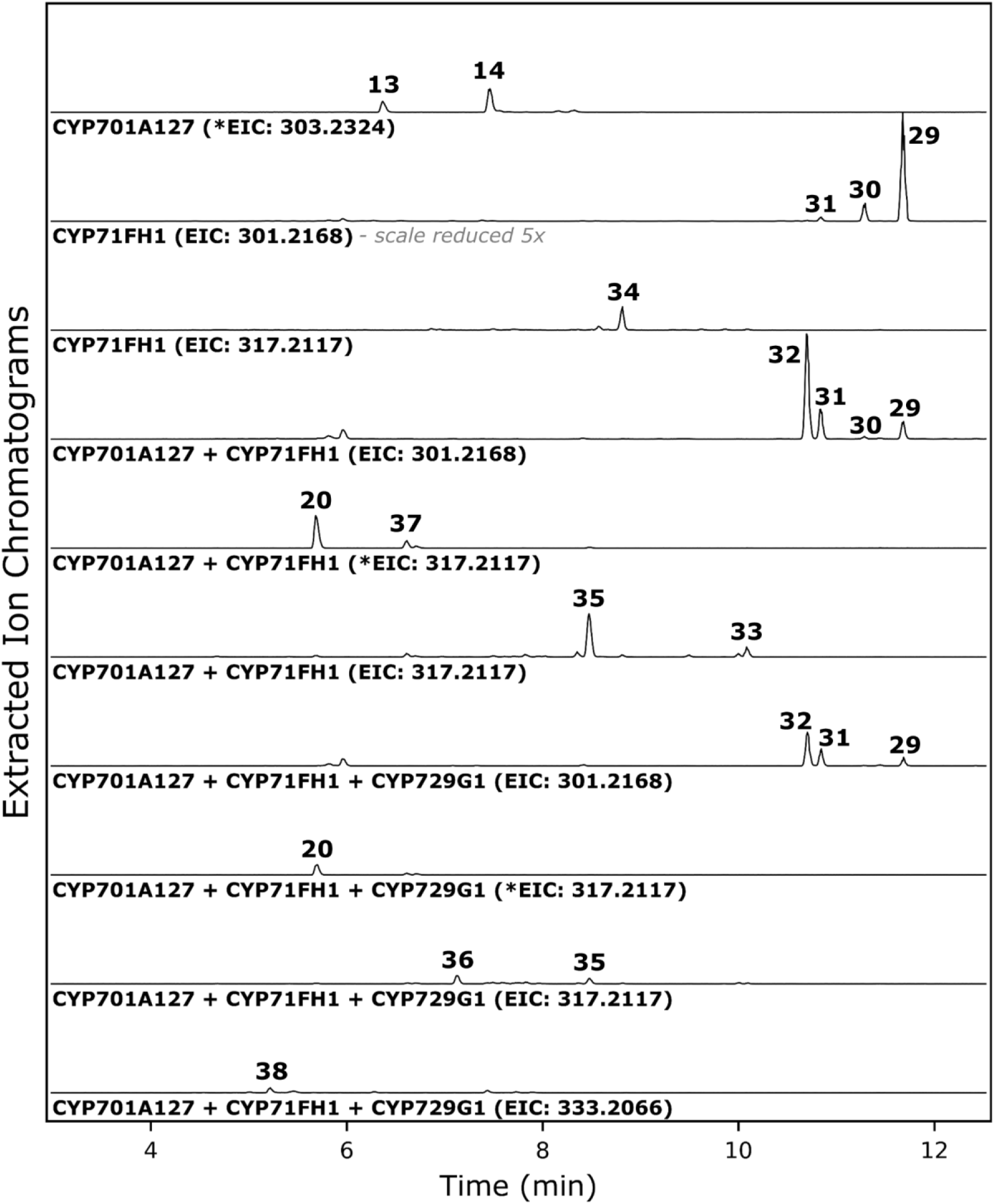
Extracted ion chromatograms in LC-MS analysis of CYP candidates. Extracted ions correspond to various predicted and confirmed masses for CYP products detected in 80% MeOH extracts from *N. benthamiana*. Chromatograms are scaled proportionally to each other except as otherwise indicated. EIC values with an asterisk indicate spectra obtained at a higher collision energy. Associated chemical formulas are given in S. Table 4. Compounds unique to this figure (numbered **29** or higher) are only visible by EIC and only accumulate in trace quantities.

**S. Figure 15:**
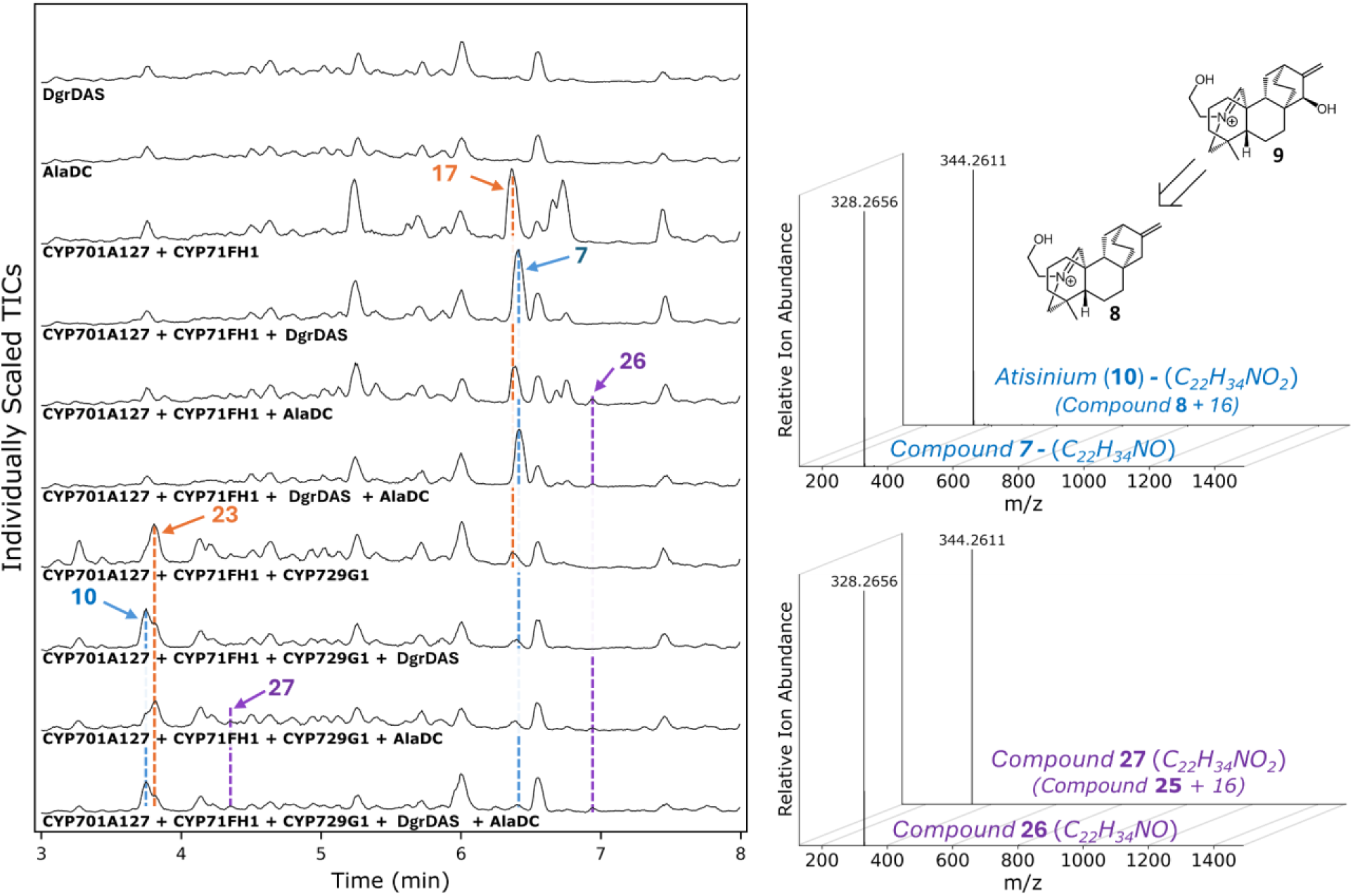
Initial testing to supplement the pathway with ethylamine through AlaDC. LC-MS chromatograms of 80% MeOH extracts of *N. benthamiana* infiltrations. Each assay includes *Cf*DXS, *Cf*GGPPS, *Dgr*TPS1, and *Dgr*TPS2 in addition to those listed. An alanine decarboxylase (AlaDC) was introduced to preceding pathway genes as an initial test to see how product profile changes with a presumed increase in supply of ethylamine. Only a trace amount of new product (**26**, or **27** with addition of CYP729G1) was observed, even when combined with *Dgr*DAS, and these products did not overlap with *Dgr*DAS’s products. Products traced in orange are products of CYPs alone, in blue are products that form with addition of *Dgr*DAS, and in purple are products that form with addition of AlaDC. Note that only select products are highlighted in this figure. A potential structure for **7** is drawn based on the difference in observed molecular weight of 16 *m/z* between atisinium (**10**) and **7**, and the conditions which result in this additional product involving an additional CYP (CYP729G1).

**S. Figure 16:**
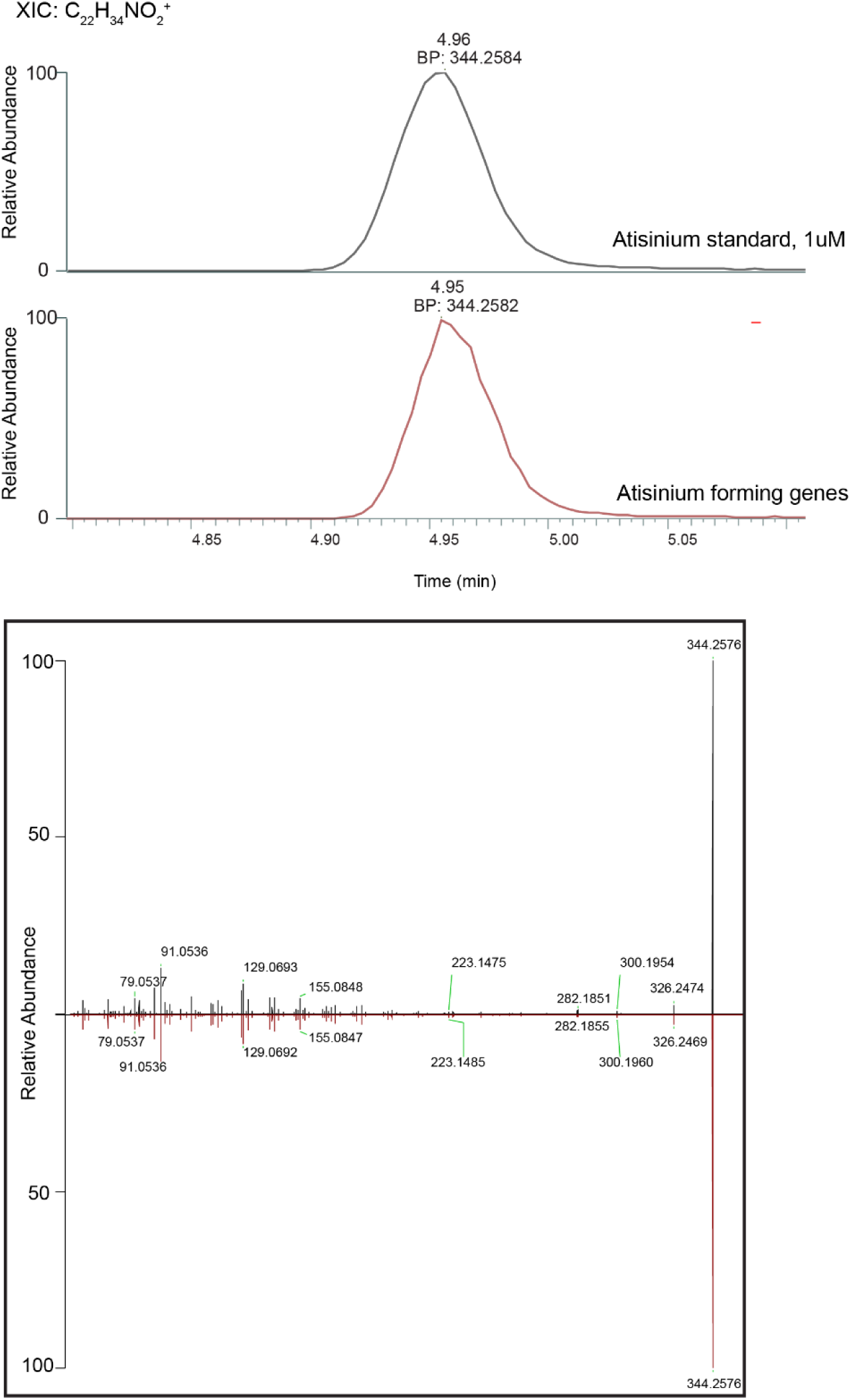
LC-MS data confirming atisinium formation. Atisinium formation resulting from infiltration of characterized genes in *N. benthamiana* (*Apl*TPS1, *Apl*TPS2, CYP71FH4, CYP701A144, CYP729G2, *Apl*DAS) was confirmed by comparison of retention times (upper panel) and MS/MS spectra (mirror plot, lower panel) with atisinium standard. MS/MS spectra were acquired at HCD 70. XIC, extracted ion chromatogram.

**S. Figure 17:**
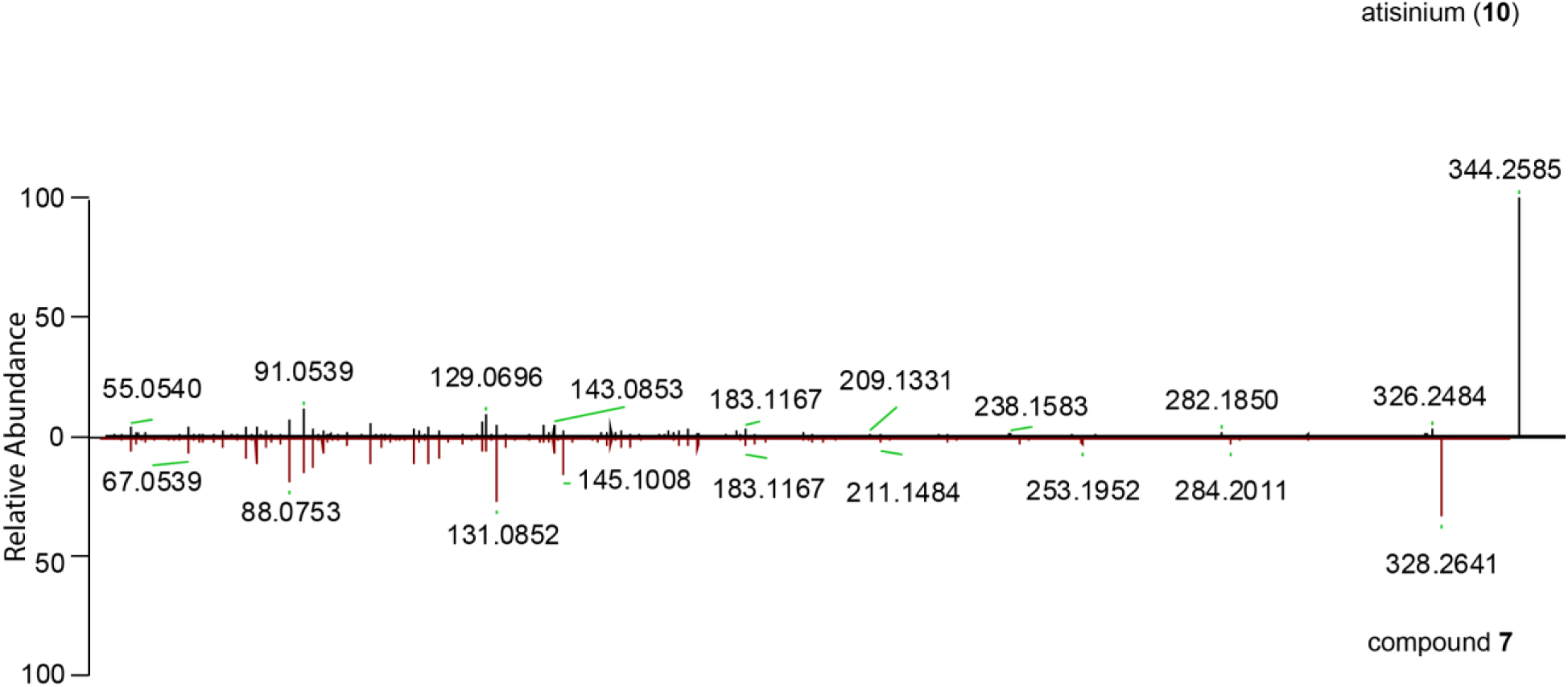
Mirror plot of MS/MS fragmentation spectra of atisinium and compound **7**. Highly similar fragmentation patterns of these compounds support their proposed structural similarity. Spectra for both compounds were acquired at HCD 70.

**S. Figure 18:**
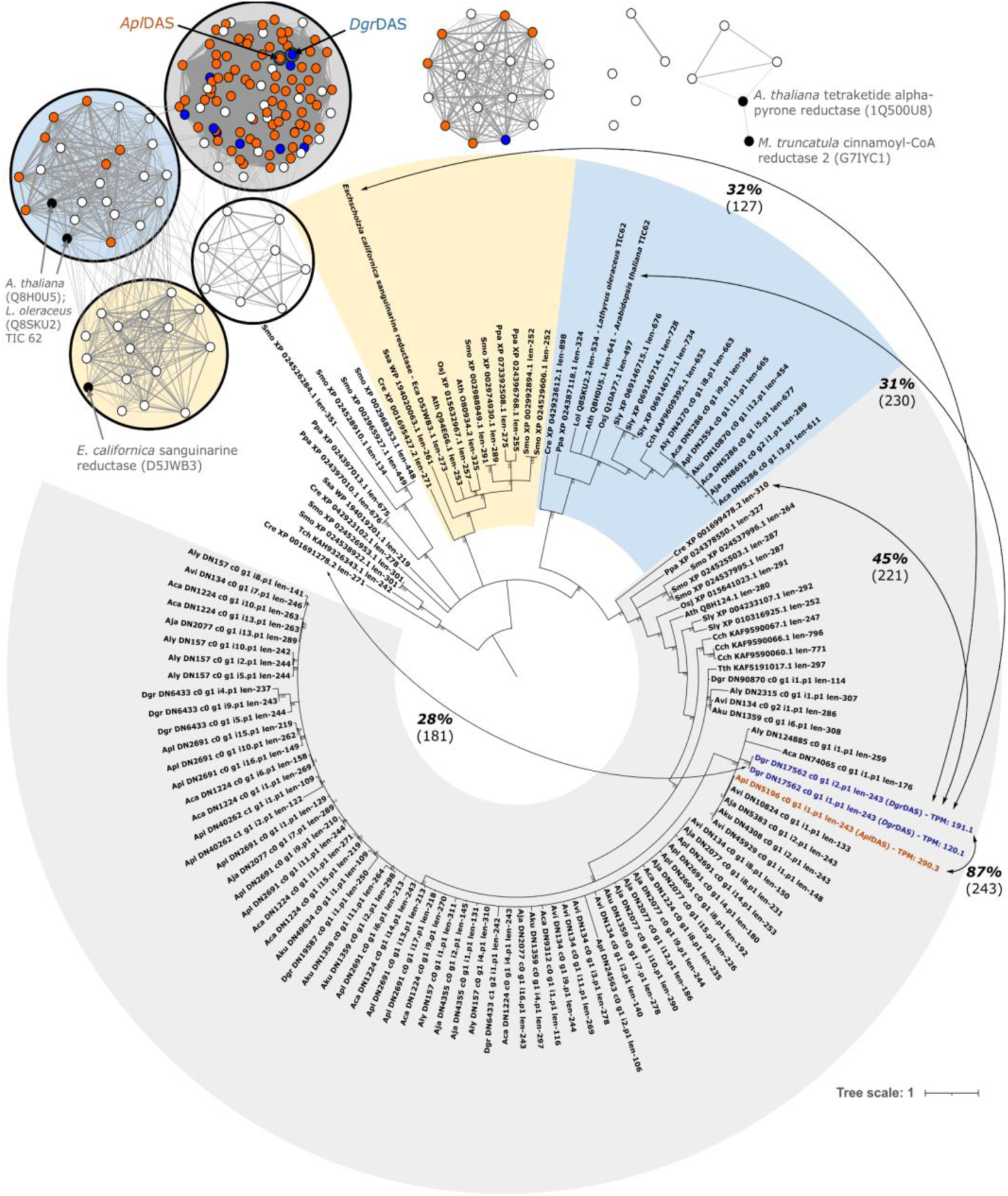
Dgr*DAS and* Apl*DAS similarity to other characterized sequences and those from model species*. Sequence similarity network of sequences matching *Dgr*DAS at a BLASTp cutoff of 1e-10, separated by 41% identity. *D. grandiflorum* sequences in blue, *Aconitum spp*. sequences in orange, characterized sequences in black, and other sequences from UniProt entries or model species in white. Circled clusters were included in the phylogenetic tree with corresponding highlighted branches. Branches with less than 50% bootstrap support have been collapsed. Select pairs of amino acid sequences with *Dgr*DAS have their percent identity annotated with alignment lengths in parentheses. The largest clade in gray includes sequences which are not currently characterized from a range of species, and their phylogenetic topology roughly matches the evolutionary relationship of these species (species abbreviations are given in S. Table 2). Within *Delphinium* and *Aconitum*, a large expansion of genes within this clade may be present, however many sequences presented here are likely misassembled (e.g. short or chimeric) due to the abundance of highly-similar sequences, which is also reflected in the poor bootstrapping support for branch topology within this section.

**S. Figure 19:**
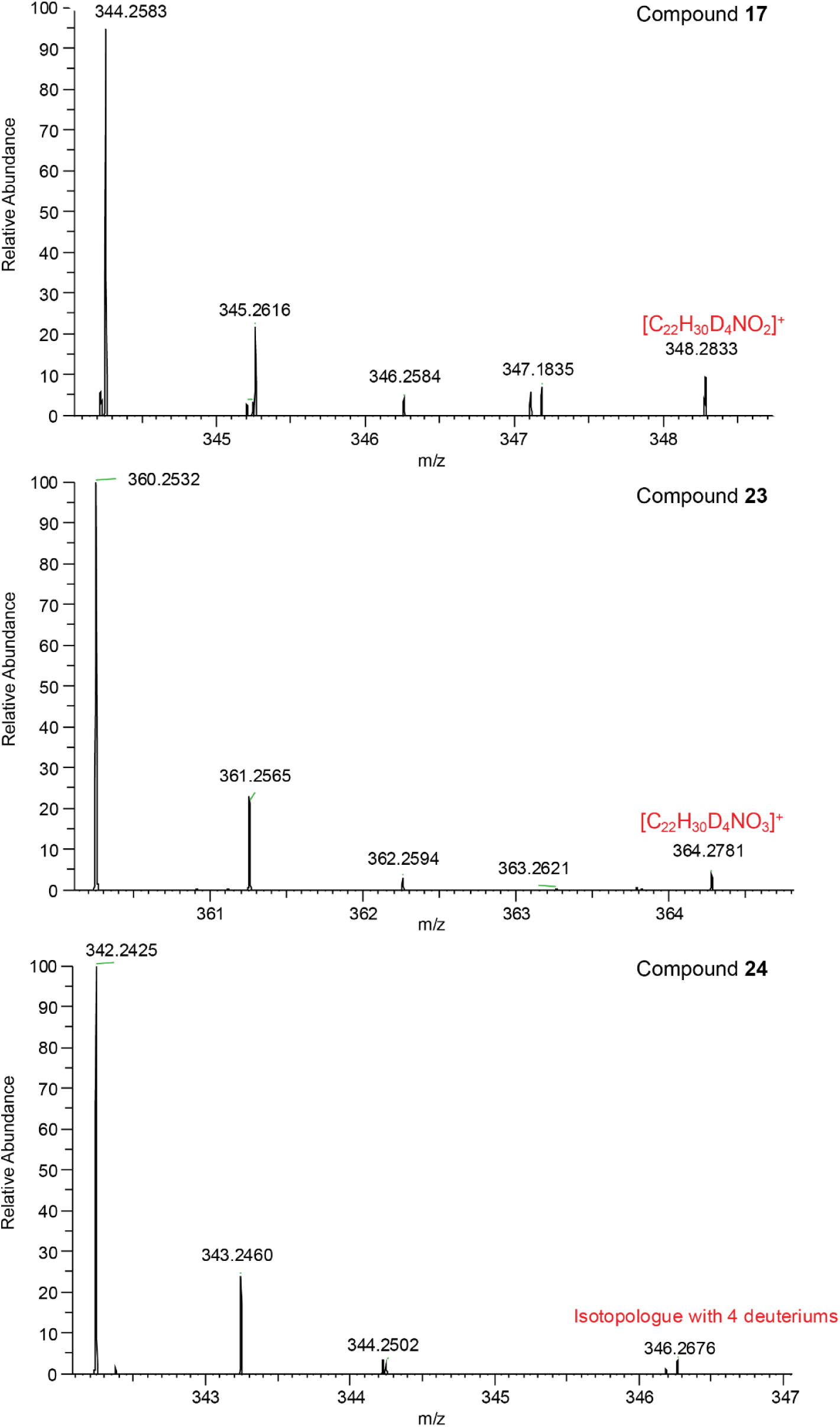
Isotope patterns of metabolites showing ethanolamine incorporation. The presence of highlighted isotopologues is consistent with labeled ethanolamine incorporation in *N. benthamiana* following infiltration with *Apl*TPS1, *Apl*TPS2, CYP701A144, CYP71FH4 and CYP729G2.

**S. Figure 20:**
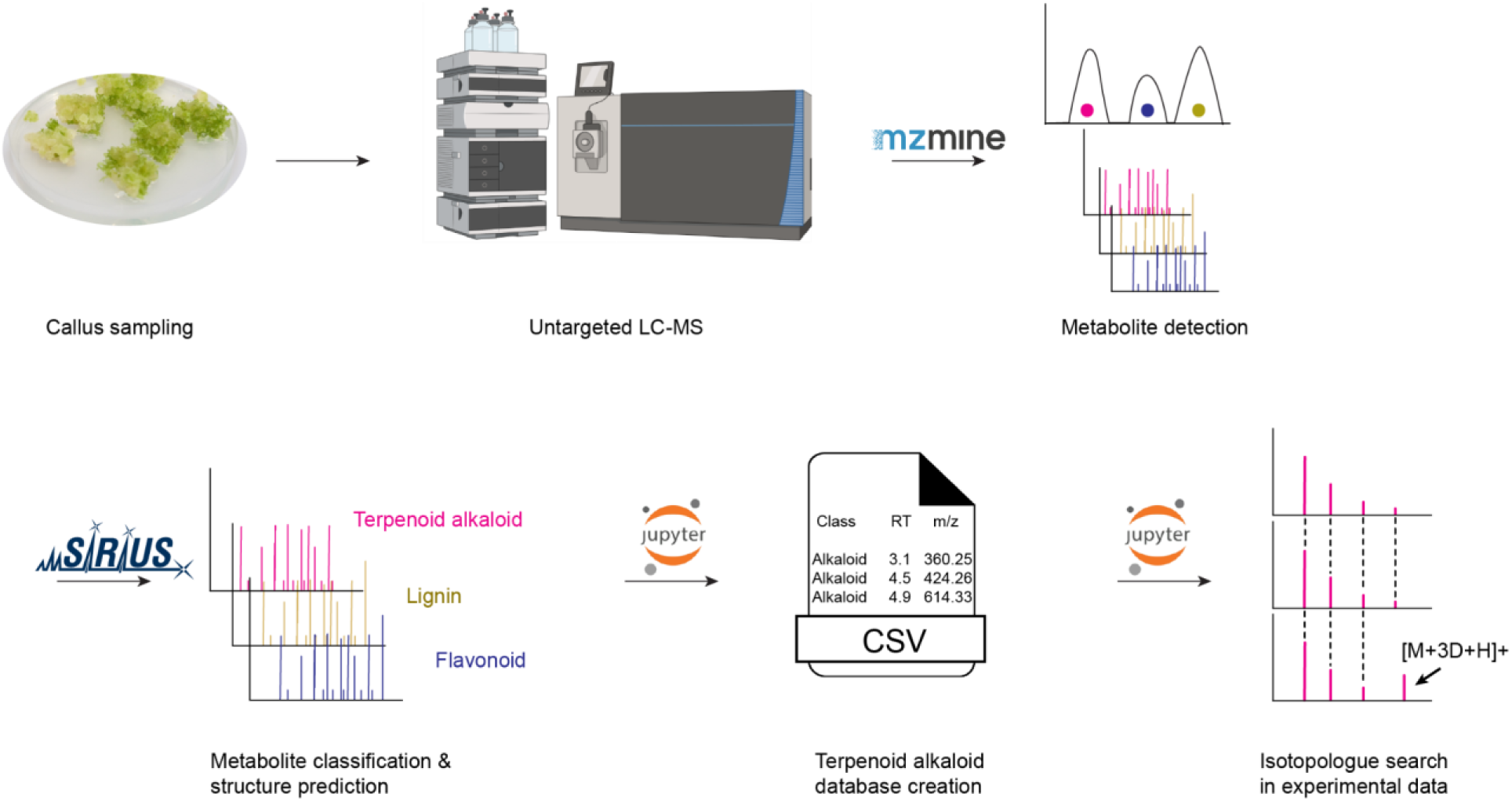
Workflow involved in identification of isotopologues from isotopically-labeled substrate feeding of A. plicatum callus cultures.

**S. Figure 21:**
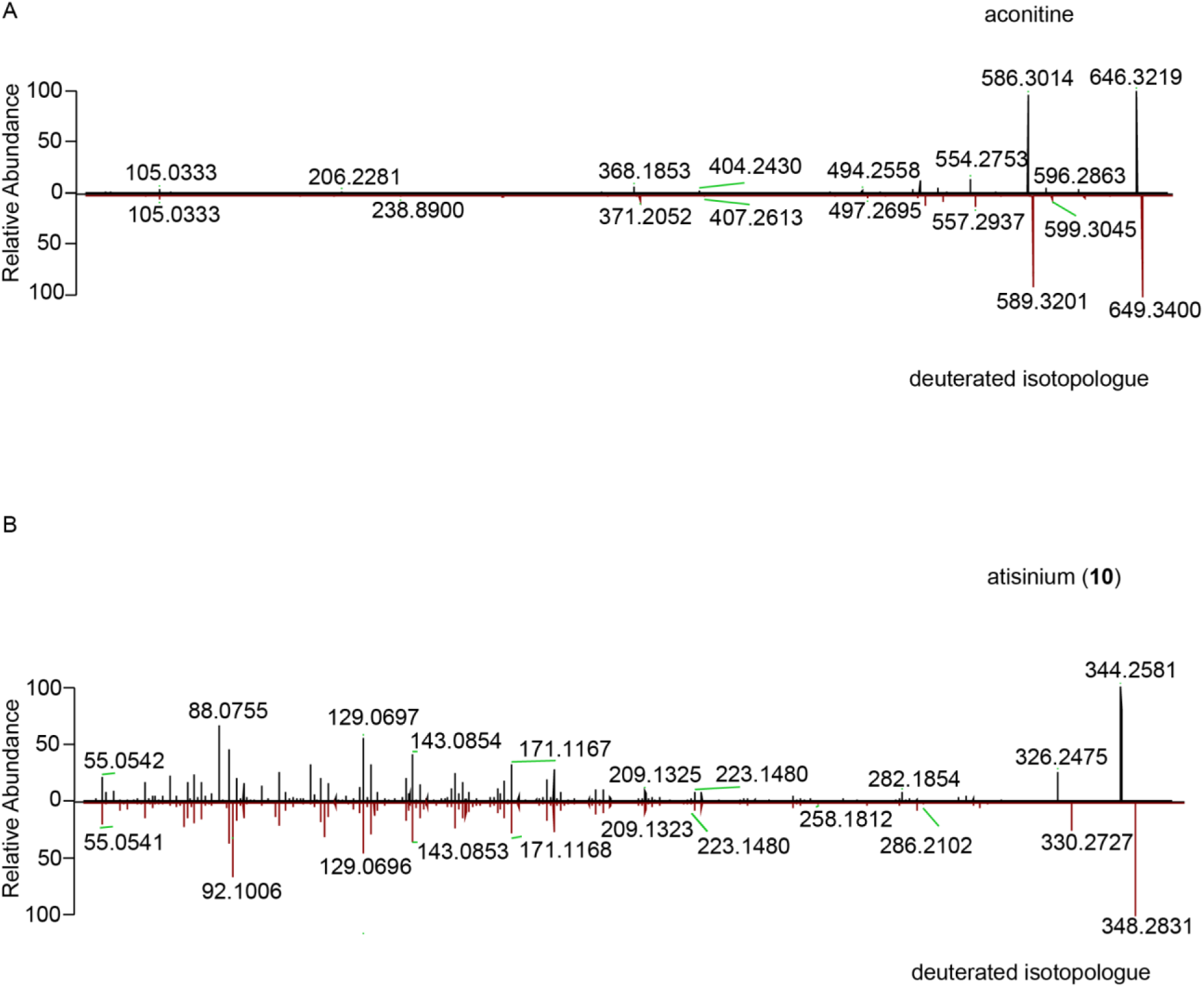
Mirror plots of aconitine, atisinium and their respective deuterated isotopologues demonstrating conserved fragmentation patterns. MS/MS fragmentation spectra of A) aconitine and its deuterated isotopologue at HCD 35 and B) atisinium and its deuterated isotopologue at HCD 70. The labeled compounds show the expected mass shift corresponding to incorporation of deuterium atoms while retaining comparable fragmentation patterns to the unlabeled compounds.

**S. Figure 22:**
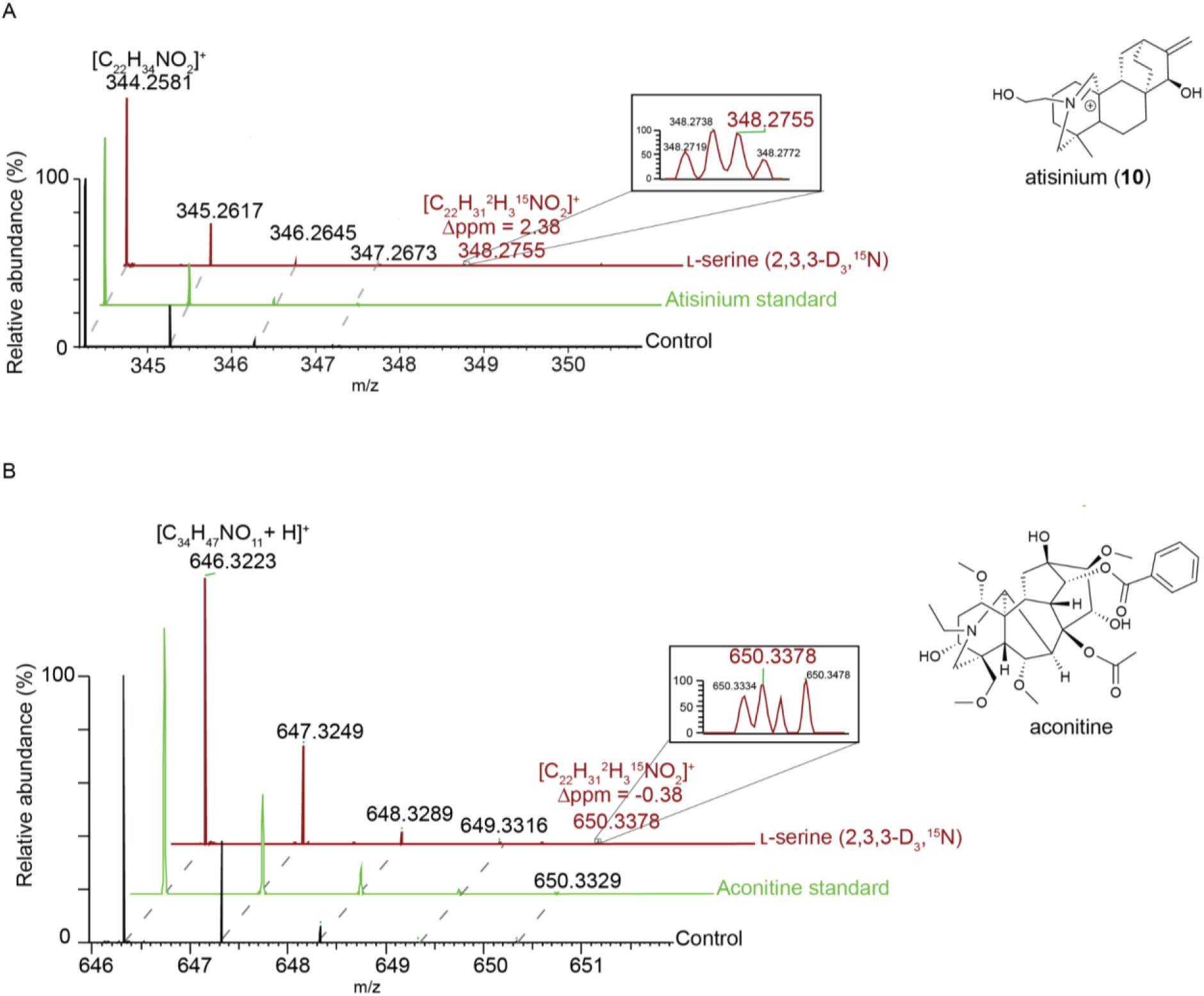
Tracing nitrogen incorporation in atisinium and aconitine. Isotope pattern of A) atisinium and B) aconitine in ʟ-serine (2,3,3-D3, ^15^N) fed callus. For each isotope pattern, the highlighted isotopologue corresponds to incorporation of three deuterium atoms and ^15^N.

**S. Table 1:** *^1^H and ^13^C chemical shifts for* ent*-atiserene (2)*. CDCl_3_ peaks were referenced to 7.26 and 77.00 ppm for ^1^H and ^13^C spectra, respectively.

| $^{13}\text{C}$ NMR | $\delta$ | $^1\text{H}$ NMR |
| --- | --- | --- |
| C1 | 39.4 | 0.79 (dd, $J = 3.7, 13.1$ Hz, 1H); 1.53 (m, 1H) |
| C2 | 18.2 | 1.38 (m, 1H); 1.59 (m, 1H) |
| C3 | 42.2 | 1.15 (m, 1H); 1.38 |
| C4 | 33.1 | - |
| C5 | 56.3 | 0.82 (dd, $J = 1.7, 12.1$ Hz, 1H) |
| C6 | 18.8 | 1.34 (m, 1H); 1.48 (ddt, $J = 13.3, 4.6, 2.4$ Hz, 1H) |
| C7 | 39.5 | 1.14 (m, 1H); 1.18 (m, 1H) |
| C8 | 33.5 | - |
| C9 | 52.8 | 1.16 (m, 1H) |
| C10 | 37.7 | - |
| C11 | 28.6 | 1.42 (m, 1H); 1.59 (m, 1H) |
| C12 | 36.6 | 2.24 (p, $J = 3.0$ Hz, 1H) |
| C13 | 28.7 | 0.99 (m, 1H); 1.94 (m, 1H) |
| C14 | 27.4 | 1.59 (m, 1H); 1.62 (m, 1H) |
| C15 | 48.3 | 1.91 (m, 1H); 2.05 (m, 1H) |
| C16 | 153.2 | - |
| C17 | 104.4 | 4.58 (q, $J = 2.1$ Hz, 1H); 4.74 (q, $J = 2.3$ Hz, 1H) |
| C18 | 33.5 | 0.87 (s, 3H) |
| C19 | 21.7 | 0.84 (s, 3H) |
| C20 | 13.9 | 0.98 (s, 3H) |

**S. Table 2:** List of species searched for sequences with homology to *Dgr*DAS and *Apl*DAS, with BioProject accessions associated with published genomes for model species. Three-letter abbreviations in parentheses correspond to those in phylogenetic trees. Select taxonomic classifications are included.

| Species | BioProject | Domain | Phylum | Class | Family |
| --- | --- | --- | --- | --- | --- |
| *Escherichia coli* (K-12) | PRJNA225 | Bacteria | Pseudomonadota | Gammaproteobacteria | Enterobacteriaceae |
| *Bacillus subtilis* | PRJNA76 | Bacteria | Bacillota | Bacilli | Bacillaceae |
| (Ssa) *Synechocystis salina* | PRJNA667061 | Bacteria | Cyanobacteriota | Cyanophyceae | Merismopediaceae |
| (Cre) *Chlamydomonas reinhardtii* | PRJNA12260 | Eukaryota | Chlorophyta | Chlorophyceae | Chlamydomonadaceae |
| (Ppa) *Physcomitrium patens* | PRJNA13064 | Eukaryota | Streptophyta | Bryopsida | Funariaceae |
| (Smo) *Selaginella moellendorffii* | PRJNA13078 | Eukaryota | Streptophyta | Lycopodiopsida | Selaginellaceae |
| (Tch) *Taxus chinensis* | PRJNA730337 | Eukaryota | Streptophyta | Pinopsida | Taxaceae |
| (Osj) *Oryza sativa japonica* | PRJNA953663 | Eukaryota | Streptophyta | Magnoliopsida | Poaceae |
| (Lol) *Lathyrus oleraceus* | (single sequence from UniProt) | Eukaryota | Streptophyta | Magnoliopsida | Fabaceae |
| (Ath) *Arabidopsis thaliana* | PRJNA10719 | Eukaryota | Streptophyta | Magnoliopsida | Brassicaceae |
| (Sly) *Solanum lycopersicum* | PRJDB16644 | Eukaryota | Streptophyta | Magnoliopsida | Solanaceae |
| (Eca) *Eschscholzia californica* | (single sequence from UniProt) | Eukaryota | Streptophyta | Magnoliopsida | Papaveraceae |
| (Tth) *Thalictrum thalictroides* | PRJNA439007 | Eukaryota | Streptophyta | Magnoliopsida | Ranunculaceae |
| (Cch) *Coptis chinensis* | PRJNA662860 | Eukaryota | Streptophyta | Magnoliopsida | Ranunculaceae |
| (Dgr) *Delphinium grandiflorum* | (transcriptome from this study) | Eukaryota | Streptophyta | Magnoliopsida | Ranunculaceae |
| (Aly) *Aconitum lycoctonum* | (transcriptome from this study) | Eukaryota | Streptophyta | Magnoliopsida | Ranunculaceae |
| (Apl) *Aconitum plicatum* | (transcriptome from this study) | Eukaryota | Streptophyta | Magnoliopsida | Ranunculaceae |
| (Avi) *Aconitum vilmorinianum* | (transcriptome from this study) | Eukaryota | Streptophyta | Magnoliopsida | Ranunculaceae |
| (Aca) *Aconitum carmichaelii* | (transcriptome from this study) | Eukaryota | Streptophyta | Magnoliopsida | Ranunculaceae |
| (Aja) *Aconitum japonicum* | (transcriptome from this study) | Eukaryota | Streptophyta | Magnoliopsida | Ranunculaceae |
| (Aku) *Aconitum kusnezoffii* | (transcriptome from this study) | Eukaryota | Streptophyta | Magnoliopsida | Ranunculaceae |
| *Homo sapiens* | PRJNA31257 | Eukaryota | Chordata | Mammalia | Hominidae |
| *Drosophila melanogaster* | PRJNA13812 | Eukaryota | Arthropoda | Insecta | Drosophilidae |
| *Saccharomyces cerevisiae* (S288C) | PRJNA43747 | Eukaryota | Ascomycota | Saccharomycetes | Saccharomycetaceae |
| *Cryptococcus neoformans* | PRJNA411 | Eukaryota | Basidiomycota | Tremellomycetes | Cryptococcaceae |

**S. Table 3:**
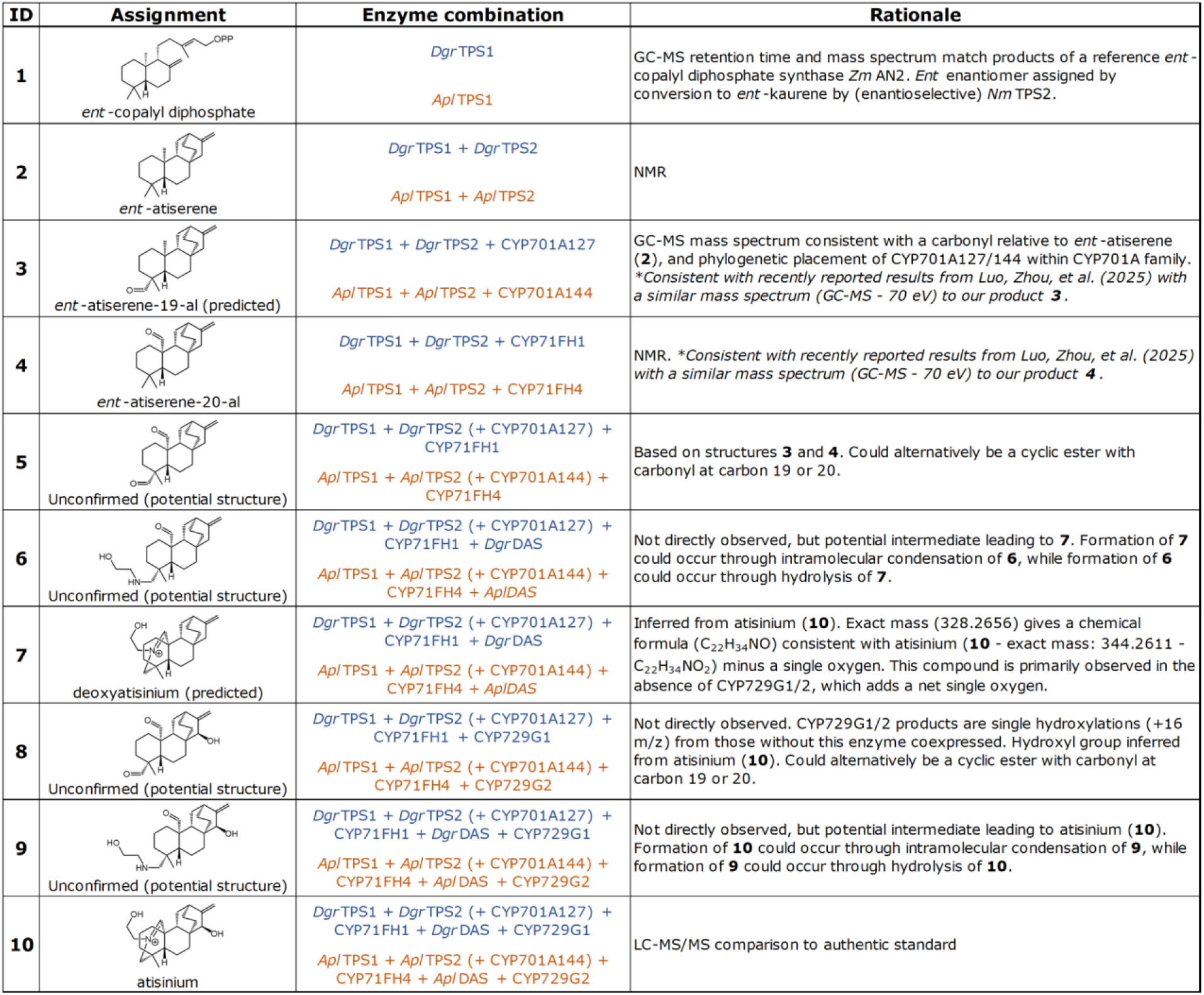
Compound numbering, structures, associated pathway enzymes, and rationale for confirmed or tentative structural assignments for all compounds drawn in Figure 1 in the main text.

| ID | Assignment | Enzyme combination | Rationale |
| --- | --- | --- | --- |
| 1 | ent-copalyl diphosphate | *Dgr*TPS1<br>*Apl*TPS1 | GC-MS retention time and mass spectrum match products of a reference *ent*-copalyl diphosphate synthase *Zm* AN2. *Ent* enantiomer assigned by conversion to *ent*-kaurene by (enantioselective) *Nm* TPS2. |
| 2 | ent-atiserene | *Dgr*TPS1 + *Dgr*TPS2<br>*Apl*TPS1 + *Apl*TPS2 | NMR |
| 3 | ent-atiserene-19-al (predicted) | *Dgr*TPS1 + *Dgr*TPS2 + CYP701A127<br>*Apl*TPS1 + *Apl*TPS2 + CYP701A144 | GC-MS mass spectrum consistent with a carbonyl relative to *ent*-atiserene (**2**), and phylogenetic placement of CYP701A127/144 within CYP701A family.<br>*Consistent with recently reported results from Luo, Zhou, et al. (2025) with a similar mass spectrum (GC-MS - 70 eV) to our product **3**. |
| 4 | ent-atiserene-20-al | *Dgr*TPS1 + *Dgr*TPS2 + CYP71FH1<br>*Apl*TPS1 + *Apl*TPS2 + CYP71FH4 | NMR. *Consistent with recently reported results from Luo, Zhou, et al. (2025) with a similar mass spectrum (GC-MS - 70 eV) to our product **4**. |
| 5 | Unconfirmed (potential structure) | *Dgr*TPS1 + *Dgr*TPS2 (+ CYP701A127) + CYP71FH1<br>*Apl*TPS1 + *Apl*TPS2 (+ CYP701A144) + CYP71FH4 | Based on structures **3** and **4**. Could alternatively be a cyclic ester with carbonyl at carbon 19 or 20. |
| 6 | Unconfirmed (potential structure) | *Dgr*TPS1 + *Dgr*TPS2 (+ CYP701A127) + CYP71FH1 + *Dgr*DAS<br>*Apl*TPS1 + *Apl*TPS2 (+ CYP701A144) + CYP71FH4 + *Apl*DAS | Not directly observed, but potential intermediate leading to **7**. Formation of **7** could occur through intramolecular condensation of **6**, while formation of **6** could occur through hydrolysis of **7**. |
| 7 | deoxyatisinium (predicted) | *Dgr*TPS1 + *Dgr*TPS2 (+ CYP701A127) + CYP71FH1 + *Dgr*DAS<br>*Apl*TPS1 + *Apl*TPS2 (+ CYP701A144) + CYP71FH4 + *Apl*DAS | Inferred from atisinium (**10**). Exact mass (328.2656) gives a chemical formula (C<sub>22</sub>H<sub>34</sub>NO) consistent with atisinium (**10** - exact mass: 344.2611 - C<sub>22</sub>H<sub>34</sub>NO<sub>2</sub>) minus a single oxygen. This compound is primarily observed in the absence of CYP729G1/2, which adds a net single oxygen. |
| 8 | Unconfirmed (potential structure) | *Dgr*TPS1 + *Dgr*TPS2 (+ CYP701A127) + CYP71FH1 + CYP729G1<br>*Apl*TPS1 + *Apl*TPS2 (+ CYP701A144) + CYP71FH4 + CYP729G2 | Not directly observed. CYP729G1/2 products are single hydroxylations (+16 m/z) from those without this enzyme coexpressed. Hydroxyl group inferred from atisinium (**10**). Could alternatively be a cyclic ester with carbonyl at carbon 19 or 20. |
| 9 | Unconfirmed (potential structure) | *Dgr*TPS1 + *Dgr*TPS2 (+ CYP701A127) + CYP71FH1 + *Dgr*DAS + CYP729G1<br>*Apl*TPS1 + *Apl*TPS2 (+ CYP701A144) + CYP71FH4 + *Apl*DAS + CYP729G2 | Not directly observed, but potential intermediate leading to atisinium (**10**). Formation of **10** could occur through intramolecular condensation of **9**, while formation of **9** could occur through hydrolysis of **10**. |
| 10 | atisinium | *Dgr*TPS1 + *Dgr*TPS2 (+ CYP701A127) + CYP71FH1 + *Dgr*DAS + CYP729G1<br>*Apl*TPS1 + *Apl*TPS2 (+ CYP701A144) + CYP71FH4 + *Apl*DAS + CYP729G2 | LC-MS/MS comparison to authentic standard |

**S. Table 4:**
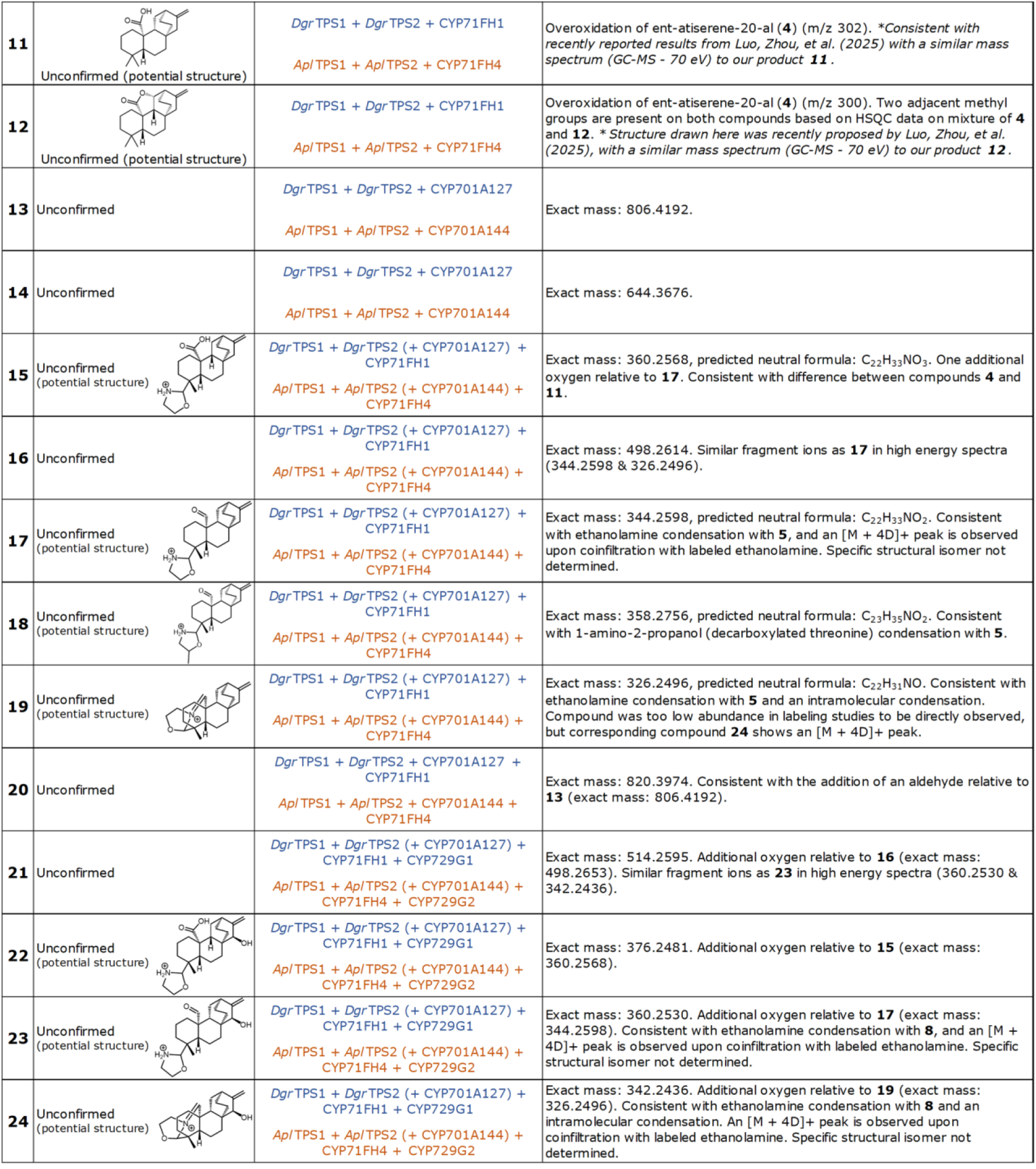
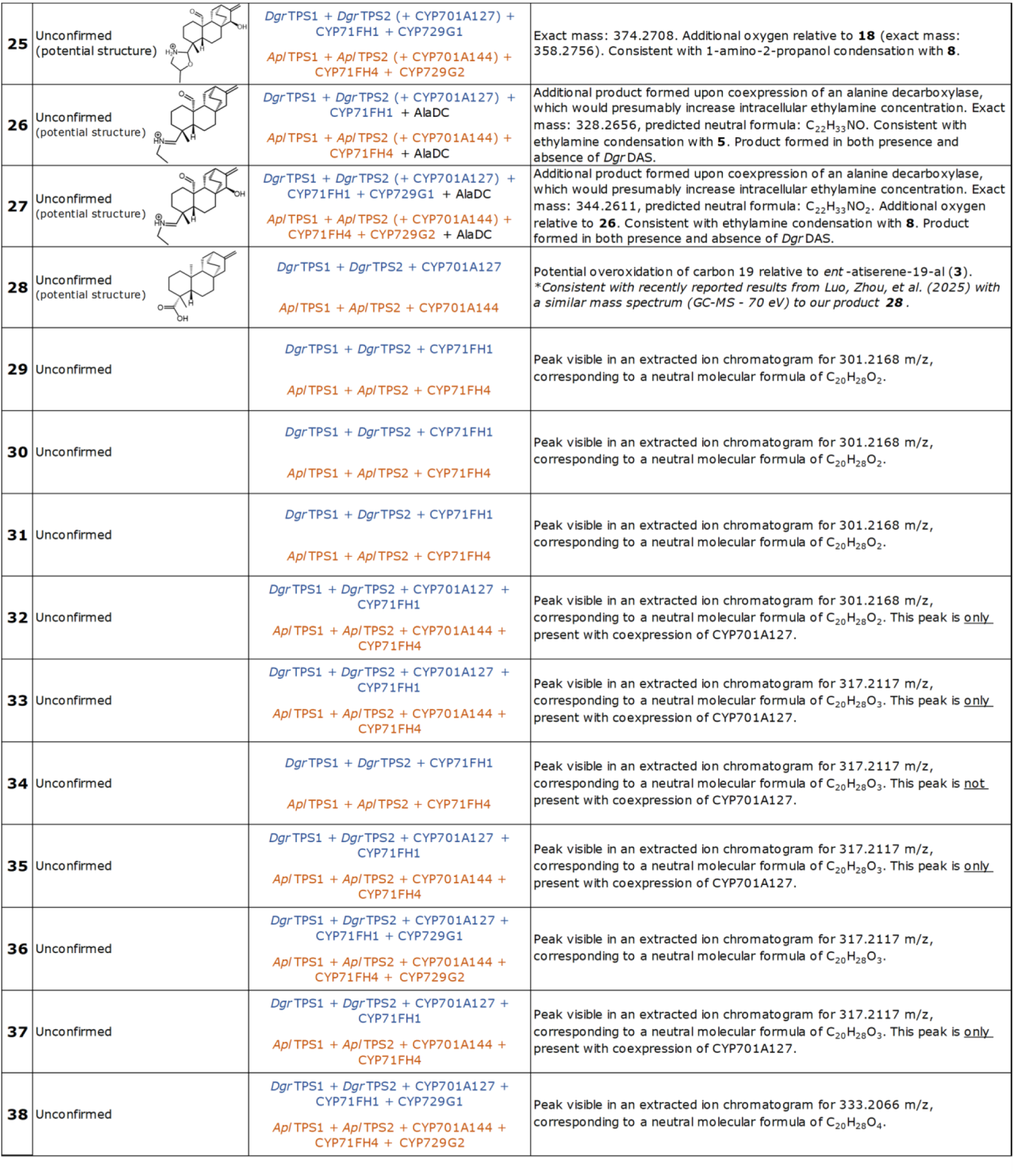
Compound numbering and associated pathway enzymes for remaining numbered compounds referenced in this study.

|  |  |  |  |
| --- | --- | --- | --- |
| **11** | Unconfirmed (potential structure) | *DgrTPS1 + DgrTPS2 + CYP71FH1*<br>*AplTPS1 + AplTPS2 + CYP71FH4* | Overoxidation of ent-atiserene-20-al (**4**) (m/z 302). *Consistent with recently reported results from Luo, Zhou, et al. (2025) with a similar mass spectrum (GC-MS - 70 eV) to our product **11**. |
| **12** | Unconfirmed (potential structure) | *DgrTPS1 + DgrTPS2 + CYP71FH1*<br>*AplTPS1 + AplTPS2 + CYP71FH4* | Overoxidation of ent-atiserene-20-al (**4**) (m/z 300). Two adjacent methyl groups are present on both compounds based on HSQC data on mixture of **4** and **12**. * Structure drawn here was recently proposed by Luo, Zhou, et al. (2025), with a similar mass spectrum (GC-MS - 70 eV) to our product **12**. |
| **13** | Unconfirmed | *DgrTPS1 + DgrTPS2 + CYP701A127*<br>*AplTPS1 + AplTPS2 + CYP701A144* | Exact mass: 806.4192. |
| **14** | Unconfirmed | *DgrTPS1 + DgrTPS2 + CYP701A127*<br>*AplTPS1 + AplTPS2 + CYP701A144* | Exact mass: 644.3676. |
| **15** | Unconfirmed (potential structure) | *DgrTPS1 + DgrTPS2 (+ CYP701A127) + CYP71FH1*<br>*AplTPS1 + AplTPS2 (+ CYP701A144) + CYP71FH4* | Exact mass: 360.2568, predicted neutral formula: C<sub>22</sub>H<sub>33</sub>NO<sub>3</sub>. One additional oxygen relative to **17**. Consistent with difference between compounds **4** and **11**. |
| **16** | Unconfirmed | *DgrTPS1 + DgrTPS2 (+ CYP701A127) + CYP71FH1*<br>*AplTPS1 + AplTPS2 (+ CYP701A144) + CYP71FH4* | Exact mass: 498.2614. Similar fragment ions as **17** in high energy spectra (344.2598 & 326.2496). |
| **17** | Unconfirmed (potential structure) | *DgrTPS1 + DgrTPS2 (+ CYP701A127) + CYP71FH1*<br>*AplTPS1 + AplTPS2 (+ CYP701A144) + CYP71FH4* | Exact mass: 344.2598, predicted neutral formula: C<sub>22</sub>H<sub>33</sub>NO<sub>2</sub>. Consistent with ethanolamine condensation with **5**, and an [M + 4D]<sup>+</sup> peak is observed upon coinfiltration with labeled ethanolamine. Specific structural isomer not determined. |
| **18** | Unconfirmed (potential structure) | *DgrTPS1 + DgrTPS2 (+ CYP701A127) + CYP71FH1*<br>*AplTPS1 + AplTPS2 (+ CYP701A144) + CYP71FH4* | Exact mass: 358.2756, predicted neutral formula: C<sub>23</sub>H<sub>35</sub>NO<sub>2</sub>. Consistent with 1-amino-2-propanol (decarboxylated threonine) condensation with **5**. |
| **19** | Unconfirmed (potential structure) | *DgrTPS1 + DgrTPS2 (+ CYP701A127) + CYP71FH1*<br>*AplTPS1 + AplTPS2 (+ CYP701A144) + CYP71FH4* | Exact mass: 326.2496, predicted neutral formula: C<sub>22</sub>H<sub>31</sub>NO. Consistent with ethanolamine condensation with **5** and an intramolecular condensation. Compound was too low abundance in labeling studies to be directly observed, but corresponding compound **24** shows an [M + 4D]<sup>+</sup> peak. |
| **20** | Unconfirmed | *DgrTPS1 + DgrTPS2 + CYP701A127 + CYP71FH1*<br>*AplTPS1 + AplTPS2 + CYP701A144 + CYP71FH4* | Exact mass: 820.3974. Consistent with the addition of an aldehyde relative to **13** (exact mass: 806.4192). |
| **21** | Unconfirmed | *DgrTPS1 + DgrTPS2 (+ CYP701A127) + CYP71FH1 + CYP729G1*<br>*AplTPS1 + AplTPS2 (+ CYP701A144) + CYP71FH4 + CYP729G2* | Exact mass: 514.2595. Additional oxygen relative to **16** (exact mass: 498.2653). Similar fragment ions as **23** in high energy spectra (360.2530 & 342.2436). |
| **22** | Unconfirmed (potential structure) | *DgrTPS1 + DgrTPS2 (+ CYP701A127) + CYP71FH1 + CYP729G1*<br>*AplTPS1 + AplTPS2 (+ CYP701A144) + CYP71FH4 + CYP729G2* | Exact mass: 376.2481. Additional oxygen relative to **15** (exact mass: 360.2568). |
| **23** | Unconfirmed (potential structure) | *DgrTPS1 + DgrTPS2 (+ CYP701A127) + CYP71FH1 + CYP729G1*<br>*AplTPS1 + AplTPS2 (+ CYP701A144) + CYP71FH4 + CYP729G2* | Exact mass: 360.2530. Additional oxygen relative to **17** (exact mass: 344.2598). Consistent with ethanolamine condensation with **8**, and an [M + 4D]<sup>+</sup> peak is observed upon coinfiltration with labeled ethanolamine. Specific structural isomer not determined. |
| **24** | Unconfirmed (potential structure) | *DgrTPS1 + DgrTPS2 (+ CYP701A127) + CYP71FH1 + CYP729G1*<br>*AplTPS1 + AplTPS2 (+ CYP701A144) + CYP71FH4 + CYP729G2* | Exact mass: 342.2436. Additional oxygen relative to **19** (exact mass: 326.2496). Consistent with ethanolamine condensation with **8** and an intramolecular condensation. An [M + 4D]<sup>+</sup> peak is observed upon coinfiltration with labeled ethanolamine. Specific structural isomer not determined. |

|  |  |  |  |  |
| --- | --- | --- | --- | --- |
| 25 | Unconfirmed<br>(potential structure) |  | *DgrTPS1* + *DgrTPS2* (+ *CYP701A127*) + *CYP71FH1* + *CYP729G1*<br>*AplTPS1* + *AplTPS2* (+ *CYP701A144*) + *CYP71FH4* + *CYP729G2* | Exact mass: 374.2708. Additional oxygen relative to **18** (exact mass: 358.2756). Consistent with 1-amino-2-propanol condensation with **8**. |
| 26 | Unconfirmed<br>(potential structure) |  | *DgrTPS1* + *DgrTPS2* (+ *CYP701A127*) + *CYP71FH1* + *AlaDC*<br>*AplTPS1* + *AplTPS2* (+ *CYP701A144*) + *CYP71FH4* + *AlaDC* | Additional product formed upon coexpression of an alanine decarboxylase, which would presumably increase intracellular ethylamine concentration. Exact mass: 328.2656, predicted neutral formula: C<sub>22</sub>H<sub>33</sub>NO. Consistent with ethylamine condensation with **5**. Product formed in both presence and absence of *DgrDAS*. |
| 27 | Unconfirmed<br>(potential structure) |  | *DgrTPS1* + *DgrTPS2* (+ *CYP701A127*) + *CYP71FH1* + *CYP729G1* + *AlaDC*<br>*AplTPS1* + *AplTPS2* (+ *CYP701A144*) + *CYP71FH4* + *CYP729G2* + *AlaDC* | Additional product formed upon coexpression of an alanine decarboxylase, which would presumably increase intracellular ethylamine concentration. Exact mass: 344.2611, predicted neutral formula: C<sub>22</sub>H<sub>33</sub>NO<sub>2</sub>. Additional oxygen relative to **26**. Consistent with ethylamine condensation with **8**. Product formed in both presence and absence of *DgrDAS*. |
| 28 | Unconfirmed<br>(potential structure) |  | *DgrTPS1* + *DgrTPS2* + *CYP701A127*<br>*AplTPS1* + *AplTPS2* + *CYP701A144* | Potential overoxidation of carbon 19 relative to *ent*-atiserene-19-al (**3**).<br>*Consistent with recently reported results from Luo, Zhou, et al. (2025) with a similar mass spectrum (GC-MS - 70 eV) to our product **28**. |
| 29 | Unconfirmed |  | *DgrTPS1* + *DgrTPS2* + *CYP71FH1*<br>*AplTPS1* + *AplTPS2* + *CYP71FH4* | Peak visible in an extracted ion chromatogram for 301.2168 m/z, corresponding to a neutral molecular formula of C<sub>20</sub>H<sub>28</sub>O<sub>2</sub>. |
| 30 | Unconfirmed |  | *DgrTPS1* + *DgrTPS2* + *CYP71FH1*<br>*AplTPS1* + *AplTPS2* + *CYP71FH4* | Peak visible in an extracted ion chromatogram for 301.2168 m/z, corresponding to a neutral molecular formula of C<sub>20</sub>H<sub>28</sub>O<sub>2</sub>. |
| 31 | Unconfirmed |  | *DgrTPS1* + *DgrTPS2* + *CYP71FH1*<br>*AplTPS1* + *AplTPS2* + *CYP71FH4* | Peak visible in an extracted ion chromatogram for 301.2168 m/z, corresponding to a neutral molecular formula of C<sub>20</sub>H<sub>28</sub>O<sub>2</sub>. |
| 32 | Unconfirmed |  | *DgrTPS1* + *DgrTPS2* + *CYP701A127* + *CYP71FH1*<br>*AplTPS1* + *AplTPS2* + *CYP701A144* + *CYP71FH4* | Peak visible in an extracted ion chromatogram for 301.2168 m/z, corresponding to a neutral molecular formula of C<sub>20</sub>H<sub>28</sub>O<sub>2</sub>. This peak is <u>only</u> present with coexpression of *CYP701A127*. |
| 33 | Unconfirmed |  | *DgrTPS1* + *DgrTPS2* + *CYP701A127* + *CYP71FH1*<br>*AplTPS1* + *AplTPS2* + *CYP701A144* + *CYP71FH4* | Peak visible in an extracted ion chromatogram for 317.2117 m/z, corresponding to a neutral molecular formula of C<sub>20</sub>H<sub>28</sub>O<sub>3</sub>. This peak is <u>only</u> present with coexpression of *CYP701A127*. |
| 34 | Unconfirmed |  | *DgrTPS1* + *DgrTPS2* + *CYP71FH1*<br>*AplTPS1* + *AplTPS2* + *CYP71FH4* | Peak visible in an extracted ion chromatogram for 317.2117 m/z, corresponding to a neutral molecular formula of C<sub>20</sub>H<sub>28</sub>O<sub>3</sub>. This peak is <u>not</u> present with coexpression of *CYP701A127*. |
| 35 | Unconfirmed |  | *DgrTPS1* + *DgrTPS2* + *CYP701A127* + *CYP71FH1*<br>*AplTPS1* + *AplTPS2* + *CYP701A144* + *CYP71FH4* | Peak visible in an extracted ion chromatogram for 317.2117 m/z, corresponding to a neutral molecular formula of C<sub>20</sub>H<sub>28</sub>O<sub>3</sub>. This peak is <u>only</u> present with coexpression of *CYP701A127*. |
| 36 | Unconfirmed |  | *DgrTPS1* + *DgrTPS2* + *CYP701A127* + *CYP71FH1* + *CYP729G1*<br>*AplTPS1* + *AplTPS2* + *CYP701A144* + *CYP71FH4* + *CYP729G2* | Peak visible in an extracted ion chromatogram for 317.2117 m/z, corresponding to a neutral molecular formula of C<sub>20</sub>H<sub>28</sub>O<sub>3</sub>. |
| 37 | Unconfirmed |  | *DgrTPS1* + *DgrTPS2* + *CYP701A127* + *CYP71FH1*<br>*AplTPS1* + *AplTPS2* + *CYP701A144* + *CYP71FH4* | Peak visible in an extracted ion chromatogram for 317.2117 m/z, corresponding to a neutral molecular formula of C<sub>20</sub>H<sub>28</sub>O<sub>3</sub>. This peak is <u>only</u> present with coexpression of *CYP701A127*. |
| 38 | Unconfirmed |  | *DgrTPS1* + *DgrTPS2* + *CYP701A127* + *CYP71FH1* + *CYP729G1*<br>*AplTPS1* + *AplTPS2* + *CYP701A144* + *CYP71FH4* + *CYP729G2* | Peak visible in an extracted ion chromatogram for 333.2066 m/z, corresponding to a neutral molecular formula of C<sub>20</sub>H<sub>28</sub>O<sub>4</sub>. |

